# The compact Casπ (Cas12l) ‘bracelet’ provides a unique structural platform for DNA manipulation

**DOI:** 10.1101/2022.12.16.519550

**Authors:** Ao Sun, Cheng-Ping Li, Zhihang Chen, Shouyue Zhang, Danyuan Li, Yun Yang, Long-Qi Li, Yuqian Zhao, Kaichen Wang, Zhaofu Li, Jinxia Liu, Sitong Liu, Jia Wang, Jun-Jie Gogo Liu

## Abstract

CRISPR-Cas modules serve as the adaptive nucleic-acid immune systems for prokaryotes, and provide versatile tools for nucleic-acid manipulation in various organisms. Here, we discovered a new miniature type V system, CRISPR-Casπ (Cas12l) (∼860 aa), from the environmental metagenome. Complexed with a large guide-RNA (∼170 nt) comprising the tracrRNA and crRNA, Casπ (Cas12l) recognizes a unique 5’ C-rich PAM for DNA cleavage under a broad range of biochemical conditions, and generates gene editing in mammalian cells. Cryo-EM study reveals a ‘bracelet’ architecture of Casπ effector encircling the DNA target at 3.4-Å resolution, substantially different from the canonical ‘two-lobe’ architectures of Cas12 and Cas9 nucleases. And the large guide-RNA serves as a ‘two-arm’ scaffold for effector assembly. Our study expands the knowledge of DNA targeting mechanisms by CRISPR effectors, and offers an efficient but compact platform for DNA manipulations.

## Introduction

The clustered regularly interspaced short palindromic repeats (CRISPR) and CRISPR-associated (Cas) genes function as the adaptive immune module for many prokaryotes and huge phages against invading nucleic acid (Al-Shayeb et al., 2020; Koonin et al., 2017b). Generally, the CRISPR immune response comprises the DNA adaptation, effector biogenesis and nucleic-acid interference stages (Wright et al., 2016). With excellent engineerable capacity, the CRISPR effectors that provide RNA guided DNA targeting and cleaving activities are also effectively repurposed as genomic, epigenomic and transcriptional manipulation tools in many organisms (Barrangou and Doudna, 2016; Knott and Doudna, 2018).

Though an increasing number of CRISPR-Cas effectors have confirmed DNA interference activity *in vitro*, only a few of them, like SpyCas9 and AsCas12a, substantially work and are widely used for efficient genome-editing *in vivo* (Cong et al., 2013; Tsuchida et al., 2022; Zetsche et al., 2015). And among these few effectors, the big molecular size of their Cas nucleases (1200-1400 amino acids(aa)) largely limits the options of delivery vehicles into the target cells. Furthermore, although several types of compact effectors with Cas nucleases <1000 aa have recently been employed for genome editing (CasPhi (Cas12j) effector, 700-800 aa protein monomer with ∼40 nt crRNA; Cas12f effector, 900-1000 aa protein dimer with ∼190 nt sgRNA; CasX (Cas12e) effector, ∼980 aa protein monomer with ∼120 nt sgRNA), the initial versions of these compact systems all exhibit weak or moderate editing efficacy and require extensive and persisted optimization for further application (Harrington et al., 2018; Kim et al., 2022; Liu et al., 2019; Pausch et al., 2020; Tsuchida et al., 2022), similar to how SpyCas9-based technology was developed in the last decade. Moreover, all these compact effectors recognize the T-rich PAM, largely limiting the targeting scope during gene-editing practices. Structural design and directed evolution have been performed to alter the PAM preference for Cas effectors, but the significant decrease of editing efficacy or fidelity has often been observed for those mutants (Kleinstiver et al., 2015; Ma et al., 2019). Therefore, compact but still efficient effectors which offer unique targeting scopes are essential to overcome the application limitations within the current gene-editing tool box.

Here, via a home-developed bioinformatics pipeline using iterative Hidden Markov model (HMM), we identified a new and compact type V CRISPR-Cas family with four orthologous proteins in the environmental metagenome. We termed this new subtype as CRISPR-Casπ, or CRISPR-Cas12l referring to the recent version of complete classification for CRISPR (Makarova et al., 2020). Different from the T-rich PAM preference within the reported type V effectors including those with compact sizes (750-1000 aa protein with 45-190 nt gRNA) (Tong et al., 2020; Wu et al., 2021), the Casπ (Cas12l) effectors (∼860 aa protein with ∼170 nt gRNA) recognize the 5’ C-rich PAM for DNA cleavage under various biochemical environments and exhibit efficient *trans*-activity promising for diagnosis application. Furthermore, even without optimization, the naive versions of Casπ (Cas12l) effectors behave effectively for DNA manipulation both in prokaryotic and eukaryotic cells, maximally reaching ∼50% of the genome-editing efficacy by the well-developed SpyCas9 and LbCas12a effectors. Cryo-EM study revealed that Casπ (Cas12l) protein presents a locked ‘bracelet’ architecture for DNA targeting, which is unique from the canonical ‘two-lobe’ Class 2 nucleases (Cas9 and Cas12). Notably, four non-reported structural domains are identified, including a 69-aa ‘proline-rich string’ loop and a ‘lock-catch’ domain which work together to tie up the Casπ (Cas12l) and lock it around the nucleic-acid target. The large sgRNA composed of the tracrRNA and crRNA folds into a ‘two-arm’ scaffold to recruit and embrace the Casπ (Cas12l) nuclease, forming the stable DNA interference effector. Collectively, our results provide a novel and compact DNA-manipulation platform to substantially expand the CRISPR tool box and offer new aspects to further explore the CRISPR biology.

## Results

### Casπ (Cas12l) is a novel type of compact nuclease guided by a large tracr-crRNA hybrid

During the last decade, huge efforts have been made to explore the CRISPR systems in prokaryotic genome and revealed a large CRISPR kingdom with a deep coverage of functional and structural diversities (Koonin et al., 2017b; Makarova et al., 2020). Nowadays, it is challenging to identify new systems to further expand the CRISPR biology. Therefore, we built an iterative bioinformatic pipeline and performed large-scale environmental sample screening over the land and ocean (Figure S1a). From the metagenome of sludge sample previously collected in Tianjin and Beijing for symbiotic bacteria research, we discovered a new Class 2 CRISPR family with three orthologous systems that bear significant phylogenetic distance from all reported subtypes (Figure 1a; Figure S1b) (Zhao et al., 2019a; Zhao et al., 2018). To reveal the entire CRISPR cassette, the metagenome was re-sequenced and updated (see Materials and methods; NCBI Accession ID: PRJNA857874).

**Figure 1.**
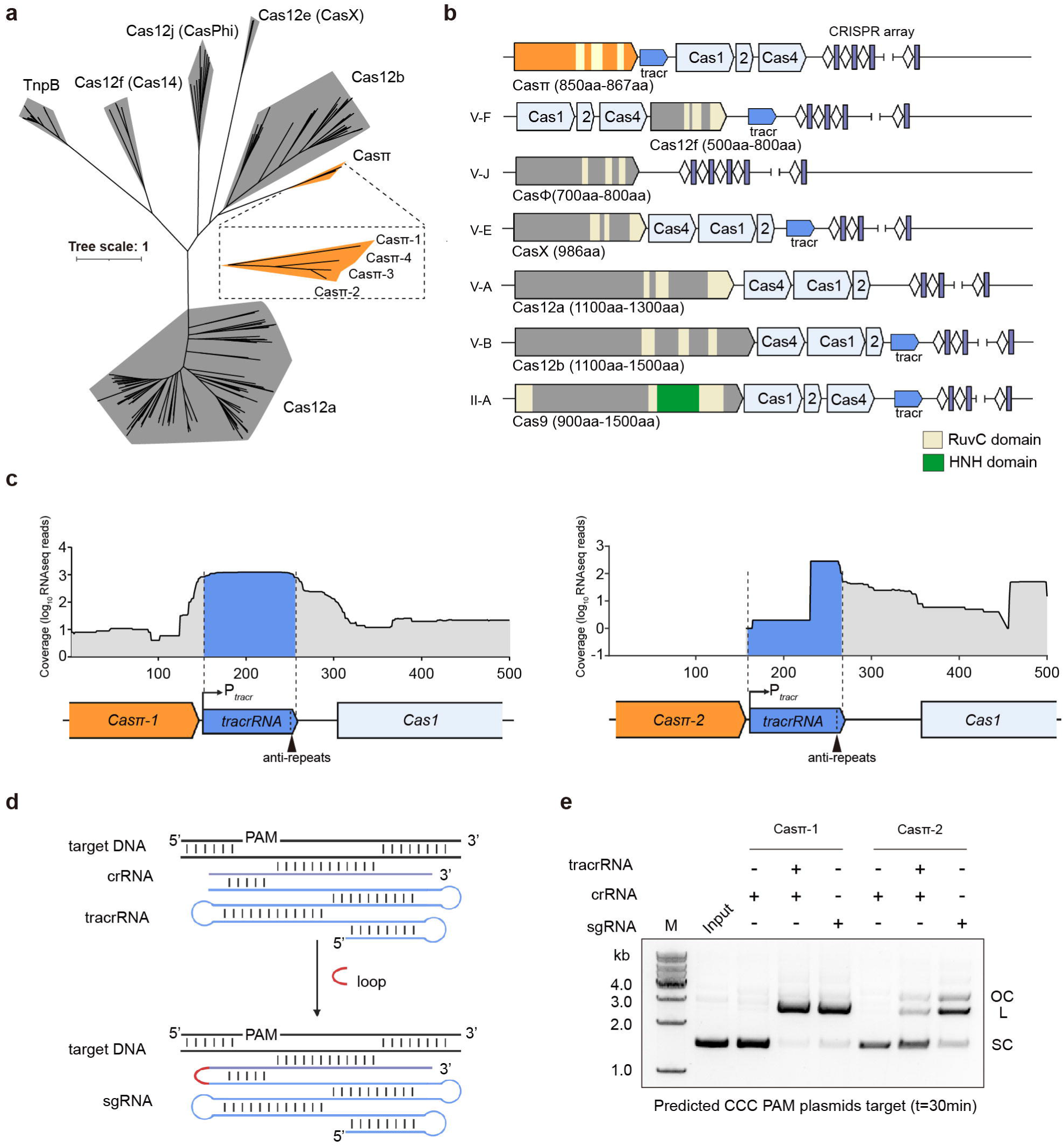
Identification of CRISPR-Casπ. (**a**) Maximum likelihood phylogenic analysis of Casπ orthologs with reported type V Cas nucleases, which are employed in genome-editing application. Bootstrap=1500. (**b**) Genomic loci illustrations of Class 2 CRISPR family members employed in genome-editing application. (**c**) Meta-transcriptome results mapped onto the native genomic loci of Casπ tracrRNA (promoter regions were predicted by BDGP and labeled with P_tracr_; anti-repeat region was labeled with arrow in black). (**d**) Schematic of Casπ dual-guide RNA; crRNA in purple, tracrRNA in cyan. The RNA loop that connects the tracrRNA and crRNA is depicted in red. (**e**) *In vitro* cleavage of plasmids containing predicted CCC PAM by both Casπ proteins using crRNA, tracr-crRNA pair or sgRNA (SC, supercoiled plasmids; L, linearized plasmids; OC, open-circle plasmids)

Overall, this novel system includes the integration module with *cas1*, *cas2* and *cas4* genes, and an uncharacterized gene encoding an 867-aa protein that we refer as Casπ (or Cas12l referring to the recent version of complete classification for CRISPR, hereafter all mentioned as Casπ for convenient description) (Figure 1b; Figure S1c). Via basic local alignment search (BLAST) in public database (Altschul et al., 1990), we further discovered a fourth orthologous system with Casπ-4 (854 aa) which shares about 45% protein sequence identity to Casπ-1 and about 62% identity to both Casπ-2 and Casπ-3 (Figure 1a; Figure S1c, d) (Kantor et al., 2017). Of note, all four CRISPR-Casπ cassettes were validated to reside in the genomes of *Armatimonadetes bacterium* (Figure S1c). Remote homology detection, structural prediction and sequence alignment identified a RuvC nuclease domain near the Casπ C-terminus, with organization reminiscent of that found in type V CRISPR-Cas systems (Figure 1b; Figure S1e) (Gabler et al., 2020; Jumper et al., 2021; Papadopoulos and Agarwala, 2007). The rest of the Casπ protein (504 N-terminal amino acids) showed no detectable similarity to any annotated protein (probability <50% and E-value >200 by HH-suite) (Gabler et al., 2020), suggesting Casπ as a novel type V nuclease. Furthermore, the genomic organization of *cas1-cas2-cas4* integration module in CRISPR-Casπ cassette is unique from the common *cas4-cas1-cas2* pattern within type V systems (Figure 1b). The 37 base-pair (bp) CRISPR-repeats within the four systems share about 68% DNA sequence identity, and the tracrRNA anti-repeat is well identified next to each *casπ* gene rather than proximal to CRISPR repeats as seen in other type V systems (Figure 1c; Figure S1d).

Since the Casπ-1 and Casπ-2 nucleases bear the largest evolution distance within this new family (Figure 1a; Figure S1d), we then chose these two orthologs for further experimental characterization. Via promoter prediction and meta-transcriptome mapping to the anti-repeat regions (see Materials and methods), the tracrRNA sequences for Casπ-1 and Casπ-2 systems were determined and substantially larger >100 nt (Figure 1c; Figure S1c, Tables S1). Further, the DNA cleavage activity of Casπ effectors guided by tracrRNA and crRNA was tested using predicated protospacer adjacent motif (PAM) by CRISPRTarget server (AGC PAM1 for Casπ-1 and CCC PAM2 for Casπ-2) (Biswas et al., 2013). While rarely recognizing PAM1, both Casπ nucleases robustly linearized the target plasmid containing PAM2 using the tracr-crRNA pair or a joint hybrid (single guide RNA, sgRNA) (Figure 1d, e; Figure S1f, g). Thus, Casπ (∼860 aa) associated with a large tracr-crRNA hybrid (∼170 nt) functions as a novel type of compact DNA interference effector.

### Casπ cleaves DNA targets using 5’ C-rich PAM distinct from other Cas12 variants

To further determine the biochemical characteristics of Casπ, we started with identifying the PAM preference of both orthologs using a plasmid library containing five randomized DNA nucleotides upstream of the protospacer (Figure 2a; Figure S2a). Deep sequencing analysis suggests that both Casπ effectors recognize the 5’-CCN-3’ PAM (Figure 2a; Figure S2b, c and Table S1). And specifically, for Casπ-1 effector, the strictness of PAM requirement increases when increasing the salt concentration in the cleavage buffer (Figure S2b). Notably, this C-rich PAM preference for Casπ is different from the T-rich PAM preference for all reported type V nucleases (Figure S2d), which will help to largely expand the targeting scope for type V-based technologies. Using the most favorable CCC PAM determined by plasmid screening assay, we observed efficient cleavage activity for both Casπ effectors on the double-stranded DNA (dsDNA) target even compared to the large *Lachnospiraceae bacterium* Cas12a (LbCas12a, 1228aa) effector (Figure 2b, Table S1). A further screening showed that both Casπ effectors can only robustly cleave the dsDNA target with CCC or CCT (CCY) PAM, indicating a more stringent PAM requirement on dsDNA target (linearized substrate) compared to plasmid target (negative supercoiled substrate) (Figure S2e, f). Gel analysis of the cleavage products from the DNA non-target strand (NTS) and target strand (TS) showed that both effectors generate a staggered cut on the dsDNA (Figure 2c). Consistent with the deep sequencing analysis result for plasmid cleavage (Figure S2a, g and h), the exact cleavage sites locate at 11-14 nt downstream of the PAM on the NTS and 2-4 nt downstream of the protospacer on the TS, thus leaving a 5’ single strand overhang of 6-12 nt on the products (Figure 2d, e). Moreover, we observed the single-stranded DNA (ssDNA) TS cleavage (*cis*-cleavage) by both effectors, and the cleavage efficacy and pattern are comparable to the TS cleavage within dsDNA (Figure S2i).

**Figure 2.**
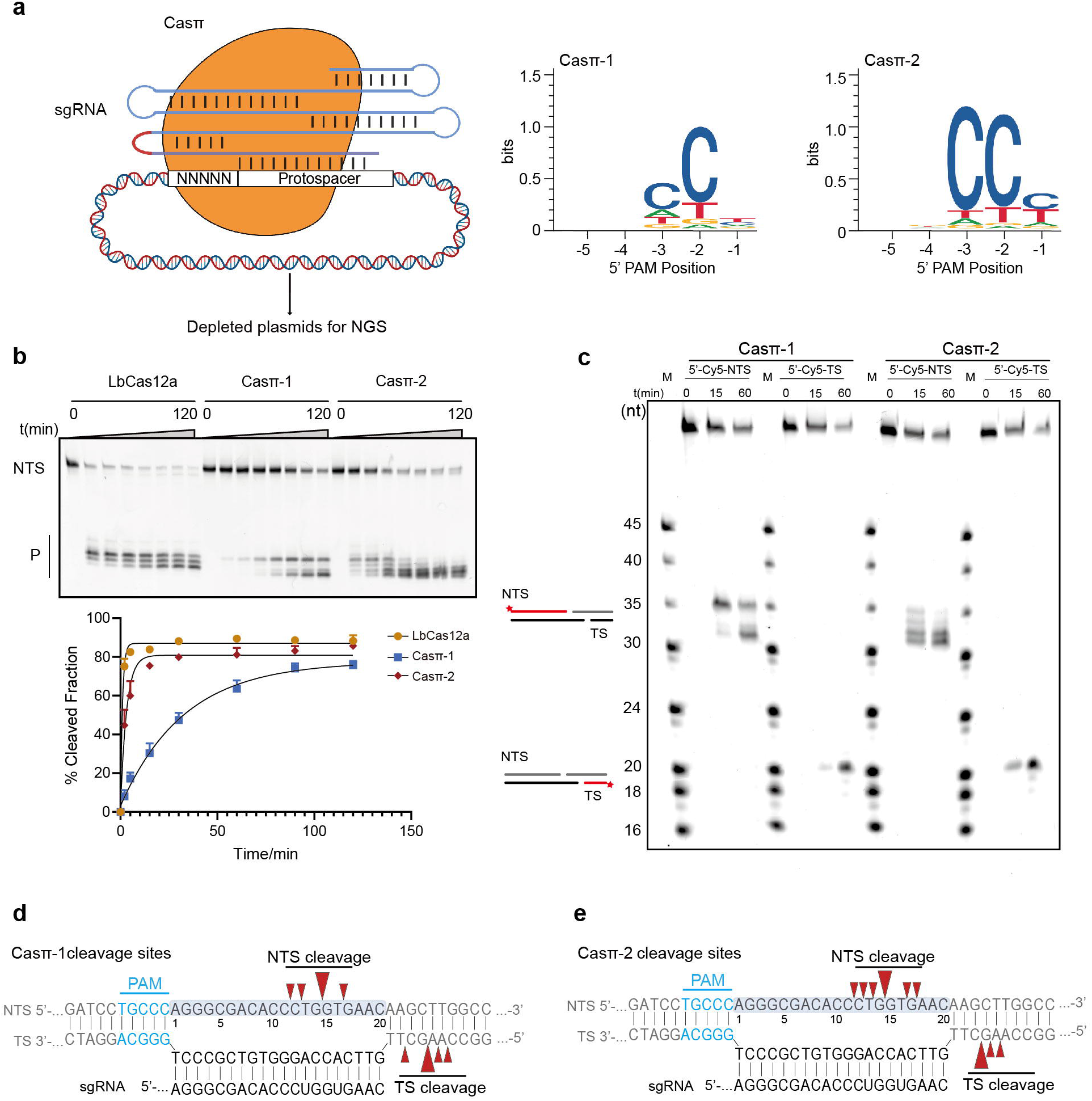
Casπ effector cleaves dsDNA with 5’ C-rich PAM. (**a**) Graphical representation of the *in vitro* PAM depletion assay and the resulting PAMs for both Casπ effectors. (**b**) Top, *in vitro* dsDNA cleavage comparison between LbCas12a, Casπ-1 and Casπ-2 revealed by denaturing PAGE. NTS denotes the non-target strand DNA which is cy5-labeled at 5’ end. P denotes the cleavage products. Bottom, the plot of NTS dsDNA cleavage efficiency by LbCas12a, Casπ-1 and Casπ-2 (n=3 each; means ± SD). (**c**) Cleavage sites mapping of Casπ-1 and Casπ-2 revealed by denaturing PAGE. Lane M shows cy5-labelled marker. (**d** and **e**) The cleavage sites for NTS and TS of Casπ-1 (**d**) and Casπ-2 (**e**) (marked in red arrows, large arrows indicate high probability of cleavage, PAM in blue) suggested both by gel analysis (n=3 each) and NGS analysis.

### Casπ exhibits substantial tolerance of biochemical conditions with efficient *trans*-activity

To explore the application potential of this unique Casπ ‘bracelet’, we performed a general screening for DNA cleavage by both effectors under various biochemical conditions *in vitro*. For RuvC-containing nucleases, divalent ions are typically important to coordinate the catalytical core for DNA hydrolysis. The ion screening suggested that either Mg^2+^ or Mn^2+^ can robustly activate the nuclease activity in Casπ (Figure 3a; Figure S3a). Further experiments also showed that Casπ overcomes several disadvantages reported in other Cas nucleases. Normally, one common drawback of most compact CRISPR-editors (<1000 aa) is their limited tolerance range of salt concentration *in vitro*. For example, the compact AsCas12f and CasPhi (Cas12j) prefer low salt concentration <150 mM NaCl for detectable dsDNA cleavage *in vitro,* due to their limited dsDNA-unwinding ability (Pausch et al., 2020; Wu et al., 2021). Meanwhile, PlmCasX (Cas12e) robustly unwinds the dsDNA for cleavage in high salt concentration condition (300-450 mM NaCl), but gets denatured and precipitated in low salt buffer <300 mM NaCl as seen *in vitro* (Liu et al., 2019). In contrast, the compact Casπ persists a stable effector status for dsDNA cleavage in a wide range of salt concentration from 50 mM to 300 mM NaCl (Figure 3b; Figure S3b). Furthermore, unlike many Cas nucleases which get denatured and precipitated in solution when being concentrated to a high protein-concentration (50-100 μM), both Casπ nucleases behave robustly for physical enrichment (30 kD molecular weight cut-off centrifugal filters; see Materials and methods) (Liu et al., 2019). Therefore, we often stock the Casπ nucleases at the ultra-high protein-concentration of 300 μM for conveniently following uses *in vitro*. Moreover, a huge limitation of employing biomolecular tools in different exogenous scenarios is that they only work efficiently in the temperatures that their source bacterial hosts prefer. To our surprise, although discovered in mesophilic environment, Casπ tolerates temperatures from 25 °C even to 65 °C (Figure 3c; Figure S3c).

**Figure 3.**
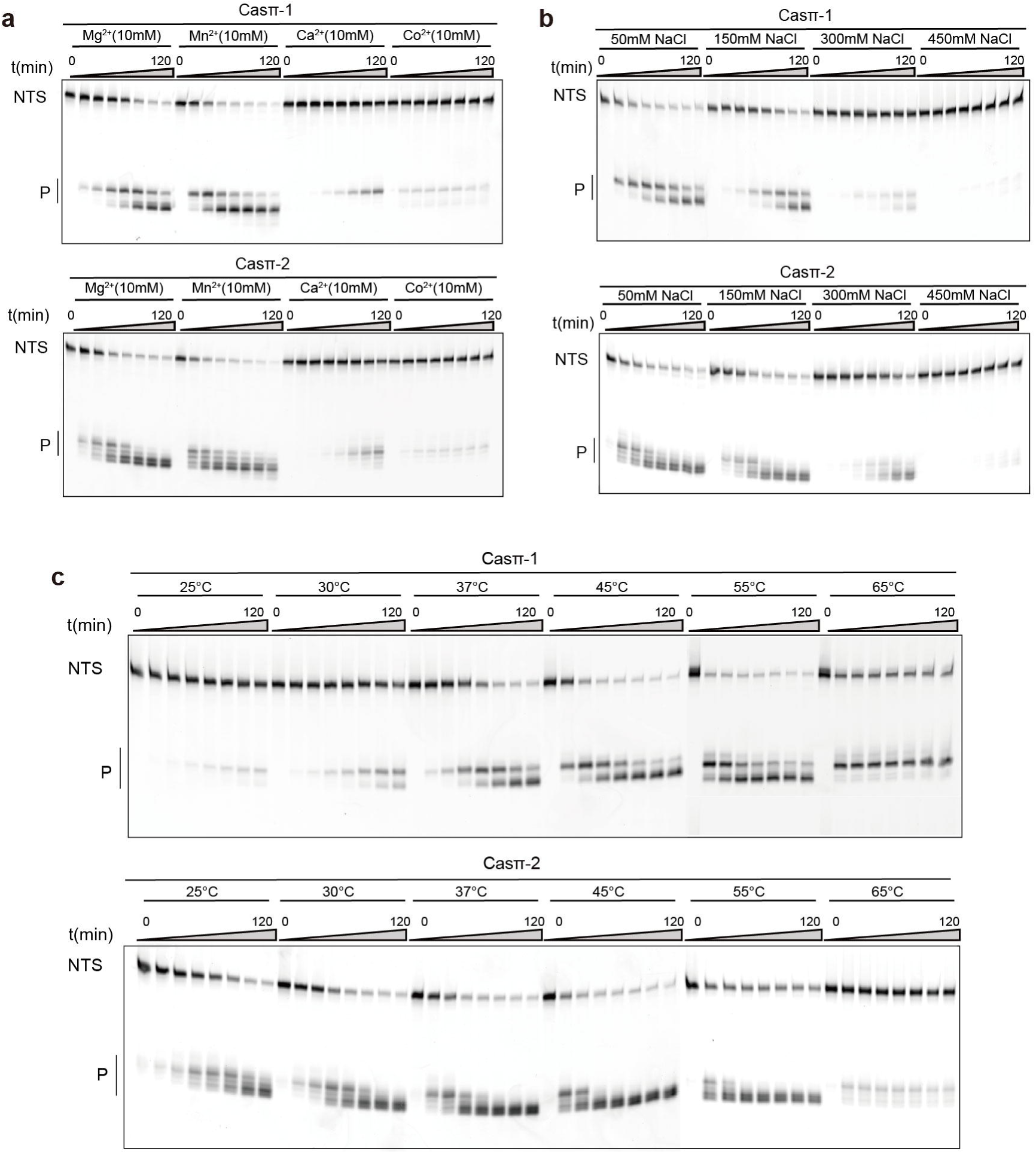
Biochemical screening of Casπ nuclease activity. (**a**) *In vitro* dsDNA cleavage by Casπ effectors using different divalent ions revealed by denaturing PAGE. NTS denotes the non-target strand DNA which is cy5-labeled at 5’ end. Bottom, P means products (n=3 each). (**b**) *In vitro* dsDNA cleavage by Casπ effectors in the buffers with different salt concentrations (n=3 each). (**c**) *In vitro* dsDNA cleavage by Casπ effectors in different temperatures (n=3 each).

To explore the cleavage specificity by Casπ effectors, we first performed the single mismatch screening on the DNA protospacer. The single mismatches between the guide and nucleotides 1-8 of the target DNA at the PAM-proximal region largely abolished the nuclease activity of Casπ (Cas12l), which suggests a “seed region” locating in the position of nucleotides 1-8 of the guide RNA (Figure 4a, b) (Huang et al., 2020; Swarts et al., 2017). Besides, single mismatches between nucleotides 13-16 at the PAM-distal region also significantly decreased the cleavage efficiency of Casπ (Cas12l) (Figure 4a, b). Additionally, many Cas12 nucleases cleave random ssDNA (*trans*-activity) when activated by ssDNA or dsDNA target (activator), which has been harnessed for nucleic-acid diagnosis (Chen et al., 2018; Harrington et al., 2018). Noteworthy, though compact in size, Casπ (Cas12l) effectors show comparable *trans*-activity to the widely used LbCas12a with either ssDNA or dsDNA activator (Figure 4c, d), indicating Casπ’s potential as nucleic-acid diagnosis tool. In summary, compared to many reported Cas effectors, Casπ presents a substantial advantage of flexibility and robustness for exogenous applications.

**Figure 4.**
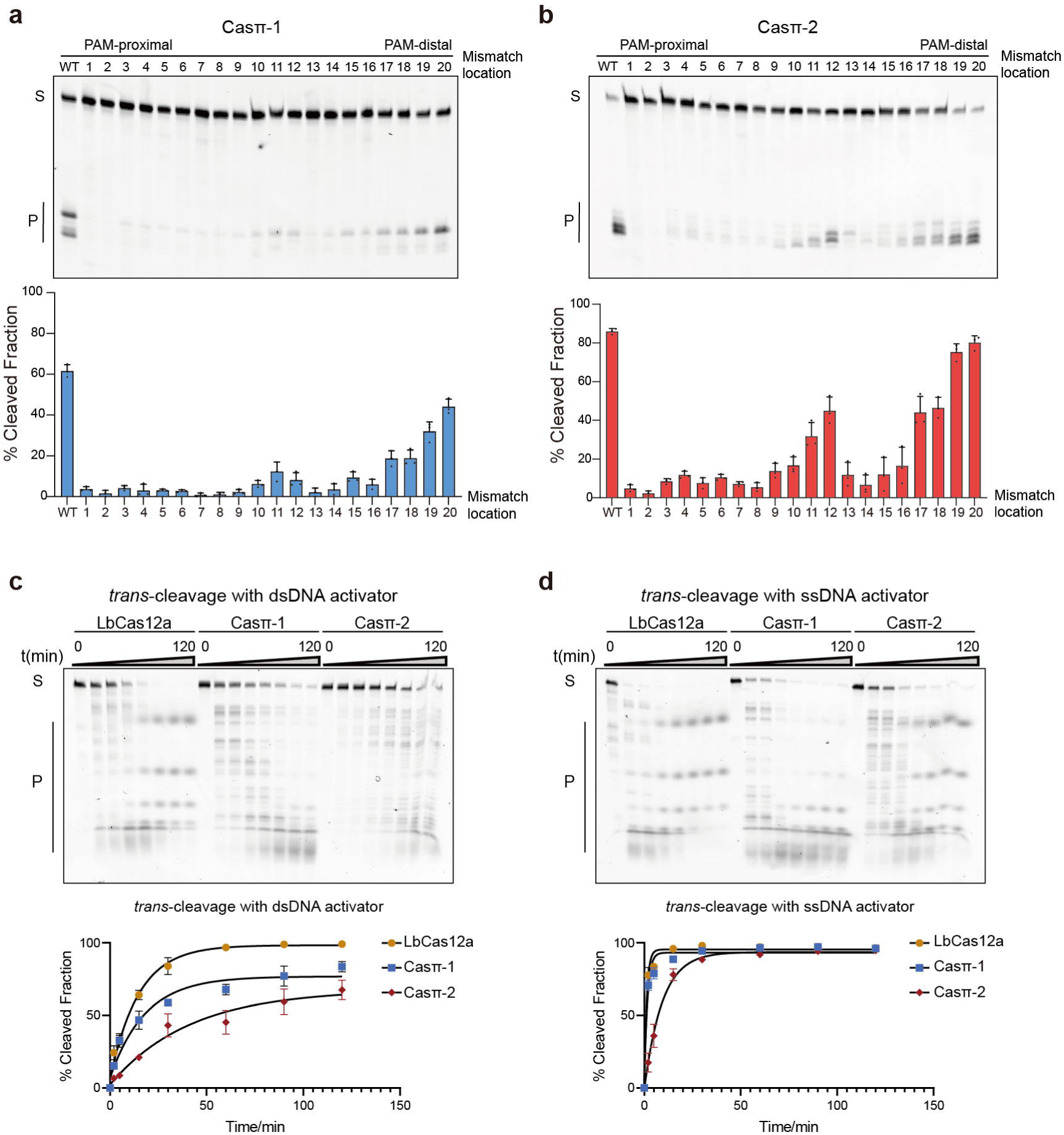
Specificity of DNA cleavage by Casπ. (**a**) Top, cleavage assay using single mismatched target dsDNA by Casπ-1 effector at one hour. Bottom, the bar plot of mismatched-dsDNA substrate cleavage efficiency (WT indicates sgRNA and dsDNA target are fully paired. Numbering means the position of the mismatches between sgRNA and dsDNA target used in this assay; n=3 each; means ± SD). (**b**) Top, cleavage assay using single mismatched target dsDNA by Casπ-2 effector at one hour. Bottom, the bar plot of mismatched-dsDNA substrate cleavage efficiency (n=3 each; means ± SD) (**c**) Top, the *trans*-DNA cleavage by LbCas12a, Casπ-1 and Casπ-2 on ssDNA substrate with dsDNA activators (S means substrates and P means products). Bottom, the plot of *trans*-ssDNA substrate cleavage efficiency by LbCas12a and Casπ-1 and Casπ-2 with dsDNA activator (n=3 each; means ± SD). (**d**) Top, the *trans*-DNA cleavage by LbCas12a, Casπ-1 and Casπ-2 on ssDNA with ssDNA activators (S means substrates and P means products). Bottom, the plot of *trans*-ssDNA substrate cleavage efficiency by LbCas12a and Casπ-1 and Casπ-2 with ssDNA activator (n=3 each; means ± SD).

### Casπ orthologs are active for DNA manipulation both in prokaryotic and eukaryotic cells

To further explore whether the compact Casπ (Cas12l) effectors can be employed for DNA cleavage in prokaryotes, we performed a plasmid interference assay using *E. coli* BW25141 strain carrying a *ccdB* toxin plasmid with arabinose-inducible promoter (Figure 5a). While few survival clones were observed in the non-targeting control due to *ccdB* toxicity, expressing either Casπ-1 or -2 with the *ccdB*-targeting sgRNA yielded significantly more survival clones (Figure 5a, b; Figure S4a, b). This plasmid interference activity was further verified via PCR analysis (Figure S4c).

**Figure 5.**
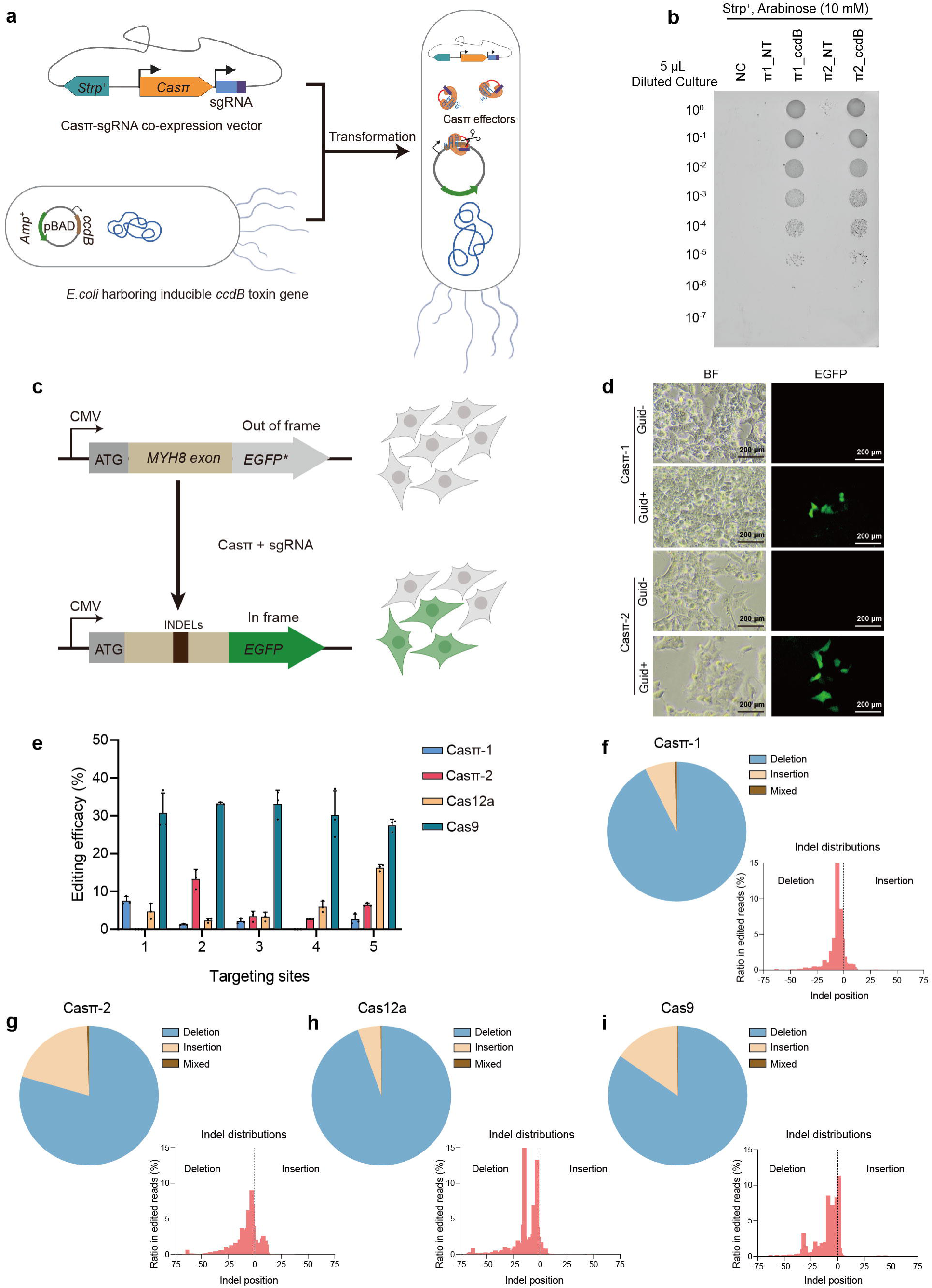
Casπ facilitates DNA manipulation in bacterial and human cells. (**a**) Schematic illustration of the plasmid interference assay. (**b**) Bacteria survival assay on culture plates containing 10 mM arabinose (NT, plasmid with Casπ and non-target sgRNA; ccdB, plasmid with Casπ and sgRNA targeting *ccdB* gene). Dilution gradient is shown in the left. (**c**) Scheme of Casπ mediated *EGFP* lighting-up in human 293A cells. (**d**) *EGFP* lighting-up results. Transfection of plasmids carrying Casπ and sgRNA activated *EGFP* (frame restored) with detectable green fluorescent signal. Both the bright filed (BF) and fluorescent images of cultured cells are shown. (**e**) Editing efficacies determined by NGS from 5 targets mediated by Casπ-1, Casπ-2, Cas12a and Cas9 (n=3 each, mean ± SD). (**f**) Analysis of INDELs generated by Casπ-1 editing within all 15 editing-experiments. Left, pie chart showing percentage of each INDEL within all 15 editing-experiments analyzed by NGS (Mixed means mixed editing with both insertion and deletion). Right, INDEL size distributions within all 15 editing-experiments. (**g**) Analysis of INDELs generated by Casπ-2 editing within all 15 editing-experiments. (**h**) Analysis of INDELs generated by Cas12a editing within all 15 editing-experiments. (**i**) Analysis of INDELs generated by Cas9 editing within all 15 editing-experiments.

Next, to investigate the genome-editing ability of Casπ in eukaryotic cells, we constructed a HEK293A cell line with the genome-integrated ORF containing the *MYH8-*exon and the out-of-frame *EGFP* (Figure 5c; see Materials and methods). Expression of either Casπ-1 or -2 with sgRNA targeting the *MYH-*exon8 efficiently lit up the cells with in-frame EGFP signal, which indicates that the DNA insertions or deletions (INDELs) were generated by Casπ editing (Figure 5d). To compare the editing activity between Casπ effectors and the well-developed LbCas12a and SpyCas9 effector, we designed five parallel targeting sites across the *MYH8*-exon (Figure S4d and Table S3). The edited genomes were PCR amplified, and the editing efficacies were validated by T7E1 assays and quantified by targeted sequencing. (Figure S4e). Next generation-sequencing (NGS) revealed that both Casπ effectors introduced INDELs nearby the cleavage sites in TS as observed *in vitro* (Figure 2d, e; Figure S4f, g). Overall, SpyCas9 presents an average editing efficacy of 30.9% across the five sites and a maximum efficacy of 37.1% at site4 (Figure 5e). LbCas12a shows an average editing efficacy of 6.7% and a maximum efficacy of 16.8% at site5 (Figure 5e). Casπ-1 shows an average editing efficacy of 2.7% and a maximum efficacy of 8.7% at site1 (Figure 5e). Casπ-2 shows an average editing efficacy of 5.4% and a maximum efficacy of 15.6% at site2 (Figure 5e). The combined INDEL analysis on the five targets shows that both SpyCas9, LbCas12a and Casπ effectors mainly generate deletions on the targeted genome (Figure 5f-i; Figure S4h). Of note, SpyCas9 may generate long deletions of ∼40 nt, while Cas12a and Casπ editing dominantly contributes to shorter deletions <25 nt (Figure 5f-i). Further, three more endogenous targets on *B2M* and *TP53* genes were edited by Casπ effectors and the editing efficacies were quantified by NGS (Figure S4i-k).

Therefore, even without any optimizations, the initial versions of Casπ effectors maximumly matched over half of the editing ability of the well-developed SpyCas9 and LbCas12a, supporting Casπ’s potential to be a competitive and compact DNA-manipulation platform with further engineering.

### Unique structural domains in Casπ responsible for DNA interference

To understand the molecular details underlying the DNA-targeting behavior by Casπ effector and provide structural information for editing optimization in future studies, we achieved the cryo-EM map of the R-loop complex containing the deactivated Casπ-1 (D537A, E643A), sgRNA and dsDNA at 3.4-Å resolution (Figure S5a-c, S6a-e). The EM density of Casπ R-loop complex is well resolved which allows us to build the complete atomic model ab initio (Figure 6a-c; Figure S6e, f and Video S1). Consistent with the primary sequence BLAST suggesting no significant similarity to reported proteins, Casπ also exhibits a unique 3D architecture compared to other CRISPR-Cas nucleases revealed by structural alignment with Dali server (Figure S7a, b).(Holm and Rosenström, 2010) Only moderate similarity was observed between Casπ and Cas12 nucleases, mainly within the RuvC domain and oligonucleotide binding domain (OBD) (Figure S7c, d). Then, referring to CasX (Cas12e) which shares the top structural similarity to Casπ and also uses a large RNA guide (Figure S7e), we further located the conserved bridge helix (BH) element and four unique structural domains within Casπ, including the lock-catch (LC) domain, proline-rich string (PRS), Helical-I domain and NTSB (non-target strand binding domain) chimera (HNC), and Casπ (Pi) C-terminal (PCT) domain (Figure 6a-c; Video S1).

**Figure 6.**
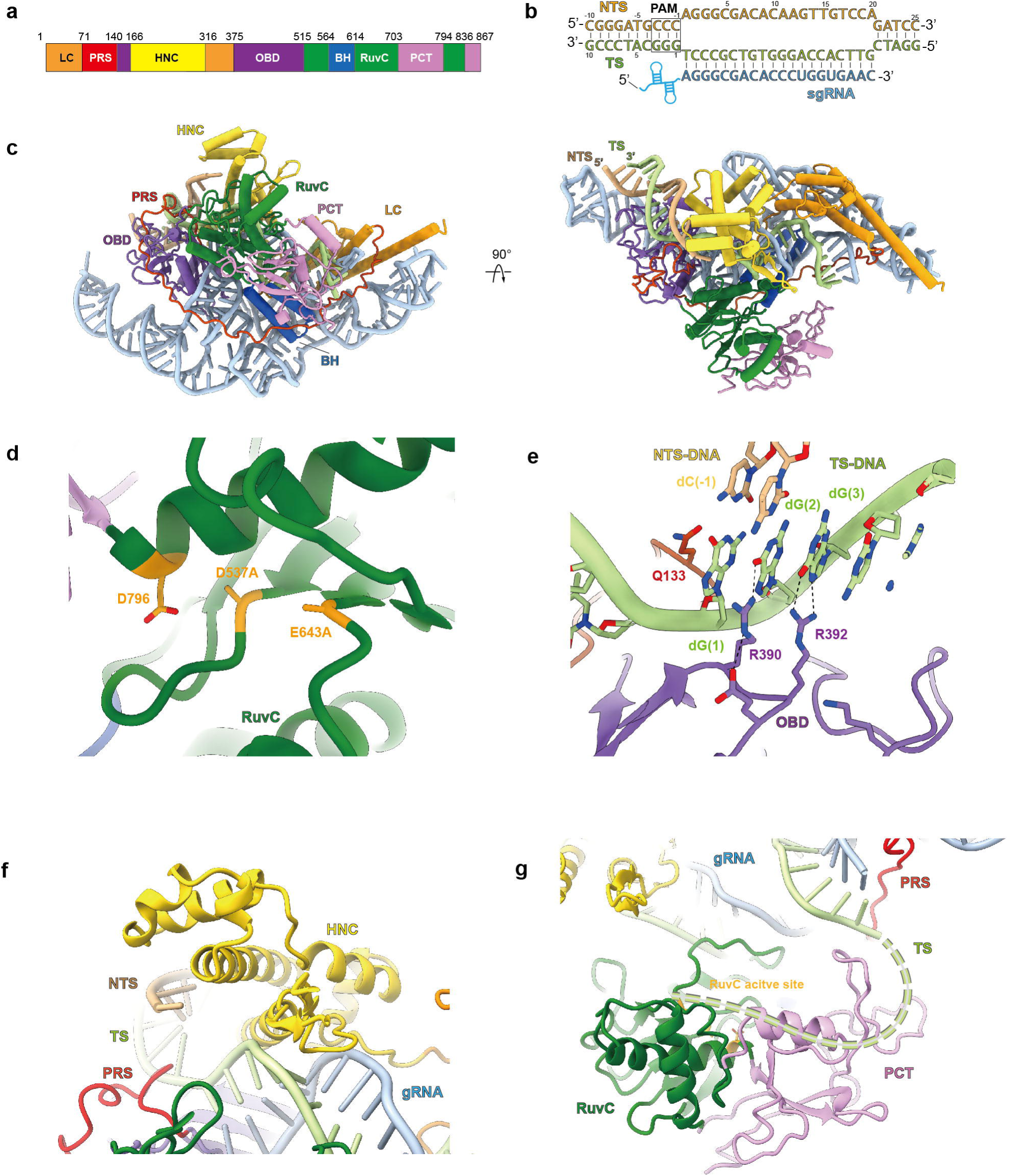
The structure of Casπ nuclease. (**a**) The domain organization aligned with primary sequence. LC domain is colored in dark orange, PRS in red, HNC in yellow, OBD in purple, RuvC in dark green, BH in dark blue, and PCT in pink. (**b**) The base-paring details for the R-loop region. The sequences for NTS (light orange color), TS (light green color) and sgRNA spacer (cyan color) are presented. The PAM region is marked with rectangle. (**c**) The atomic model for Casπ R-loop complex. The protein domains, DNA and sgRNA are colored referring to panel a and b. The front and top views are presented. (**d**) The structural details within Casπ RuvC domain (dark green color). The three catalytic residues were highlighted with dark yellow color. And in this complex, D537 and E643 were mutated to alanine. (**e**) The molecular details for PAM recognition. The amino acids involving for dG(2) and dG(3) recognition are labeled. The key hydrogen bonds are shown by dashed lines. (**f**) The structural details within Casπ HNC domain (yellow color). (**g**) The structure of Casπ PCT domain (pink color) and TS-DNA loading model. The 5’ end of TS DNA is hypothetically modeled as dashed line (light green) and loaded into RuvC nuclease pocket by PCT domain.

The RuvC domain in Casπ displays a canonical DNA-cleavage pocket with the conserved triplet of catalytical residues D537, E643 and D796 (Figure 6d). D537 and E643 are mutated to alanine in this study for stabilizing the complex (Figure S5a-c). Different from other type V CRISPR nucleases which prefer T-rich PAM, two unique residues in Casπ OBD domain, Arg390 and Arg392, were observed to recognize the two guanine nucleotides (dG(2) and dG(3) in the TS) complementary to the CCN PAM (in the NTS) (Figure 6b, e). Both the single mutations (R390A or R392A) and double mutation (R390A/R392A) totally abrogated the nuclease activity of Casπ (Figure 6e; Figure S8a, b). In addition, the side chain of Gln133 inserts into the downstream site of PAM duplex, which may generate local dsDNA melting for sgRNA-spacer invading as discussed in other type V nucleases (Figure 6e) (Swarts et al., 2017).

The HNC domain, which presents as a structural chimera of Helical-I domain and NTSB domain in CasX, interacts both with the seed region of sgRNA-DNA heteroduplex at the PAM-proximal region and also the backbone of DNA non-target strand to stabilize the R-loop conformation (Figure 6f; Figure S8c). Meanwhile, neither primary sequence BLAST or structural search for PCT domain (Trp703 to Asp794 and Arg836 to Ile867) reveals any suggestive similarity to annotated proteins, indicating that this unique feature is specific to Casπ nucleases (Figure S7b). Since the PCT domain sits at similar primary and spatial locations as the target-strand loading (TSL) domain of CasX (Figure S8d), we then hypothesize that the PCT domain may help with the target strand loading into RuvC nuclease domain (Figure 6g), and this needs to be further explored in future studies (Liu et al., 2019).

### Casπ presents a ‘bracelet’ architecture encircling the nucleic-acid target

Strikingly, a long ‘proline-rich string’ (PRS) loop composed of 69 amino acids (Pro72 to Trp140) is largely resolved in the EM map (Figure 7a; Figure S6f and Video S1). There are 14 prolines and 17 charged residues within this ‘string’ which makes it adopt high structural accessibility and electrostatic capacity to tie up the whole complex via multi-interactions with other protein domains, sgRNA and also the DNA target (Figure 7a; Figure S9a). Directly next to the PRS N-terminus, Casπ folds into a two-helix structure (Met1-Asp71) which serves as a ‘lock’ and tightly interact with a three-helix ‘catch’ module (Val317-Ala375) through multiple interactions, such as the hydrogen bonds (E28 and Y61 interact with R339 and E337, respectively) (Figure 7b), the charged interactions and the van der Waals interactions (not shown in the panel). Via this unique structure never observed in other Cas nucleases, the ‘lock-catch’ (LC) domain further locks the ‘tie-up’ conformation mediated by the PRS (Figure 7a; Figure S9a and Video S1). Moreover, the ‘lock’ part in LC domain intensively interacts with the sgRNA stem to stabilize the assembly of R-loop complex (Figure 7b; more details discussed in next section). Remarkably different from the canonical ‘two-lobe’ architecture for Class 2 Cas nucleases, the PRS and LC domains string all other protein domains together, and make the Casπ look as a locked ‘bracelet’ encircling the nucleic-acid target (Figure 7c, d; Figure S9b, c).

**Figure 7.**
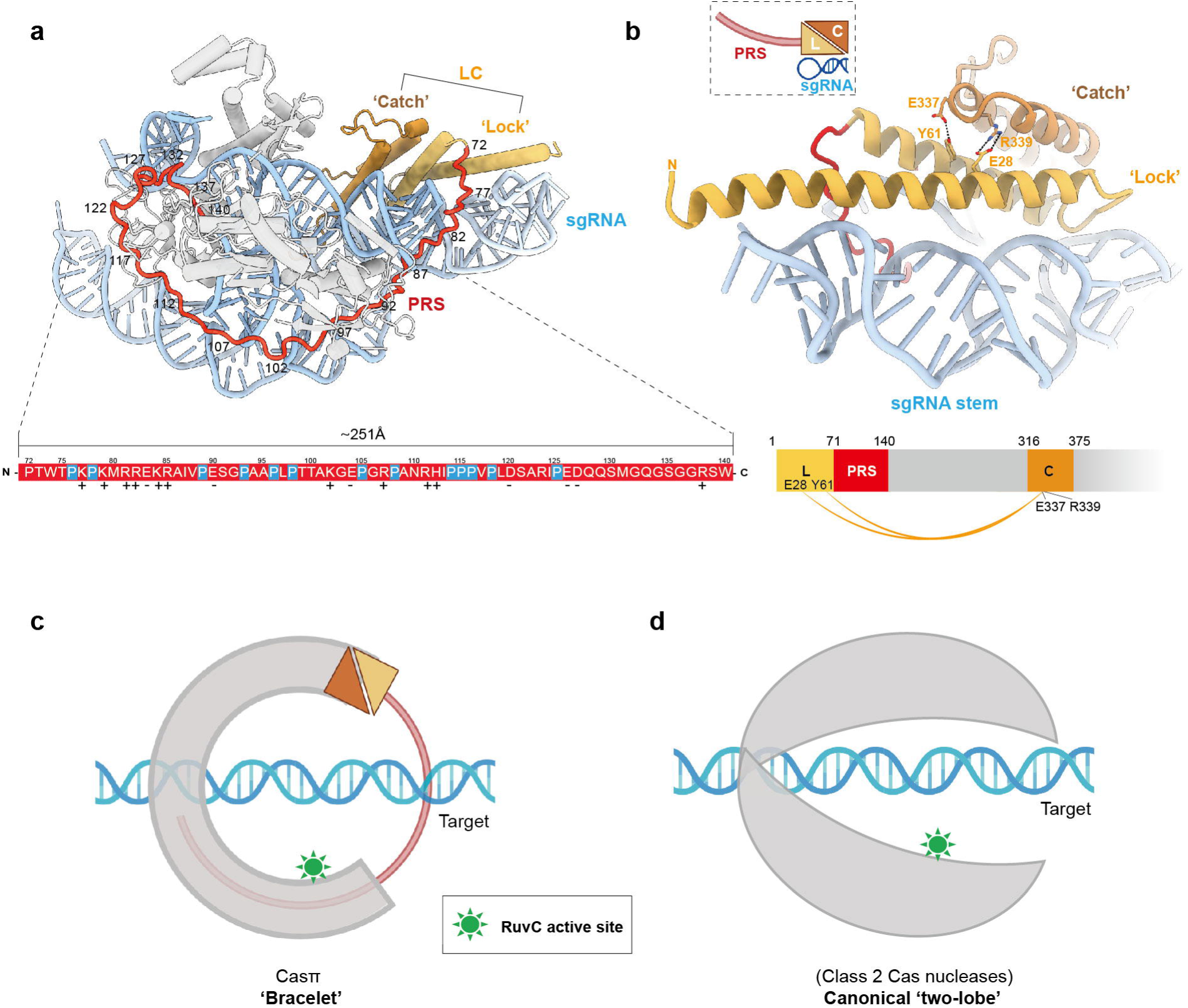
The structure of ‘bracelet’ architecture of Casπ. (**a**) The structural distribution of PRS domain across the Casπ R-loop complex. For clear presentation, the sgRNA and DNA are shown in cyan. ‘Lock’ part in LC domain is shown in light orange and the ‘catch’ part in deep orange. PRS is shown in red and all other protein domains in grey. The primary sequence for PRS loop is colored in red and shown in the bottom panel. The prolines in PRS are specially depicted in blue. The positively charged amino acids are labeled with ‘+’ and negatively charged ones with ‘-’. (**b**) The structural details of LC domain. The elements are colored referring to panel A. Side chains of the amino acids involved in the interactions between ‘lock’ and ‘catch’ are shown, and the formed hydrogen bonds are presented as dashed-lines. The charged interactions and van der Waals interactions are not shown in the panel. The cartoon shapes for the ‘lock (L)’, ‘catch (C)’, PRS and sgRNA stem are outlined and presented at the top left. The domain organization of ‘lock (L)’, ‘catch (C)’ and PRS is presented at the bottom. The interactions between ‘lock (L)’ and ‘catch (C)’ are connected with orange curves. (**c**) A cartoon model for Casπ ‘bracelet’. The PRS are modeled as a half-ring colored in red. The LC are modeled referring to panel b, and all other domains are modeled as a half-ring colored in grey. The sgRNA stem and the nucleic-acid target are also modeled and labeled, accordingly. The RuvC active sites are indicated in green color. (**d**) The cartoon model of two-lobe architecture for canonical Class 2 Cas nuclease. The protein part was colored in grey and nucleic acid part in cyan. The RuvC active sites are indicated in green color.

### The large tracr-crRNA hybrid forms a ‘two-arm’ scaffold for effector assembly

The compact Casπ uses a large guide-RNA (tracr-crRNA hybrid) for DNA interference. Well-resolved in the cryo-EM map (Figure S6), the sgRNA hybrid presents as a ‘two-arm’ architecture and embraces the Casπ monomer forming the ribonucleoprotein (RNP) effector (Figure 8a). Referring both to the 2D and 3D structural details, we located four structural elements within this large sgRNA scaffold: arm-I (A-I), junction region (JR), arm-II (A-II) and pseudoknot region (PR) (Figure 8a, b). Both A-I and A-II are built by the three-way junction, and these two three-way junctions are connected by the junction region (JR). While A-I (previously labeled as ‘sgRNA stem’ in Figure 7b) forms intensive interactions with Casπ protein (Figure 8c, d), A-II largely stretches out from the effector complex (Figure 8a, b). Noteworthy, both 12nt and 24nt truncation on the A-II increased the DNA cleavage activity by Casπ, suggesting a promising engineering site on sgRNA for improving the genome-editing capability (Figure S10a, b). As well, this stretched A-II may provide a flexible engineering site for functional module integration without affecting the Casπ effector assembly. In addition, beyond the electrostatic interactions with RNA backbone (Figure 8c), the binding between Casπ and sgRNA is also developed in a sequence specific way. For example, the bases of nucleotides C48 and G49 in A-I was respectively recognized by Arg23 and Arg26 residues in the LC domain (Figure 8c, d). Moreover, the U_148_GAAAG_153_ in crRNA part pairs with the C_100_UUUCA_105_ loop from the tracrRNA part, forming a pseudoknot structure (corresponding to the PR) followed by the single-stranded spacer (Figure 8a, b). This PR element tightly binds to Casπ PRS, BH, RuvC and OBD domains via backbone interactions and base-specific recognitions (Figure 8c, e and f). Noteworthily, the sgRNA PR also gets shielded by the Casπ PRS domain (Figure 8e). In summary, majorly mediated by the A-I and PR elements, the sgRNA hybrid provides a structurally continuous ‘two-arm’ scaffold to recruit the Casπ ‘bracelet’ via both backbone interactions and base-specific recognitions, forming a compact and ‘locked’ effector for DNA interference (Figure 8; Video S1).

**Figure 8.**
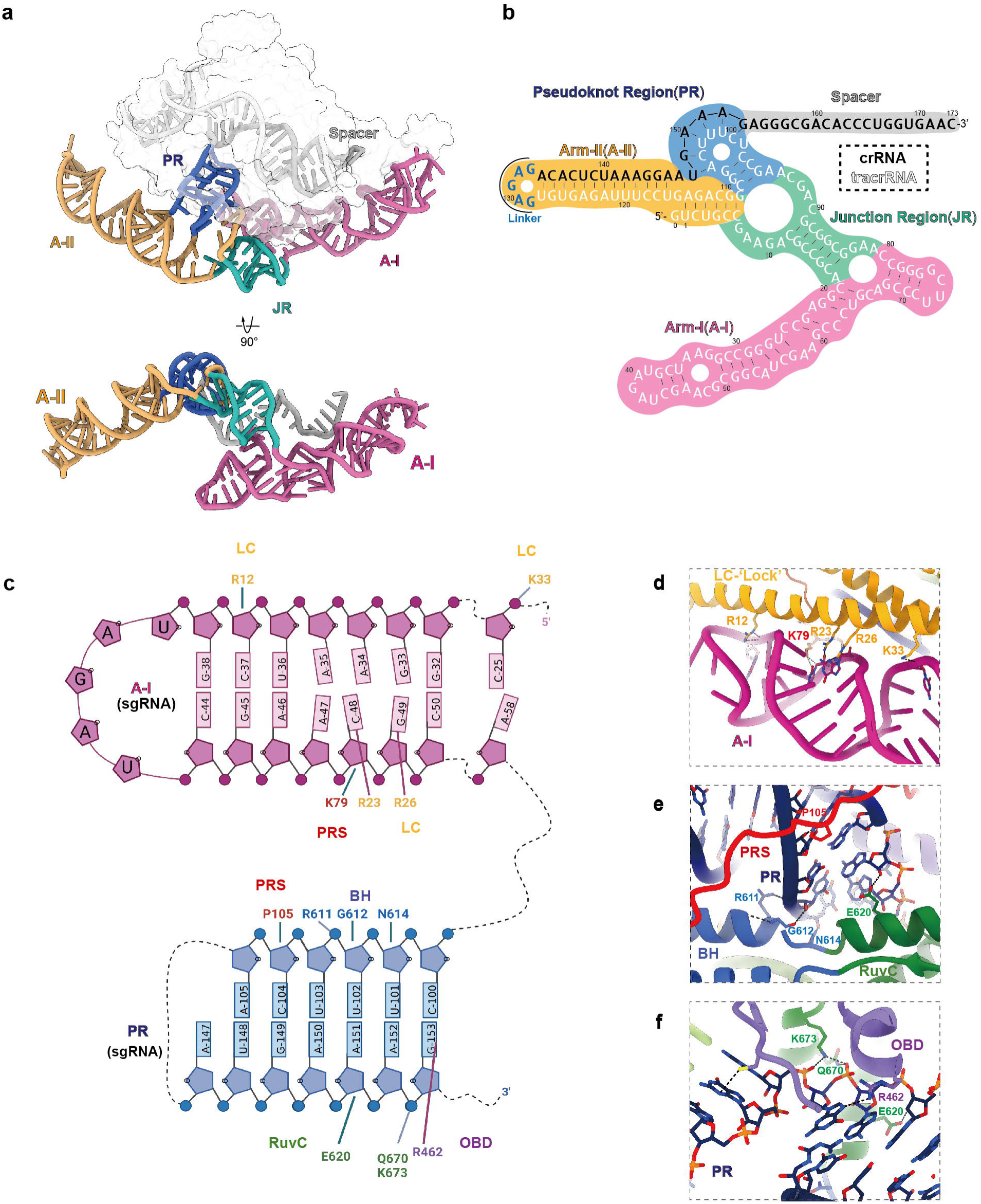
The structure of Casπ sgRNA. (**a**) The overall 3D structure of sgRNA. The A-I region is colored in plum, JR in green, A-II in orange, PR in blue and spacer in grey. Both front and bottom views are shown. The protein density is shown by transparent surface in the top panel. (**b**) The secondary structure details for the sgRNA. The background of different regions is colored according to panel A. The sequences for tracrRNA part, joint-loop and crRNA part are shown in white, blue and black, respectively. (**c**) The interaction details between Casπ protein and the sgRNA. Only the sgRNA A-I and PR regions are shown in this cartoon. The protein domains, associated amino acids and RNA nucleotides are labeled. The interaction pairs are linked with solid lines. (**d**) The structural details for the interaction interface between LC domain and A-I element. (**e** and **f**) The interaction details between PR element and PRS, BH (**e**), RuvC and OBD (**f**) domains of Casπ. The protein domains are colored and labeled referring to Figure 6a.

## Discussion

### Casπ provides a unique DNA-targeting platform with a large potential given further engineering

In this study, via large-scale bioinformatic screening and manual annotation, we identified the CRISPR-Casπ as a novel type V system distant from reported families which provides unique potentials for gene editing application, like the C-rich PAM preference, compact size, tolerance of various biochemical conditions and efficient *trans*-activity. Significantly, without any optimization, the naive version of Casπ effectors (∼860 aa) maximally matches over half of the editing ability by the well-developed SpyCas9 and LbCas12a. Then, this strongly suggests that Casπ has a huge potential to be largely improved via rational design or directed evolution, similar to how SpyCas9 or other effector-based technologies were developed in the last decade. Meanwhile, our cryo-EM study revealed the ‘bracelet’ architecture for Casπ which provides a brand-new structural platform for function-module integration and engineering. Furthermore, given the well-illustrated recognition details by Casπ protein, the ‘two-arm’ sgRNA also offers large engineering capacity, especially within the stretched-out A-II element.

### Strictness for PAM preference varies in different scenarios

PAM sequence is essential for dsDNA targeting by Class 2 Cas nuclease, and it is often determined by the cleavage of plasmid library containing randomized PAM either *in vitro* or *in vivo*. In our experience, Cas effectors usually show more robust cleavage on the plasmid target than linearized dsDNA (Tsuchida et al., 2022), as plasmids contain melting bubbles in the supercoil conformation(Adamcik et al., 2012). And compared to the plasmid, a more stringent PAM requirement was observed on the linearized dsDNA target (Figure S2e, f). Moreover, we also found that the dC gradually dominated the third position of the PAM in the depletion analysis for Casπ-1 while increasing the salt concentration in the cleavage buffer, which indicates a more stringent PAM preference for Casπ-1 effectors in high salt buffer (Figure S2b). Similar patterns were observed in CasX enzymes (non-published data). Referring to previous biophysical studies, either linearizing the plasmid (relax the supercoil and re-anneal the bubbled strands in plasmids) or increasing the salt concentration (stabilize the dsDNA conformation) may contribute to ‘tougher’ targets for Cas effectors to unwind (Adamcik et al., 2012). Therefore, we would suggest that a stringent PAM sequence determined with the ‘tough’ condition (linearized dsDNA target in buffer with the highest salt concentration that Cas effectors can tolerate) may be the prioritized choice for gene-editing application.

### A hypothetical evolution trend underlying Class 2 CRISPR effectors starting from the ‘RNA world’

The wet-lab validation and structural information allow us to accurately identify the functional size of each component in Cas effectors, especially for the tracrRNA whose exact length are usually challenging to determine bioinformatically. When arranging the structurally validated Class 2 effectors (using tracr-crRNA guide) together with our newly discovered Casπ effector, an interesting trend was observed: the size of tracr-crRNA hybrid (RNA part) gradually decreases as the Cas protein size increases within the RNP effectors (Figure S11a-d). Moreover, analysis of 383 bioinformatically identified Cas9 effectors also suggest a negative linear correlation (correlation coefficient of -0.439) between the sizes of the tracr-crRNAs and Cas proteins (Figure. S11e. Considering that the linear correlation is sensitive to extreme values, we only selected the effectors with Cas9’s molecular weight between 100,000-200,000 Da and tracr-crRNA between 30,000-60,000 Da for analysis). Notably, a recent structural study shows that the IscB effector (commonly-acknowledged ancestor for type II Cas9 effectors) comprises an IscB nuclease monomer smaller than reported Cas9s and the ωRNA significantly larger than reported tracr-crRNA hybrids (Figure S11a).(Schuler et al., 2022)

Then starting from the IscB or other ancestors like TnpB for type V effectors,(Altae-Tran et al., 2021; Karvelis et al., 2021; Koonin et al., 2017a) this trend may suggest an RNA-protein co-evolution path underlying the CRISPR effectors (Figure S11a, b) (Cech, 2009; Poole et al., 1998). As proteins play more robust structural and enzymatic roles than RNAs, during the molecular evolution, the functional and structural size of the RNA part are gradually replaced by Cas protein for efficient DNA interference (Figure S11a, b). And this has actually often been the case that the CRISPR effectors with large Cas proteins and small gRNAs work better for DNA editing than the effector with small Cas protein and large gRNA.(Li et al., 2020; Liu et al., 2019; Pausch et al., 2020)

Further, even ancestral to the IscB or TnpB ‘intermediate’ ancestors, it is also reasonable to hypothesize the RNA and RNA-dominated ancestors for CRISPR effectors, in which the RNA part (ribozymes) but not the protein may play the enzymatic role for nucleic acid interference (Figure S11f).(Chylinski et al., 2013; Chylinski et al., 2014; Joyce and Szostak, 2018; Poole et al., 1998; Robertson and Joyce, 2012; Wilson and Szostak, 1999) Though probably not exist in the current protein dominated world, reconstruction of those RNA and RNA-dominated ancestors originated from the ‘RNA world’ will provide brand-new insights for molecular tool development, as well as the evolutionary evidence of enzymatic function transition from RNA to protein. While due to lacking available knowledge, our current discussion is only focused on the molecular size of a limited number of CRISPR effectors. Thereby, a large-scale identification of new CRISPR effectors in the current protein dominated world and a comprehensive understanding of the functional and structural replacement events between the RNA and protein may help to understand the ‘co-evolutionary principle’ starting from the ‘RNA world’ (Figure S11f). Using this ‘co-evolutionary principle’, it’s promising to reconstitute those RNA and RNA-dominated ancestors *in silico*.

## Materials and Methods

### Metagenomics

The genetic materials were purified from bioreactor sludge sample as previously described, and sequenced on the Illumina NovaSeq 6000 platform using the PE150 sequencing strategy (Zhao et al., 2019b). All raw datasets were trimmed by Trim Galore v0.6.5 using default parameters, which generated data containing clean reads that were subsequently assembled using SPAdes v3.15.4 for detection of CRISPR-Cas system (Nurk et al., 2013).

### Casπ detection and phylogenic analysis of type V CRISPR systems

The assembled contigs were scanned for Cas nucleases using hidden Markov model (HMM) profiles, which were built using the HMMER (Potter et al., 2018), based on Cas nuclease sequence alignments from Clustal Omega (1.2.4) (Sievers et al., 2011). CRISPR arrays were identified using local version of the CRISPRCasFinder (4.2.20) and CRISPRidentify (v1.1.0) (Couvin et al., 2018; Mitrofanov et al., 2021). Loci that contained both *cas1* and the CRISPR array were further analyzed to identify the proteins located within the range from 20,000 nucleotides upstream to 20,000 nucleotides downstream of the CRISPR array. Potential functions of these proteins were annotated by HMMs and the local version of eggNOG mapper (2.1.6, eggNOG DB version: 5.0.2, MMseqs2 version: 13.45111) (Cantalapiedra et al., 2021; Huerta-Cepas et al., 2019). Proteins larger than 600 amino acids were selected as potential Class 2 Cas nucleases with nucleic-acid interference activity, and were further clustered by phylogenetic analysis.

For phylogenetic analysis, sequences of reported Cas nucleases were collected from UniProt database by searching key words of each nucleases, like Cas9 and Cas12a(UniProt: the universal protein knowledgebase in 2021, 2021; Burstein et al., 2017; Harrington et al., 2018; Makarova et al., 2020; Pausch et al., 2020). Sequence alignment of Casπ with the selected type V Cas nucleases was generated using Clustal Omega (1.2.4) (Sievers et al., 2011). Phylogenic reconstruction was performed using IQ-TREE2 (2.0.7) with VT+F+R7 as the substitution model and 1500 bootstrap sampling (Minh et al., 2020). Reconstruction result was visualized and edited using iTOL v6.5.8(Letunic and Bork, 2021).

### Protein sequence and CRISPR repeat analysis

The protein and CRISPR repeat sequences of four Casπ orthologs were analyzed by Clustal Omega server with default parameters (Sievers et al., 2011), and the two heatmaps illustrating the sequence similarity were built using the similarity score matrix. For protein alignment with other type V CRISPR, the protein sequences of four Casπ orthologs were aligned with LbCas12a, AsCas12a, AaCas12b and DpbCas12e proteins using NCBI COBALT program (Papadopoulos and Agarwala, 2007), and the key amino acids in RuvC domains of Casπ were inferred from the alignment results (Liu et al., 2019; Papadopoulos and Agarwala, 2007; Shmakov et al., 2015; Zetsche et al., 2015).

### tracrRNA identification and PAM prediction

For CRISPR-Casπ system, tracrRNA 3’-region was determined by anti-repeat identification, transcriptome mapping and promoter prediction. Anti-repeats were searched against a 5k bp window upstream the CRISPR locus using blastn with (E-value <0.2) (Altschul et al., 1990). Subsequently, the meta-transcriptomic reads of the sludge sample were extracted and mapped to their native genome locus around the anti-repeat region to analyze the tracrRNA expression. The transcript coverage was calculated by log_10_ formula. Finally, the 5’-boundary of tracrRNA was determined by promoter prediction using BDGP-Promoter Prediction program (Reese, 2001). All tracrRNAs were determined in this manner as shown in Figure 1c.

To predict the PAM sequence for Casπ-1 and Casπ-2, all the spacers present in both CRISPR arrays were manually extracted and aligned against the default databases using CRISPRTarget to search the potential protospacer sequences (Biswas et al., 2013). Sequences 3-bp upstream of the identified protospacers were extracted and aligned to predict the PAM sequences. The PAMs ranking at the top for both Casπs were further used for plasmid cleavage *in vitro*.

### Plasmids construction

Bacteria- and human-codon optimized *casπ-1* and *casπ-2* genes were ordered from Sangon Biotech. For Casπ protein expression in *E.coli*, *Casπ* genes were cloned into pET28a-based vector with an N-terminal hexa-histidine tag and a SUMO tag by homologous recombination (One Step Seamless Cloning Mix, CWBIO). For the D537A and E643A mutations in RuvC domain, R390A and R392A mutations in OBD domain of Casπ-1, mutated fragments were PCR amplified via mutagenetic PCR primers containing mutated sequences and inserted into pET28a-based vector by homologous recombination. For PAM depletion assay, the plasmid library containing five randomized nucleotides upstream of the target sequence was constructed as previously described (Karvelis et al., 2015). For *in vitro* plasmids cleavage, pUC19-based plasmids containing target sequence with different PAMs were constructed via homologous recombination. For bacterial plasmid interference, pBAD-driven arabinose inducible *ccdB* toxin plasmid (p11-LacY-wtx1) was requested from Prof. Wei Li group in the institute of zoology, Chinese Academy of Sciences (Chen and Zhao, 2005). *Casπ* genes were cloned into MCSI of pCDFDuet vector by Gibson assembly with a sgRNA region, containing 2 *Sap*I sites for target spacer exchange by Golden Gate, inserting into MCSII of pCDFDuet (gRNA spacer sequences were listed in Table S3).

For constructing the EGFP report cell line, the CMV-driven fusion fragment of *MYH8* (270bp), a flanking sequence (32bp) and *EGFP* (1436bp) was cloned into psi-LVRU6MP vector by Gibson assembly. For cell editing assay, plasmids vector was obtained from circular PCR amplification of pBLO62.5 (Addgene plasmid #123124) with two primers respectively pairing to N-terminal and C-terminal NLS sequence (Tsuchida et al., 2022). Subsequently, *Casπ*s were inserted into the region downstream of the CMV promoter and N-terminal NLS by homologous recombination. Then, sgRNAs (containing 2 *Sap*I sites for spacer insertion) were inserted into the circular-PCR amplified vector containing *Casπ* genes with a U6 promoter and a poly-T terminal signal by homologous recombination. Primers containing the target spacer sequences were annealed and phosphorylated prior to Golden Gate assembly (*Sap*I restriction sites) for stuffer–spacer exchange insertion (target protospacer sequences were listed in Table S3).

A list of plasmids and a brief description are summarized in Table S2.

### Protein expression and purification

Casπ expression plasmids were transformed into *E.coli* BL21(DE3) (TIANGEN) and incubated overnight at 37 °C on LB-Kan^+^ agar plates (50 μg/ml Kanamycin). Single colony was overnight cultured as seed in LB-Kan^+^ medium (50 μg/ml Kanamycin) at 37 °C. Each 1 L of LB-Kan^+^ medium (50 μg/ml Kanamycin) was then inoculated with 100 ml seed culture and incubated at 37 °C. As the culture OD reached 1.0, the protein expression was induced with 0.2 mM IPTG for 20 hrs at 16 °C. Bacterial cells were collected and resuspended in lysis buffer (800 mM NaCl, 20 mM HEPES-Na pH 7.5, 10% Glycerol, 40 mM Imidazole, 1 mM TCEP and 1 mM PMSF) and lysed by sonication. The lysate was centrifuged at 15000 g for 80 min at 4 °C and applied to Ni-NTA gravity column. The resin was then washed with 20 column volumes (CV) of wash buffer (500 mM NaCl, 20 mM HEPES-Na pH 7.5, 10% Glycerol, 40 mM Imidazole, 1 mM TCEP), resuspended in 5 column volumes (CV) of tag-removal buffer (500 mM NaCl, 20 mM HEPES-Na pH 7.5, 10% Glycerol, 40 mM Imidazole, 1 mM TCEP and 0.6 μg/ml ulp1 protease) for 1hr incubation at 4 °C. Next, the supernatant was loaded into 5 ml HiTrap Heparin HP column (GE Healthcare) and eluted with a linear gradient of heparin elution buffer (A buffer: 20 mM HEPES-Na pH 7.5, 10% Glycerol, 1 mM TCEP; B buffer: 2 M NaCl, 20 mM HEPES-Na pH 7.5, 10% Glycerol, 1 mM TCEP). Elution fractions with Casπ were pooled together and concentrated using 30kD molecular weight cut-off centrifugal filters (Merck Millipore), and further purified by size exclusion chromatography column (Superdex 200 Increase 10/300, GE Healthcare) with S200 buffer (400 mM NaCl, 20 mM HEPES-Na pH 7.5, 10% Glycerol, 1 mM TCEP). Protein concentration were measured by NanoDrop One (Thermo Scientific) and stocked in −80 °C after flash-frozen in liquid nitrogen. The Casπ protein samples are usually stocked with the concentration of 300 μM. LbCas12a was expressed as previously described (Chen et al., 2018).

### *In vitro* transcription of CRISPR RNA

DNA sequences containing T7 RNA polymerase promoter upstream of the Casπ tracrRNA, crRNA and sgRNA were assembled by overlap PCR and validated by Sanger sequencing. The validated sequences were then PCR amplified as the template for *in vitro* transcription (IVT). All reactions were performed in IVT buffer (30 mM Tris pH 8.1, 25 mM MgCl_2_, 0.01% Triton, 2 mM spermidine) with 4 mM NTP mix and 0.4 mg/ml T7 RNA polymerase. The transcribed product was loaded into 10% Urea-PAGE with 2× formamide loading (95% formamide, 0.02% SDS, 0.02% BPB, 0.01% xylene cyanole FF, 1 mM EDTA) for electrophoresis. The gel region containing the target RNA band was extracted, smashed and soaked in soaking buffer (0.38 M NaAc pH 5.2, 0.8 mM EDTA, 0.8% SDS) for 8 hrs at 4 °C. The dissolved RNA was then concentrated using 3kD molecular weight cut-off centrifugal filters (Merck Millipore) and stocked in −80 °C. The RNA samples are usually stocked with the concentration of 50 μM. The RNA sequences and related description are listed in Table S3.

### PAM depletion assay and analysis

PAM depletion assay was performed as previously described with modification (Figure S2a) (Karvelis et al., 2015). Plasmids containing a PAM library were transformed into *E.coli* DH5α (TIANGEN) and incubated overnight at 37 °C on LB-Amp^+^ agar plates (100 μg/ml Ampicillin), and then all colonies were harvested to extract the plasmids using HighPure Maxi Plasmid Kit (TIANGEN). For cleavage reaction, sgRNA was diluted to the concentration of 30 μM in refolding buffer (50 mM KCl, 5 mM MgCl_2_) and refolded at 72 °C for 5 min, and then slowly cooled down to room temperature (RT). Subsequently, active RNP complexes were assembled with incubating 1 μM Casπ protein with 1.2 μM sgRNA in assembly buffer (100 mM NaCl, 10 mM HEPES-Na pH 7.5, 1 mM TCEP, 5 mM MgCl_2_) at RT for 30 min. The reaction was initiated by adding 20 nM plasmid and performed as three individual replicates in cleavage buffers (50 mM to 300 mM NaCl, 10 mM HEPES-Na pH 7.5, 1 mM TCEP, 10 mM MgCl_2_) at 37 °C for 1 hr, and then quenched with loading buffer (Gel Loading Dye Purple 6X, NEB) supplemented with 20 mM EDTA and 25 μg/ml heparin. The cleaved products were analyzed and purified by electrophoresis on the 1.2% agarose gel with GelRed staining (Vazyme). Then, the end of linearized products was repaired by T4 DNA polymerase (Thermo Fisher Scientific) with 1 mM dNTP (Sangon Biotech). And dA oligo was further added to the 3’ end of the products by Dreamtaq polymerase (Thermo Fisher Scientific) with 1 mM dATP (Sangon Biotech). Adapters with 3’ dT overhang were ligated with the products containing 3’ dA overhang by fast T4 DNA ligase (Beyotime). The DNA fragments containing the recognized PAM sequence were PCR amplified using a primer pairing to the adapter and the other primer pairing to the 120bp upstream region of the PAM. Next, the PCR amplified PAM-containing products were purified by VAHTS DNA Clean Beads (Vazyme) and further amplified by TIANSeq Fast DNA Library Prep Kit (TIANGEN) for Illumina Novaseq PE150 sequencing. In control groups, the plasmids were treated with blank buffer instead of Casπ effectors, and DNA fragments containing PAM library were directly amplified by two primers covering the PAM region for following process as described above. The depletion fold-change for each PAM were analyzed using the number of matched reads in Casπ and control groups normalized with total reads.

A list of depleted PAMs and related fold-change values are summarized in Table S1.

### *In vitro* cleavage assays

For cleavage assays with labeled NTS, the dsDNA substrate was prepared by PCR extension using a 65 nt ssDNA template and a 5’-cy5 labeled 16nt primer (order from Sangon Biotech). Then the extended dsDNA was purified by DNA Clean & Concentrator-25 (Zymo Research) and diluted to 1 μM in nuclease-free water (Invitrogen). The sgRNA was diluted to the concentration of 30 μM in refolding buffer (50 mM KCl, 5 mM MgCl_2_) and refolded as described above. Subsequently, Casπ effectors were assembled in a 1:1.2 protein to sgRNA ratio (1 μM Casπ protein and 1.2 μM refolded sgRNA) in assembly buffer (100 mM NaCl, 10 mM HEPES-Na pH 7.5, 1 mM TCEP, 5 mM MgCl_2_) at RT for 30 min. The reaction was started by mixing 1 μM RNP with 20 nM dsDNA substrate in cleavage buffer (150 mM NaCl, 10 mM HEPES-Na pH 7.5, 1 mM TCEP, 10 mM MgCl_2_) at 37 °C and aliquots were collected at the following time points: 0, 2, 5, 15, 30, 60, 90 and 120 min. For biochemical screenings, only the reaction buffers were modified accordingly, such as the salt concentration (50, 150, 300 or 450 mM NaCl with 10 mM HEPES-Na pH 7.5, 1 mM TCEP, 10 mM MgCl_2_), type of divalent ions (10 mM Mg^2+^, Mn^2+^, Ca^2+^ or Co^2+^ with 150 mM NaCl, 10 mM HEPES-Na pH 7.5, 1 mM TCEP) and temperatures (25, 30, 37, 45, 55 or 65 °C with 150 mM NaCl, 10 mM HEPES-Na pH 7.5, 1 mM TCEP, 10 mM MgCl_2_). The products were analyzed as mentioned above.

For cleavage assays with labeled TS, the 5’-cy5 labeled TS ssDNA were synthesized by Sangon Biotech and diluted to 10 μM in nuclease-free water (Invitrogen). dsDNA was prepared by mixing 5’-cy5 labeled TS and unlabeled complementary oligo at the molar ratio of 1:1.2 in annealing buffer (10 mM HEPES-Na pH 7.5, 150 mM KCl), followed by heating for 5 min at 95 °C and slow cool down to RT. Cleavage reactions were initiated by mixing 1 μM RNP with 20 nM ssDNA or dsDNA substrate in cleavage buffer (150 mM NaCl, 10 mM HEPES-Na pH 7.5, 1 mM TCEP, 10 mM MgCl_2_) at 37 °C and product aliquots were collected at the following time points: 0, 2, 5, 15, and 60 min.

For mismatched-cleavage assay, the dsDNA substrates with single mismatches were prepared by PCR extension using a 65 nt ssDNA template with single mismatch and a 5’-cy5 labeled 16nt primer (order from Sangon Biotech). Then the extended dsDNA was purified by DNA Clean & Concentrator-25 (Zymo Research) and diluted to 1 μM in nuclease-free water (Invitrogen). Cleavage reactions were initiated by mixing 1 μM RNP with 20 nM dsDNA substrate in cleavage buffer (150 mM NaCl, 10 mM HEPES-Na pH 7.5, 1 mM TCEP, 10 mM MgCl_2_) at 37 °C and product aliquots were collected at 1 hour.

For *trans*-cleavage assay, 1 μM Casπ or LbCas12a RNP was first incubated with 1.5 μM dsDNA or ssDNA activator at 37 °C for 30min. Then 20 nM 5’-cy5 labeled random 60-nt ssDNA was mixed into the reaction. The product aliquots were collected at the following time points: 0, 2, 5, 15, 30, 60, 90 and 120min.

All cleavage products collected above were quenched with 2×Urea-loading buffer (8 M urea and 2 mM Tris-Cl pH 7.5) supplemented with 20 mM EDTA and 25 μg/ml heparin, then analyzed in 15% urea-PAGE and visualized using Amersham Typhoon 5 (GE Healthcare). Product bands were quantified using ImageJ and cleaved fraction was calculated using the intensity of product bands divided by input intensity.(Schneider et al., 2012) Curves of cleavage efficiency were plotted using a One-Phase-Decay model in Prism 8 (GraphPad).

For plasmids cleavage assay, 1 μM Casπ RNP effectors were incubated with 20 nM target plasmids at 37 °C for 30 min and then quenched with loading buffer (Gel Loading Dye Purple 6X, NEB) supplemented with 20 mM EDTA and 25 μg/ml heparin. The samples were analyzed by electrophoresis on a 1.2% agarose gel with GelRed staining (Vazyme). For non-labeled dsDNA cleavage assay, the dsDNA target was PCR amplified from the plasmid containing the protospacer and purified by DNA Clean & Concentrator-25 (Zymo Research). The reaction was initiated with incubating 1 μM Casπ RNP effectors with 20 nM dsDNA target at 37 °C for 30 min and then quenched with loading buffer (Gel Loading Dye Purple 6X, NEB) supplemented with 20 mM EDTA and 25 μg/ml heparin. The samples were analyzed by electrophoresis on the 1.2% agarose gel with GelRed staining.

All experiments were performed at least by three times for replicability. A list of oligonucleotides used in this study and related description are summarized in Table S3.

### Cleavage sites determination

The cleavage products and sites on dsDNA were analyzed by electrophoresis using 15% urea-PAGE as described above. To determine the cleavage sites on plasmids, linearized plasmids were purified and subjected to NGS libraries construction for Illumina Novaseq PE150 sequencing as previously described in PAM depletion assay. Paired-end reads were mapped to the target sequence using BWA and 3’-ends were selected to determine the cleavage sites. The abundance of each site was normalized to the total reads and plotted using Prism 8 (GraphPad).

### Plasmid interference in bacteria

*E. coli* BW25141 cells were requested from prof. Guangdong Shang group in College of Life Sciences, Nanjing Normal University. *E. coli* BW25141 competent cells carrying the *ccdB* toxin plasmid (p11-LacY-wtx1) was prepared following the protocol previously described.(Chen and Zhao, 2005) For each group, 200 ng plasmid expressing Casπ and sgRNA (ccdB-targeting or non-targeting) were electroporated into 50 μL competent cells with 0.2 cm cuvette (BIO-RAD) under 2.5 kV using Eppendorf eporator (Eppendorf). After 1.5 hrs of recovering in 5 mL SOC medium (Sangon Biotech) under 37 °C, the bacterial cells were enriched by centrifugation and resuspended with 5 mL liquid LB-Strep^+^ medium (50 μg/mL streptomycin), and cultured for an extra 8 hrs. Subsequently, to investigate the effects to bacterial survival by Casπ editing, 5 µL of culture with gradient dilutions from 10^0^ to 10^-7^ was spotted onto the LB-Amp^+^ agar plates (100 μg/mL ampicillin) or LB-Strep^+^-Ara^+^ agar plates (50 μg/mL streptomycin, 10 mM arabinose) respectively and incubated overnight at 37 °C. In the meantime, to validate the transformation efficacy of Casπ-sgRNA expression plasmids, 10 μL of culture was spread on LB-Strep^+^ agar plates (50 μg/mL streptomycin) for overnight incubation at 37 °C, colony number on each plate was manually counted. 5 μL of edited bacterial cells was used for PCR validation of the plasmid interference with Phanta Max Super-Fidelity DNA Polymerase Mastermix (Vazyme).

### Construction of EGFP report cell line

To obtain a natural target sequence with diverse targeting windows (different GC contents and PAMs), a sequence survey was performed in mouse genome. Via screening by 20 nt-window, we allocated a 270 bp fragment within the *Mus musculus* myosin heavy polypeptide 8 (*MYH8*) exon (NCBI accession: NM_177369.3 (3650-3919)) which presents a well distribution of targeting windows with various GC contents (30% to 85%) and PAMs (Table S1). And this region shows low sequence similarity to human genome. Frameshifting *EGFP* (3n+2) was created by fusing the *MYH8* fragment, a 32 bp random flanking sequence and *EGFP* ORF (1436 bp). The *MYH8-EGFP* was further inserted into lentiviral packaging plasmid. The LV-MAX lentiviral production system (Thermo Fisher Scientific) was used to produce the lentivirus for inserting the *MYH8-EGFP* (3n+2) fragments into HEK293A cell genome via infection. The selection and enrichment of genome-modified cells were performed according to the manufacturer’s protocol (Thermo Fisher Scientific).

### Gene editing assay in human cells

For EGFP activation editing assay in human cells, the EGFP HEK293A report cells were cultured in DMEM (Gibco) medium supplemented with 10 % (v/v) FBS (Gemini) and 1% (v/v) penicillin streptomycin (Gibco) at 37 °C in 5% CO_2_. About 8.0×10^4^ cells were seeded onto the each well of 48-well plate for ∼16 hr incubation. When the cell confluency reached 60-70%, 300 ng plasmid expressing NLS-Casπ- or Cas9-P2A-PuroR-NLS with sgRNA (MYH8-targeting and non-targeting) was transfected into the cells within each well using lipofectamine 3000 (Life Technologies) according to the manufacturer’s protocols. One-day after transfection, the old medium was replaced by fresh DMEM-Puro^+^ medium ( 1.5 μg/mL puromycin, Sigma) for 3-day culturing. Then the enriched cells were further cultured for another 3 days using fresh DMEM medium without puromycin for gene-editing analysis. The EGFP signal was observed with fluorescent microscopy (Nikon Eclipse TS2FL fluorescence microscope). Edited cells were also collected and stored at −80 °C. For more endogenous gene editing assay, the HEK293T cells were treated the same as mentioned above, but transfected with NLS-Casπ-P2A-PuroR-NLS with sgRNA targeting other endogenous genes.

A list of targeting sequences is summarized in Table S3.

### Evaluation of gene editing efficacy

For T7E1 assay, Genome of edited cells was extracted using Ezup Column Animal Genomic DNA Purification Kit (Sangon Biotech). The edited genome was used as the template for the PCR amplification of target region using Phanta Max Super-Fidelity DNA Polymerase Mastermix (Vazyme) (primers listed in Table. S3). The PCR product was gel-purified, and about 200 ng purified DNA was re-annealed for T7 endonuclease I (T7E1) cleavage assay referring to the manufacturer’s protocol (Vazyme). Cleavage products were analyzed by electrophoresis using 2% agarose gel with GelRed staining (Vazyme).

For NGS, ∼210 bp regions nearby the target protospacers were amplified via PCR with Q5 polymerase (NEB) and primers containing Illumina adaptor sequences. Amplicons were verified by electrophoresis using 2% agarose gel with GelRed staining (Vazyme), purified by VAHTS DNA Clean Beads according to the manufacturer’s protocol (Vazyme) and further loaded onto Illumina Novaseq PE150 sequencing by Tianjin Novogene Bioinformatic Technology Co., Ltd. Sequencing reads were analyzed by CRISPResso2 with the following parameters: quantification window centered at 3 bp for Casπ-1 (2 bp for Casπ-2, 1 bp for Cas12a and -3 bp for Cas9) according to cleavage sites of both Casπs (Figure S2g, h), quantification window size of 14 bp for both Casπs (8 bp for Cas9), and plot window size of 40 bp (to visualize large indels).(Clement et al., 2019) Cells treated with plasmids carrying codon optimized Cas genes with a non-targeting sgRNA were evaluated at every spacer sequence within every read as a negative control. Percentage of each indel plotted (regardless of substitution) was based on the results of modified reads from the CRISPResso2 output. For the indel size distribution plots, unmodified reads (indel length of 0 bp) were plotted as 0% of the total reads for clarify and the remaining reads were grouped and plotted based on the modified results.

### Reconstitution of Casπ R-loop complex

Deactivated Casπ-1 (dCasπ-1, D537A, E643A) was purified as described above. The sgRNA was diluted to 40 μM in refolding buffer (50 mM KCl, 5 mM MgCl_2_) and refolded as mentioned above. The dCasπ-1-sgRNA binary was reconstituted by incubating 20 μM dCasπ-1 and 25 μM sgRNA for 30 min at RT in a total volume of 150 μl assembly buffer (100 mM NaCl, 10 mM HEPES-Na pH 7.5, 1 mM TCEP, 5 mM MgCl_2_).To facilitate the R-loop formation, the bubbled dsDNA substrate with 10 nt mismatch in the protospacer was used for R-loop ternary complex assembly. The bubbled dsDNA was diluted to 30 μM in 150 μl assembly buffer, and mixed with 150 μl binary complex at RT for 30min incubation. Subsequently, the assembled sample was purified by size exclusion column (Superdex 200 Increase 10/300, GE Healthcare) in SEC buffer (150 mM NaCl, 10 mM HEPES-Na pH 7.5, 1 mM TCEP, 0.1% Glycerol, 5 mM MgCl_2_, 0.1% Glycerol) at 4 °C. After flash freezing by liquid nitrogen, the aliquots of purified sample were stocked in −80 °C. The reconstituted complex was usually stocked at the concentration of 3 μM. A list of DNA oligonucleotides and sgRNA sequences with brief descriptions are presented in Table S3.

### Cryo-EM sample preparation and data collection

4 μL purified Casπ R-loop complex (∼1.5 μM) was cross-linked by BS3 (Sigma-Aldrich) and applied to the graphene oxide grid (Quantifoil Au 1.2/1.3, 300 mesh), which was glow-discharged (in a HARRICK PLASMA) for 10 s at middle level after 2 min evacuation. The grid was then blotted by a pair of 55 mm filter papers (Ted Pella) for 0.5 s at 22 °C with 100% humidity, and flash-frozen in liquid ethane using FEI Vitrobot Marke IV. Cryo-EM data were collected on a Titan Krios electron microscope operated at 300 kV equipped with a Cs-corrector and Gatan K3 direct electron detector with Gatan Quantum energy filter using EPU. Micrographs were recorded in counting mode at a nominal magnification of 105,000×, resulting in a physical pixel size of 0.856 Å per pixel. The defocus was set between −1.5 μm to −2.5 μm. The total exposure time of each movie stack led to a total accumulated dose of 50 electrons per Å^2^ which fractionated into 32 frames. More parameters for data collection are shown in Table S4.

### Image processing and 3D reconstruction

The raw dose-fractionated image stacks were 2 × Fourier binned, aligned, dose-weighted, and summed using MotionCor2 (Zheng et al., 2017). CTF-estimation, blob particle picking, 2D reference-free classification, initial model generation, final 3D refinement and local resolution estimation were performed in cryoSPARC (Punjani et al., 2017). Two rounds of 3D reference-based classification were performed in RELION (Kimanius et al., 2016). The details of data processing were summarized in Figure S5 and Table S4.

### Model building and refinement

The initial protein model was generated using AlphaFold2 and manually revised in UCSF-Chimera and Coot (Emsley and Cowtan, 2004; Jumper et al., 2021; Pettersen et al., 2004). The DNA substrates and sgRNA were manually built in Coot based on the cryo-EM density. The complete model was refined against the EM map by PHENIX in real space with secondary structure and geometry restraints (Adams et al., 2010). The final model was validated in PHENIX software package. The structural validation details for the final model are summarized in Table S4.

### Quantification and statistical analysis

Statistical details for each experiment can be found in the figure legends and the corresponding methods details. Graphs show the average of replicates with individual points overlaid, unless stated otherwise.

## Supporting information

Integrated Supplementary file

## Resource availability

### Lead contact

Readers are welcome to comment on the online version of the paper. Correspondence and requests for materials should be addressed to the lead contact J.J.G.L.

### Materials availability

Plasmids generated in this study will be deposited to Addgene or are available upon request.

### Data and code availability

The electron density maps have been deposited to the Electron Microscopy Data Bank (EMDB) under the accession numbers of EMD-33983 which are publicly available as of the data of publication. The atomic coordinates and structure factors have been deposited to the Protein Data Bank (PDB) under the accession number of 7YOJ which are publicly available as of the data of publication. The raw cryo-EM micrographs and movies used in this study will be shared by corresponding author upon request. The raw sequencing result of metagenome is uploaded to NCBI database with the accession ID of PRJNA857874. Any additional information required to re-analyze the data reported in this paper is available from the corresponding author upon request.

## Acknowledgements

EM data were collected at the Tsinghua Cryo-EM facility and Shuimu Bioscience. The data were analyzed using the Bio-Computation platform at the Tsinghua University Branch of the Chinese National Center for Protein Sciences (Beijing). We thank the supports from the Tsinghua University Technology Center for Protein Research, Genome Sequencing and Analysis at Tsinghua University. We thank J.L. Lei, X.M. Li and X.D. Li for expert electron microscopy assistance. We thank T. Yang, Y.K. Wang, A.B. Jia for computational support. We thank D. Chia, Y. Lin and N. Liu for their kind revise on this manuscript. The work was supported by the National Key Research and Development Program (2022YFF1002801 to J.J.G.L.), the Ministry of Agriculture and Rural Affairs of China (J.J.G.L.), National Natural Science Foundation of China (Grant No.32150018 to J.J.G.L and No.32101195 to S.Z.), and start-up funds from Tsinghua University, Beijing (J.J.G.L.).

## Author contributions

J.J.G.L supervised the project. J.J.G.L, J.W, A.S., C.P.L., Z.C., S.Z. and S.L. designed the experiments. C.P.L., S.Z., K.W, S.L. and J.L collected and analyzed the environmental metagenome. C.P.L. and S.Z. built the bioinformatic pipeline and discovered the new system. A.S. purified the Casπ proteins and performed all the biochemical assays and analysis. J.W. and C.P.L. did the structural analysis and built the atomic model. Z.C., A.S., D.L., Y.Y. and Y.Z. did the gene editing experiments in bacterial and mammalian cells. J.J.G.L, J.W., A.S., C.P.L. and Z.C. wrote the manuscript with help from all authors.

