## Supplementary material for "The compact Casπ (Cas12l) ‘bracelet’ provides a unique structural platform for DNA manipulation": Integrated Supplementary file


**List:**

Figure S1-S11

Table S1-S4

Video S1

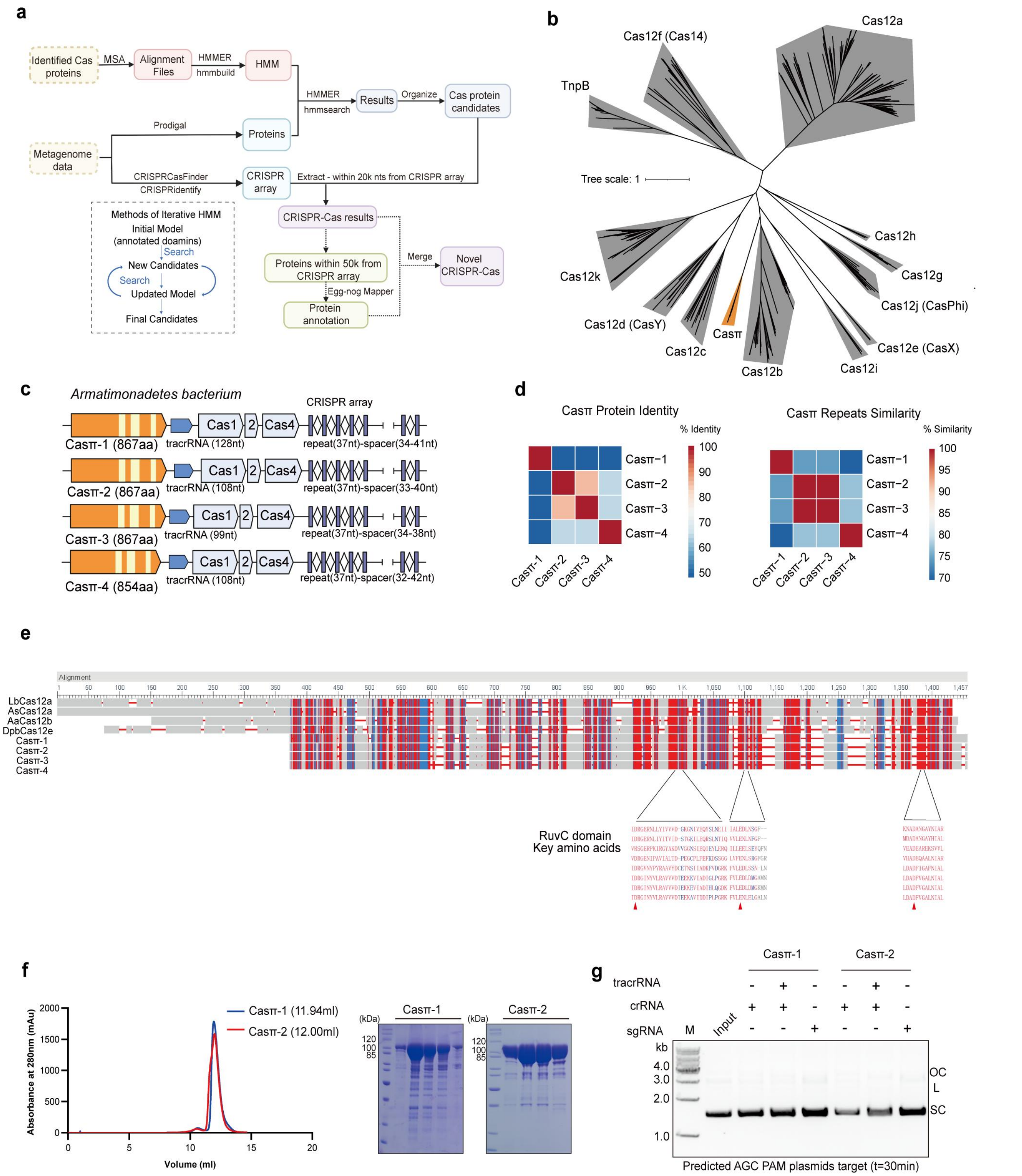


**Figure. S1 Casπ contains an active RuvC domain for DNA cleavage.**

(**a**) Bioinformatics pipeline of identifying novel CRISPR-Cas systems. In general, the prediction of CRISPR-Cas systems is based on the identification of CRISPR arrays and Cas proteins. The architecture of CRISPR array is relatively conserved and can be predicted by public software, such as CRISPRCasFinder and CRISPRidentify. Meanwhile, Cas proteins can be predicted by canonical protein homology search tools, such as HMMER and Pfam based on Hidden Markov models (HMM). However, these existed tools can only identify 'new' proteins which share relatively high sequence similarity to annotated Cas proteins. Therefore, we develop an unbiased workflow using iterative HMM to identify novel Cas candidates. For example, to identify novel type V Cas nucleases, RuvC domains from annotated Cas12a nucleases in the Pfam database (Pfam accession number: PF18516) were used to build an initial HMM for the first round searching of RuvC-containing proteins in all accessible databases. 1,349 protein candidates within 300 to 1600 amino acids (un-annotated in Pfam) were found, and all 'RuvC' domains within those candidates are used to build an improved HMM for the next round search of new candidates in all accessible databases. Using this iterative pipeline, the HMM was continuously updated with new candidates found in the current round and yielded to an updated pool of Cas-nuclease candidates from the next round search. A typical 2~10 rounds of iterative HMM and search were used for generating the final pool of novel type V Cas nucleases. Further, we filtered the candidates in the final pool by their functions annotated by eggNOG mapper, similarities to the final HMM (E-value), sizes and the confidence of putative CRISPR arrays for manual screening. Particularly in this study, Casπ showed up in the candidate pool with three rounds of iterative HMM and search.

(**b**) Maximum likelihood phylogenic analysis of Casπ with type V subtypes a-k. Casπ proteins are outlined in orange, with other subtypes in grey. Bootstrap=1500, Casπ protein sequences are shown in Table S1.

(**c**) Genomic architectures of all four CRISPR-Casπ systems.

(**d**) Similarity matrix built and visualized using heatmap for Casπ protein sequences (left) and repeat sequences (right).

(**e**) Multiple sequence alignment of Casπ with LbCas12a, AsCas12a, AaCas12b and DpbCas12e. Key amino acids of RuvC domain are marked by red arrows.

(**f**) Purification of both Casπ proteins. The SEC curves were aligned referring to the injection volume. The peak fractions were analyzed by SDS-PAGE.

(**g**) *In vitro* cleavage of plasmids containing predicted AGC PAM by crRNA only, tracrRNA and crRNA pair or sgRNA by Casπ effectors (SC, supercoiled plasmids; L, linearized plasmids; OC, open-circle plasmids).

**a**

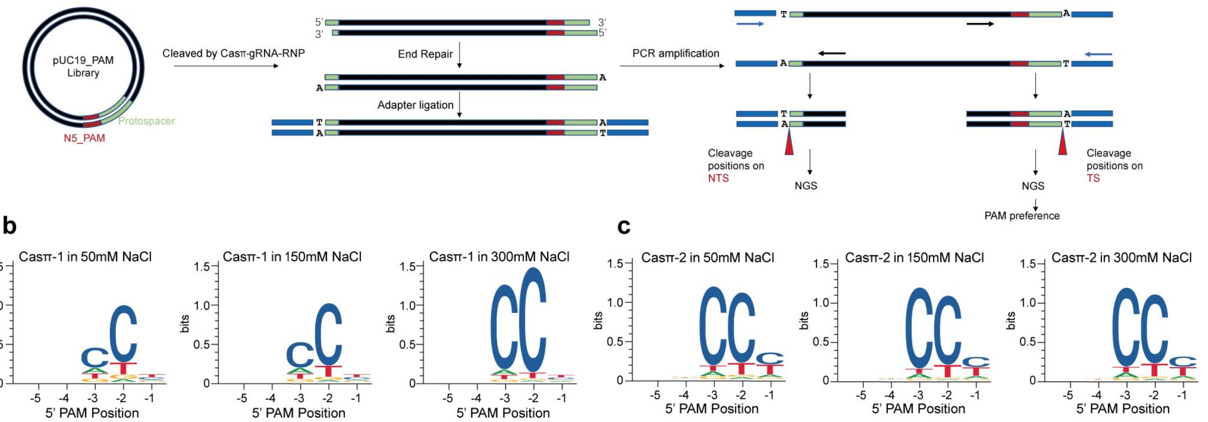


**d**

| Subtype | Protein | Length(aa) | PAM | tracrRNA/scoutRNA |
|---|---|---|---|---|
| | Casπ | 854-867 | CCN | tracrRNA |
| TypeV-A | Cas12a | 1100-1300 | TTTV | No |
| TypeV-B | Cas12b | 1100-1500 | TTN | tracrRNA |
| TypeV-C | Cas12c | 1200-1300 | TN | scoutRNA |
| TypeV-D | Cas12d (CasY) | 1200-1300 | TR | No |
| TypeV-E | Cas12e (CasX) | 986 | TTCN | tracrRNA |
| TypeV-F | Cas12f (Cas14) | 500-800 | TTTN | tracrRNA |
| TypeV-G | Cas12g | 800-900 | No | tracrRNA |
| TypeV-H | Cas12h | 900-1000 | RTR | No |
| TypeV-I | Cas12i | ~1100 | TTN | No |
| TypeV-J | Cas12j (CasΦ) | 700-800 | GTN | No |
| TypeV-K | Cas12k | ~700 | TBN | tracrRNA |

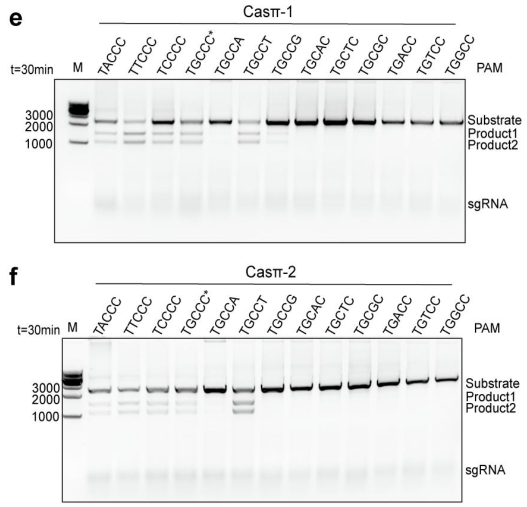


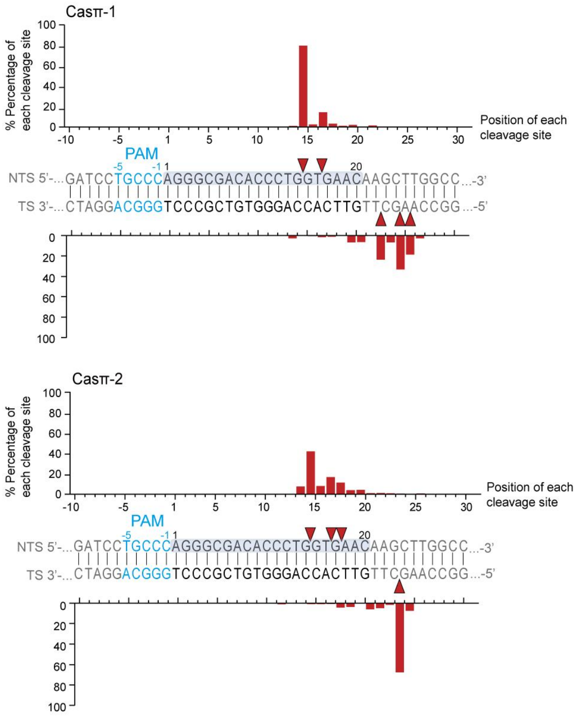


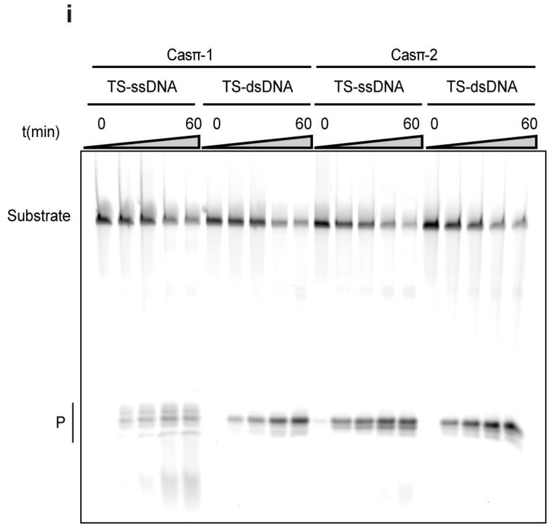

**Figure. S2 Casπ recognizes protospacer with 5’ C-rich PAM and creates a staggered end with 5’ overhang.**

(**a**) Pipeline of N5-PAM screening assay.

(**b**) PAM preference of Casπ-1 effector in buffers with different salt concentrations.

(**c**) PAM preference of Casπ-2 effector in buffers with different salt concentrations.

(**d**) Comparison of the Cas nuclease size, PAM preference and gRNA composition among type V Casπ and Cas12a-k systems.

(**e** and **f**) PAM preference validation for Casπ-1 (**e**) and Casπ-2 effectors (**f**) with dsDNA substrate. The PAM sequence designed in each dsDNA target is shown on top of each lane. M means DNA marker.

(**g** and **h**) Percentage distribution of Casπ-1 (**g**) and Casπ-2 effectors (**h**) cleavage sites analyzed by NGS data. (Percentage value was calculated by the number of reads belonging to each cleavage site divided by the number of total reads for all sites, X axis indicates the position of each nucleotide).

(**i**) Comparison of ssDNA and dsDNA cleavage by Casπ effectors. The ssDNA target containing complementary sequence to sgRNA spacer was labeled as TS-ssDNA. The dsDNA target containing complementary sequence to sgRNA spacer was labeled as TS-dsDNA.

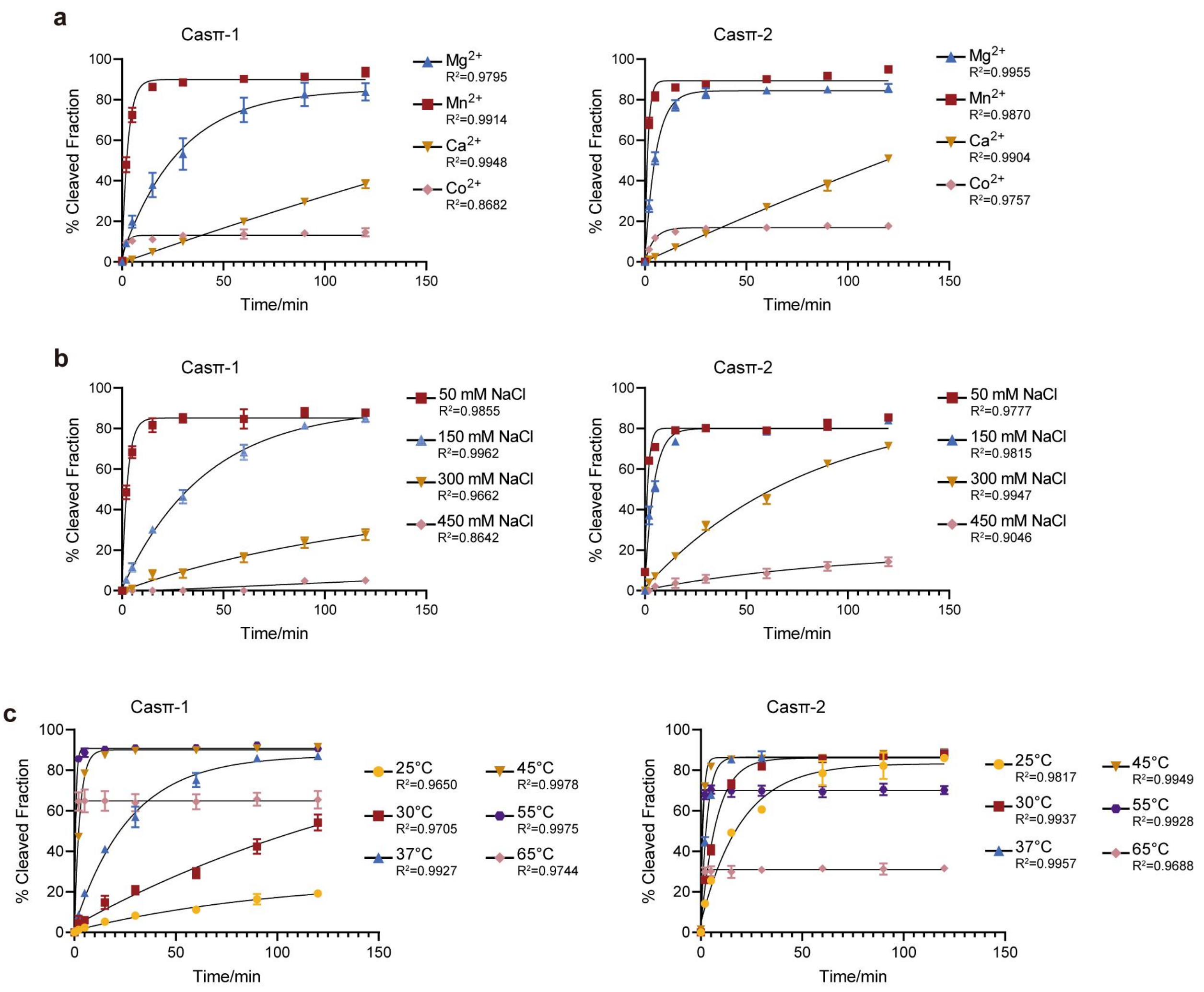


**Figure. S3 Cleavage efficiency of Casπ nuclease in various biochemical conditions.**

(**a**) The plot for cleavage efficiency based on three replicates of the dsDNA cleavage by Casπ effectors using different divalent ions. NTS denotes the non-target strand DNA which is cy5-labeled at 5' end. Bottom, P means products (n=3 each; means ± SD). The rate constant k values for Casπ-1 in $Mg^{2+}$, $Mn^{2+}$, $Ca^{2+}$ and $Co^{2+}$ are 0.03460, 0.3497, 0.0015 and 0.5170 (fraction/minute), respectively. The rate constant k values for Casπ-2 in $Mg^{2+}$, $Mn^{2+}$, $Ca^{2+}$ and $Co^{2+}$ are 0.1833, 0.6899, 0.0015 and 0.2227 (fraction/minute), respectively.

(**b**) The plot for cleavage efficiency based on three replicates of the dsDNA cleavage by Casπ effectors using different salt concentrations. (n=3 each; means ± SD). The rate constant k values for Casπ-1 in 50mM NaCl, 150mM NaCl, 300mM NaCl and 450mM NaCl are 0.3730, 0.02434, 0.0079 and $4.717e^{-6}$ (fraction/minute), respectively. The rate constant k values for Casπ-2 in 50mM NaCl, 150mM NaCl, 300mM NaCl and 450mM NaCl are 0.6748, 0.2285, 0.01228 and 0.01027 (fraction/minute), respectively.

(**c**) The plot for cleavage efficiency based on three replicates of the dsDNA cleavage by Casπ effectors in different temperatures. (n=3 each; means ± SD). The rate constant k values for Casπ-1 in 25 ℃, 30 ℃, 37 ℃, 45 ℃, 55 ℃ and 65 ℃ are 0.00987, 0.005353, 0.03663, 0.3849, 1.426 and 2.781 (fraction/minute), respectively. The rate constant k values for Casπ-2 in 25 ℃, 30 ℃, 37 ℃, 45 ℃, 55 ℃ and 65 ℃ are 0.05026, 0.1285, 0.3340, 0.8713, 1.754 and 1.640 (fraction/minute), respectively.

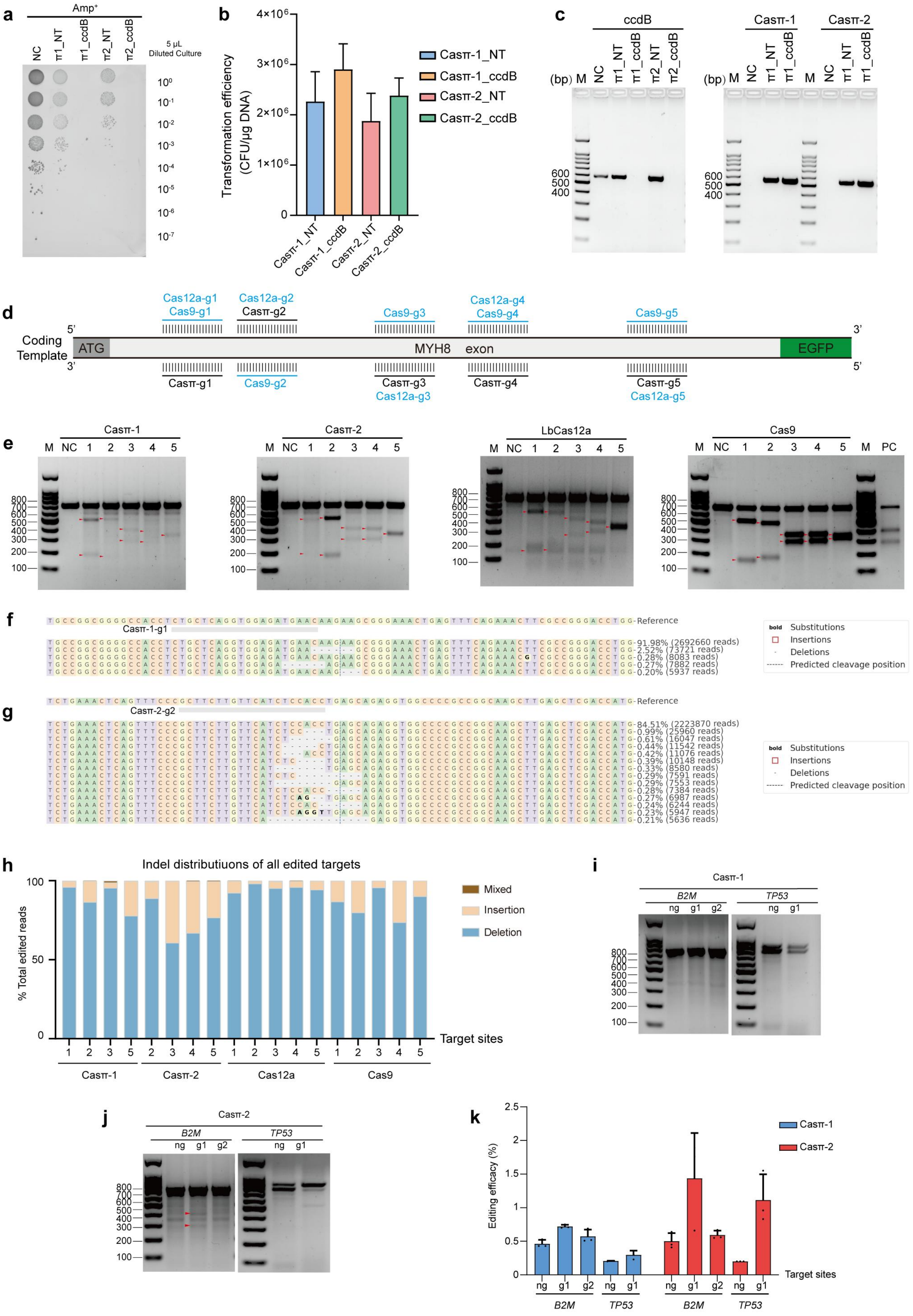
a
Amp+
NC
π1_NT
π1_ccdB
π2_NT
π2_ccdB
5 μL Diluted Culture
10^0 10^-1 10^-2 10^-3 10^-4 10^-5 10^-6 10^-7
b
Transformation efficiency (CFU/μg DNA)
Casπ-1_NT
Casπ-1_ccdB
Casπ-2_NT
Casπ-2_ccdB
c
ccdB
Casπ-1
Casπ-2
(bp) M
600 500 400
d
Coding Template
ATG
MYH8 exon
EGFP
Cas12a-g1
Cas9-g1
Cas12a-g2
Casπ-g2
Cas9-g3
Cas12a-g4
Cas9-g4
Cas9-g5
Casπ-g1
Cas9-g2
Casπ-g3
Cas12a-g3
Casπ-g4
Casπ-g5
Cas12a-g5
e
Casπ-1
Casπ-2
LbCas12a
Cas9
M NC 1 2 3 4 5
PC
800 700 600 500 400 300 200 100
f
Reference
Casπ-1-g1
91.98% (2692660 reads)
2.52% (73721 reads)
0.28% (8083 reads)
0.27% (7882 reads)
0.20% (5937 reads)
bold Substitutions
Insertions
Deletions
Predicted cleavage position
g
Reference
Casπ-2-g2
84.51% (2223870 reads)
0.99% (25960 reads)
0.61% (16047 reads)
0.44% (11542 reads)
0.42% (11076 reads)
0.39% (10148 reads)
0.33% (8580 reads)
0.29% (7591 reads)
0.29% (7553 reads)
0.28% (7384 reads)
0.27% (6987 reads)
0.24% (6244 reads)
0.23% (5947 reads)
0.21% (5636 reads)
h
Indel distributiuons of all edited targets
% Total edited reads
Mixed
Insertion
Deletion
Target sites
Casπ-1
Casπ-2
Cas12a
Cas9
i
Casπ-1
B2M
TP53
ng g1 g2
j
Casπ-2
B2M
TP53
ng g1 g2
k
Editing efficacy (%)
Casπ-1
Casπ-2
Target sites
B2M
TP53

**Figure. S4 Casπ mediated gene manipulation in prokaryotic and eukaryotic cells.**

(**a**) Bacteria survival result on LB-$Amp^{+}$ agar plates. (NT, plasmid with Casπ and non-target sgRNA; ccdB, plasmid with Casπ and sgRNA targeting *ccdB* gene).

(**b**) Transformation efficiency of Casπ plasmids validated on LB-$Strep^{+}$ agar plates (n=3 each, mean ± SD).

(**c**) Left panel, PCR detection of ccdB plasmids in edited cells. The two PCR primers respectively locates at the cleavage-site upstream and downstream. Right panel, PCR detection of Casπ plasmids in edited cells.

(**d**) Distribution of targeting sites across *MYH8* exon for Casπ, Cas12a and Cas9 effectors.

(**e**) T7E1 cleavage on the re-annealed target amplified from edited genome of Casπ-1, Casπ-2, Cas12a and Cas9. Cleavage products were indicated by red arrows. PC indicates the positive control offered in the manufacture kit for T7E1 assay.

(**f**) Summary of indels generated by Casπ-1 on the MYH8-target site1. Only indels with frequencies ≥ 0.2% of the total reads are shown.

(**g**) Summary of indels generated by Casπ-2 on the MYH8-target site2. Only indels with frequencies ≥ 0.2% of the total reads are shown.

(**h**) INDEL distributions of all five targets by Cas effectors analyzed by NGS of the MYH8 target (Mixed means insertion and deletion, only targets with editing efficacies ≥ 1% are shown).

(**i**) T7E1 cleavage on the re-annealed target amplified from edited genome of Casπ-1 on *B2M* and *TP53* (ng means non-target sgRNA; n=3 each, mean ± SD).

(**j**) T7E1 cleavage on the re-annealed target amplified from edited genome of Casπ-2 on *B2M* and *TP53*. Cleavage products were indicated by red arrows (n=3 each, mean ± SD).

(**k**) Editing efficacies determined by NGS for 3 more targets mediated by Casπ-1 and Casπ-2 (n=3 each, mean ± SD).

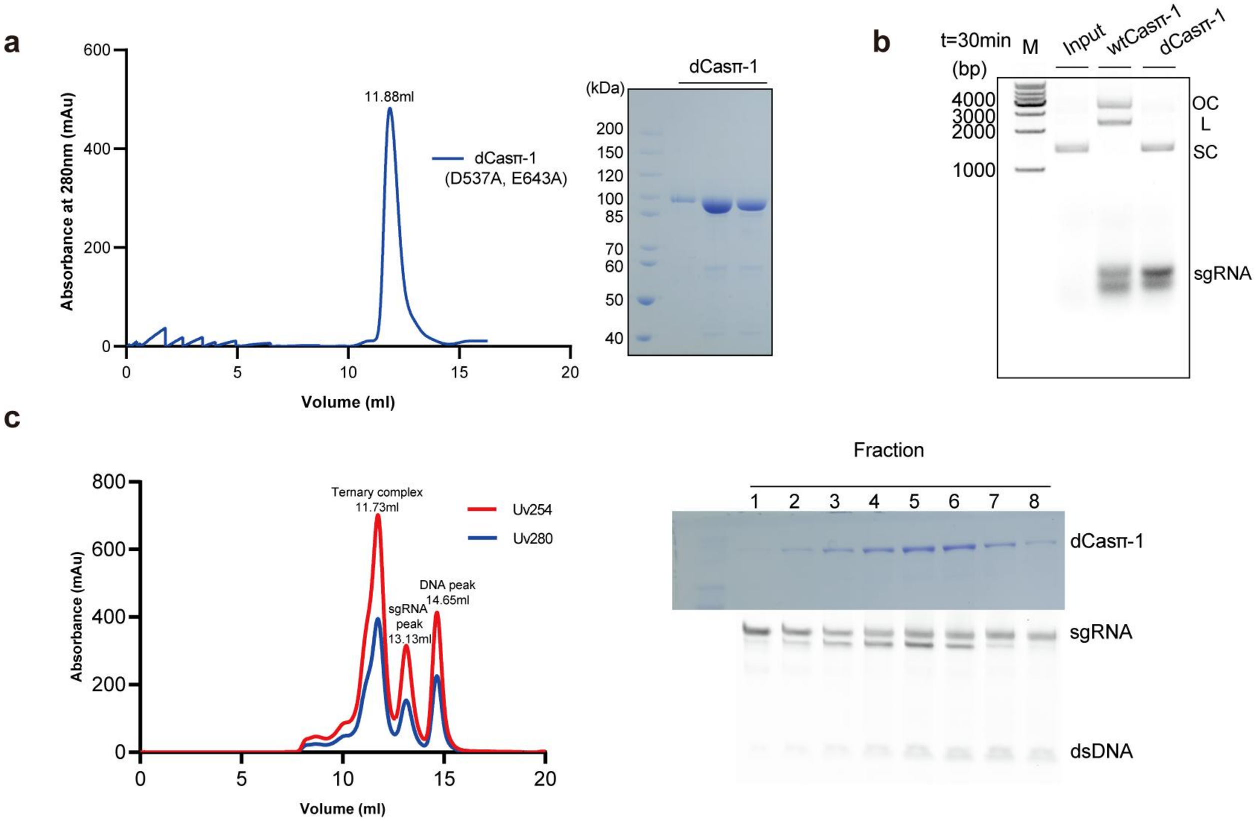


**Figure. S5 Reconstitution of Casπ-sgRNA-dsDNA R-loop complex.**

(**a**) Purification of deactivated Casπ-1 (D537A, E643A) protein. The peak fractions were analyzed by SDS-PAGE.

(**b**) Plasmid cleavage by wild type Casπ-1 (wtCasπ-1) and deactivated Casπ-1(dCasπ-1). (SC, supercoiled plasmids; L, linearized plasmids; OC, open-circle plasmids).

(**c**) Reconstitution and purification of dCasπ-1-sgRNA-dsDNA R-loop complex by size-exclusion chromatography. The reconstituted complex was analyzed by SDS-PAGE (top) and urea-PAGE (bottom). The sample labeled with 5 from the peak fraction was used for cryo-EM study.

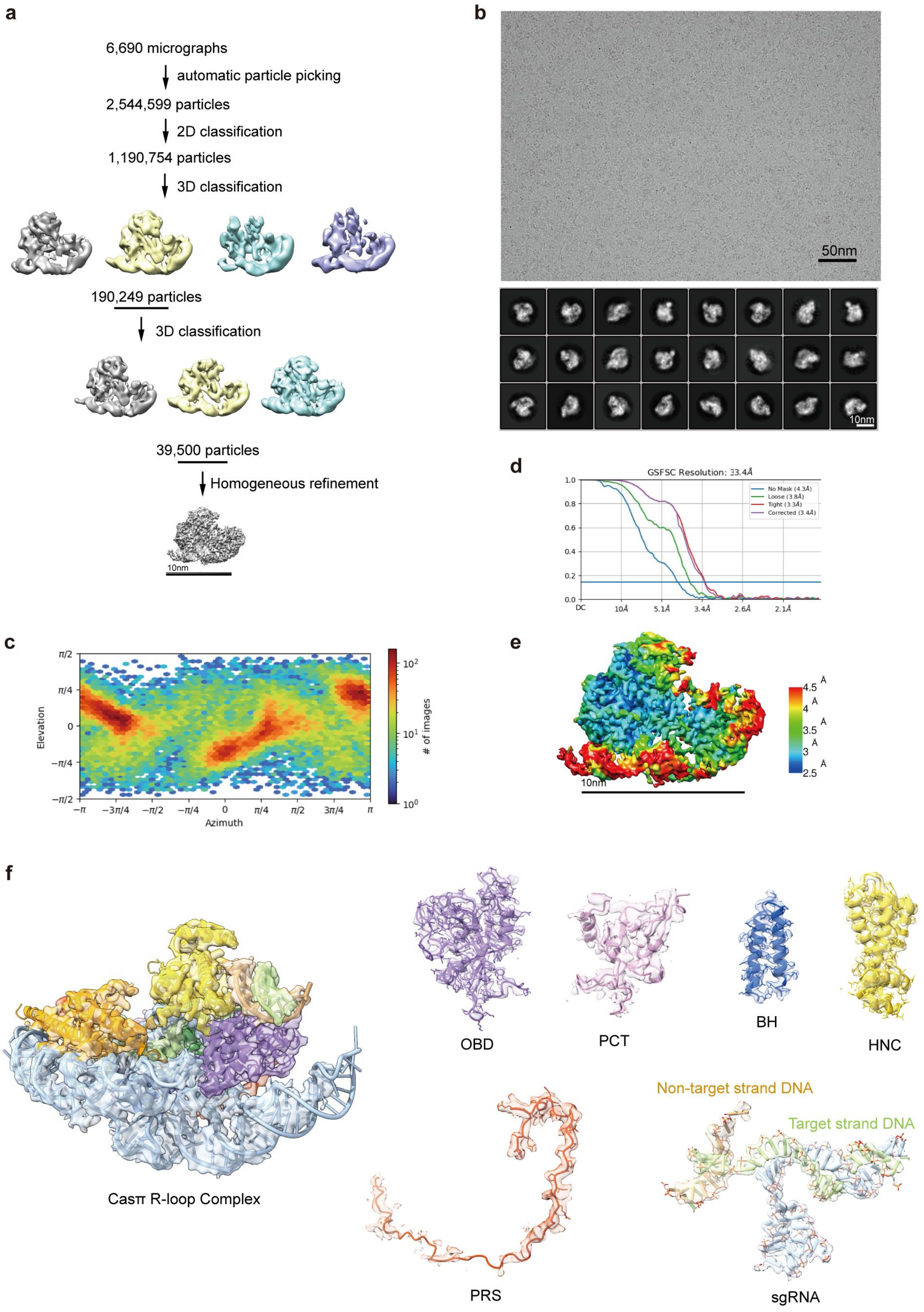


**Figure. S6 Cryo-EM analysis of Casπ-1 R-loop complex.**

(**a**) The workflow for single particle analysis using cryoSPARC.

(**b**) The representative cryo-EM micrograph and 2D class-averages of Casπ-1 R-loop complex.

(**c**) Fourier shell correlation (FSC) curves calculated using two independent half-maps. The resolution for the B-factor corrected final map was 3.4 Å.

(**d**) Euler angle distribution of the refined particles for the final map. Panel C and D are directly adopted from the standard outputs of cryoSPARC.

(**e**) The validation of local resolution. The resolution-range from 2.5 Å to 4.5 Å was shown in the map.

(**f**) The fitting between the cryo-EM map and the atomic model. These maps are shown at the contour threshold of 4.2 times sigma. The overall-fitting and local-details for OBD domain, PCT domain, BH domain, HNC domain, PRS domain, the gRNA and the DNA target were shown.

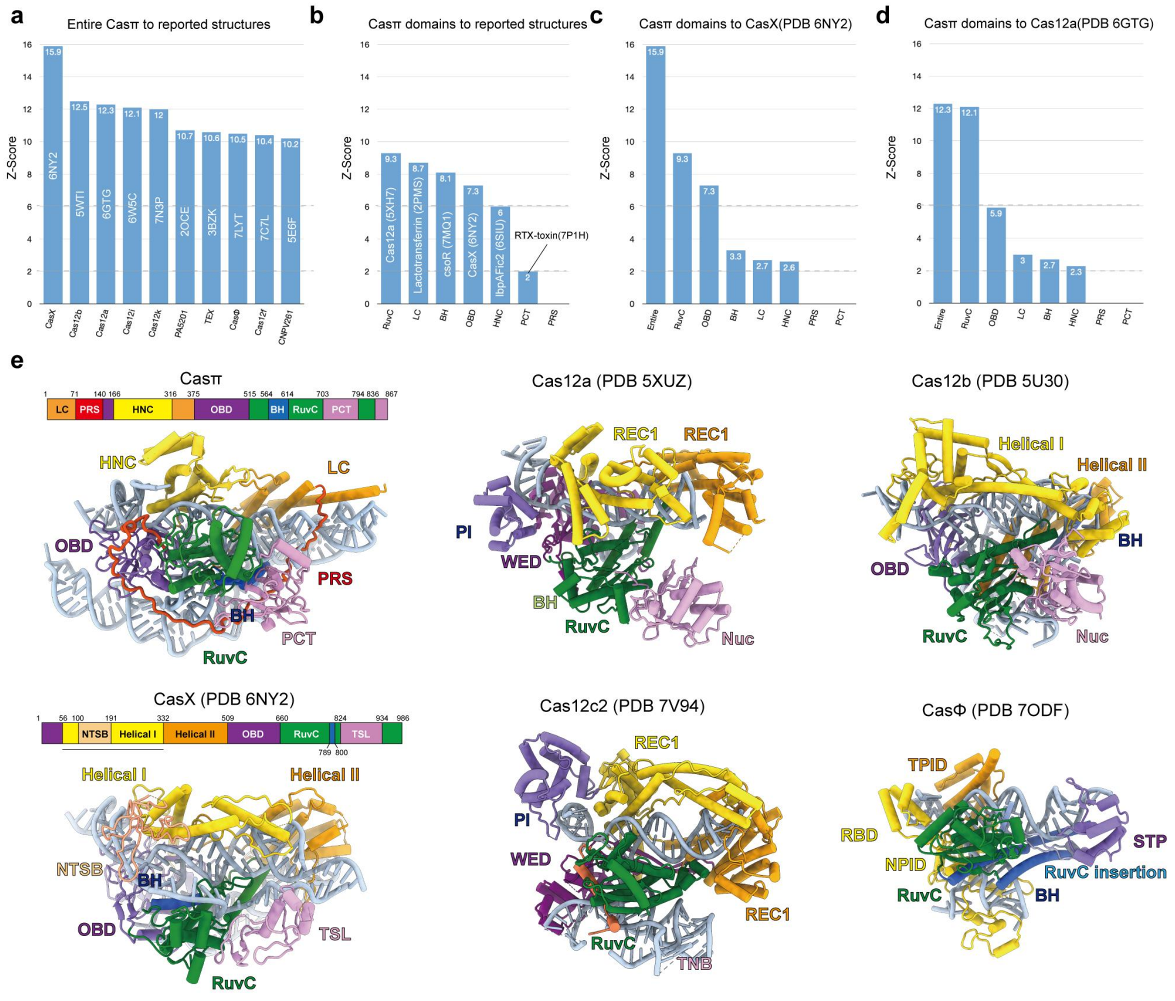

**Figure. S7 Overall structural comparison among representative type V effectors.**

(**a**) The overall structural similarity between Casπ-1 and reported structures. The 10 structures with top Z-scores of similarities to Casπ-1 are listed.

(**b**) Structural similarity of protein domains within Casπ-1 to all reported structures. The reported structure with the highest Z-score of similarity to each Casπ-1 domain was shown.

(**c**) The structural similarity between each Casπ-1 domain and DpbCasX.

(**d**) The structural similarity between each Casπ-1 domain and LbCas12a.

(**e**) The structures of representative type V effectors. All models were shown from the top view with the same size scale. For Casπ and CasX, the domain organization within the primary sequence were shown. The protein domains within each structure are colored and labeled accordingly.

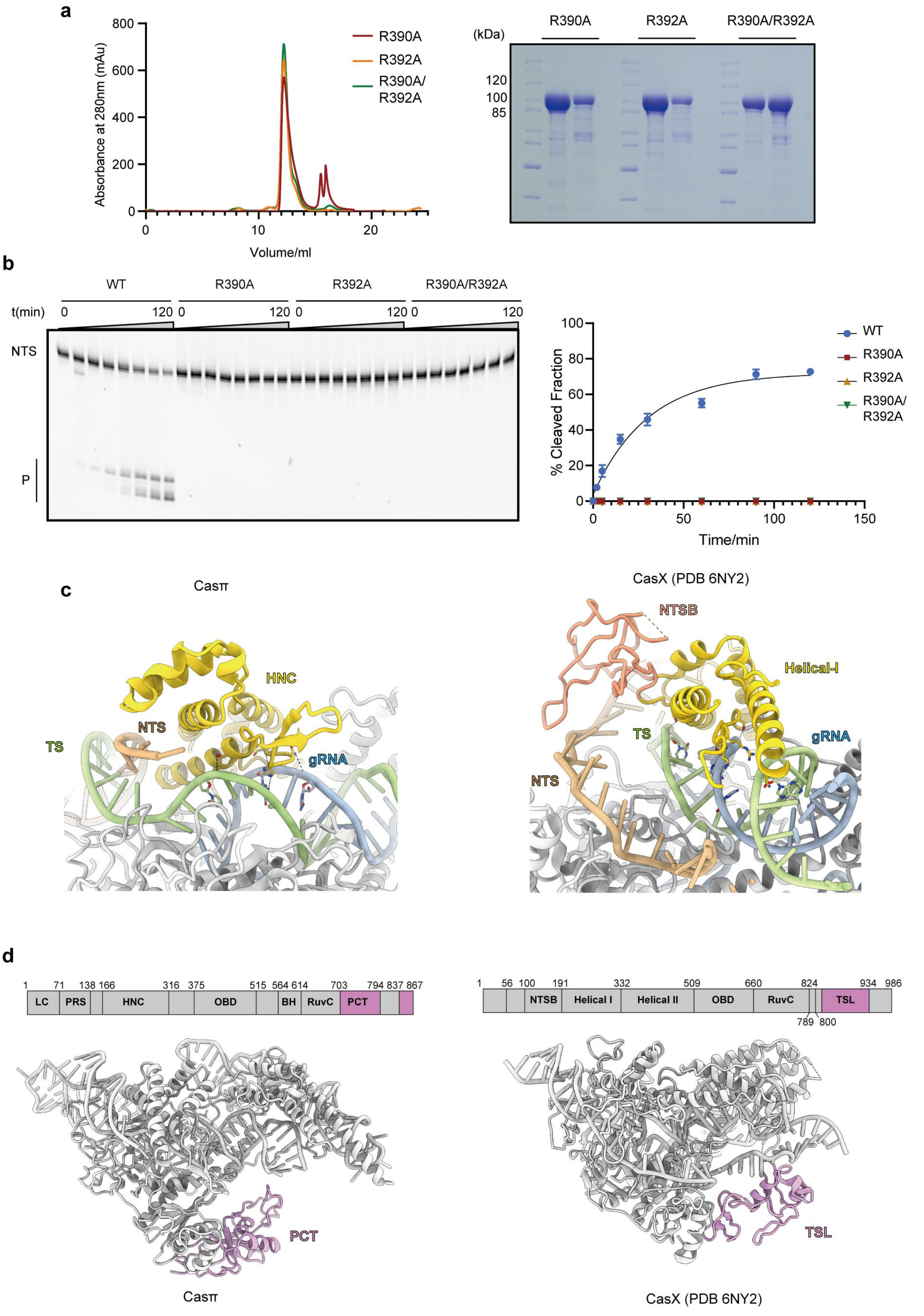
a
Absorbance at 280nm (mAu)
800
600
400
200
0
0
10
20
Volume/ml
R390A
R392A
R390A/
R392A
(kDa)
R390A
R392A
R390A/R392A
120
100
85
b
WT
R390A
R392A
R390A/R392A
t(min)
0
120
0
120
0
120
0
120
NTS
P
% Cleaved Fraction
100
80
60
40
20
0
0
50
100
150
Time/min
WT
R390A
R392A
R390A/
R392A
c
Casπ
CasX (PDB 6NY2)
HNC
NTS
TS
gRNA
NTSB
Helical-I
TS
gRNA
NTS
d
1
71
138
166
316
375
515
564
614
703
794
837
867
LC
PRS
HNC
OBD
BH
RuvC
PCT
1
56
100
191
332
509
660
824
934
986
NTSB
Helical I
Helical II
OBD
RuvC
TSL
789
800
PCT
TSL
Casπ
CasX (PDB 6NY2)

**Figure. S8 Local analysis of OBD, HNC and PCT domain in Casπ protein.**

(**a**) Left, purification of three Casπ mutants (R390A, R392A and R390A/R392A). The SEC curves were aligned referring to the injection volume. Right, the peak fractions were analyzed by SDS-PAGE.

(**b**) Left, *in vitro* dsDNA cleavage by wild type Casπ (WT) and Casπ mutants revealed by denaturing PAGE. Right, the plot of dsDNA substrate cleavage efficiency by WT and mutants of Casπ (n=3 each; means ± SD).

(**c**) Structural comparison between Casπ HNC domain and the NTSB to Helical-I region of CasX.

(**d**) Comparison between Casπ PCT domain and CasX TSL domain. Both the primary and 3D location of each domain in the entire protein were presented.

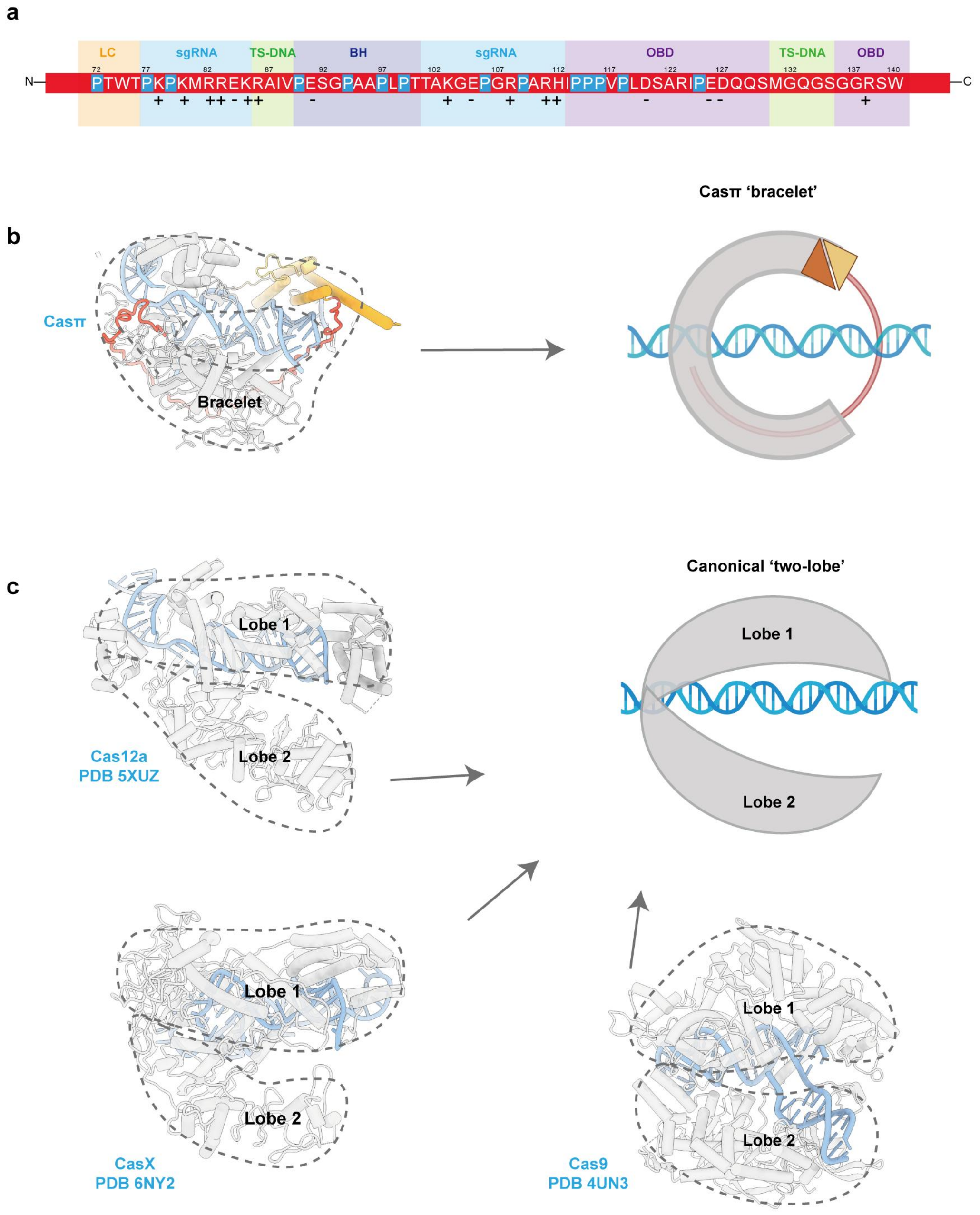


**Figure. S9 The bracelet and two-lobe architectures for Class 2 effectors.**

(**a**) The amino-acid regions within PRS domain that respectively interact with other parts of the Casπ R-loop complex. The other parts were shown by colored rectangles underlying the PRS cartoon.

(**b**) The structure and 'bracelet' model for Casπ. The protein part in the structure was traced with grey dash lines. Protein domains and nucleic acids were colored referring to Figure 6A. The stem elements in sgRNA are hided for clear presentation.

(**c**) The representative structures for reported Class 2 effectors. The real structures for Cas12a, CasX, and Cas9 were shown on the left and bottom panels. The protein part (traced with grey dash lines) was colored in grey and nucleic acid part in cyan. The single strand region of NTS was hided for clear presentation. The cartoon of two-lobe architecture for canonical Class 2 nuclease was shown on the top-right panel.

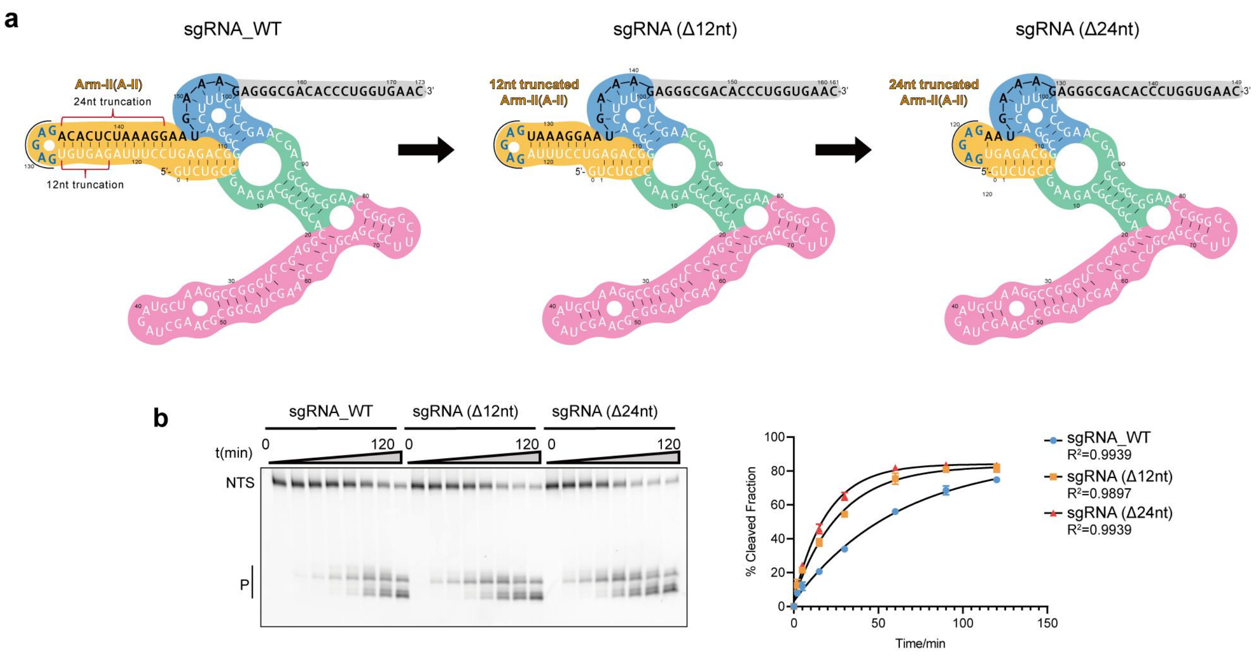


**Figure. S10 sgRNA engineering on the A-II region.**

(**a**) Left, the naive version of the sgRNA (sgRNA_WT); Middle, the secondary structure of sgRNA (Δ12nt) with 12-nt truncation on the A-II region; Right, the secondary structure of sgRNA (Δ24nt) with 24-nt truncation on the A-II region.

(**b**) Left, *In vitro* dsDNA cleavage by Casπ-1 effector using different sgRNAs; Right, the plot of cleavage efficiency (n=3; means ± SD), The rate constant k values for Casπ-1 with sgRNA_WT, sgRNA (Δ12nt) and sgRNA (Δ24nt) are 0.0165, 0.03702 and 0.05125 (fraction/minute).

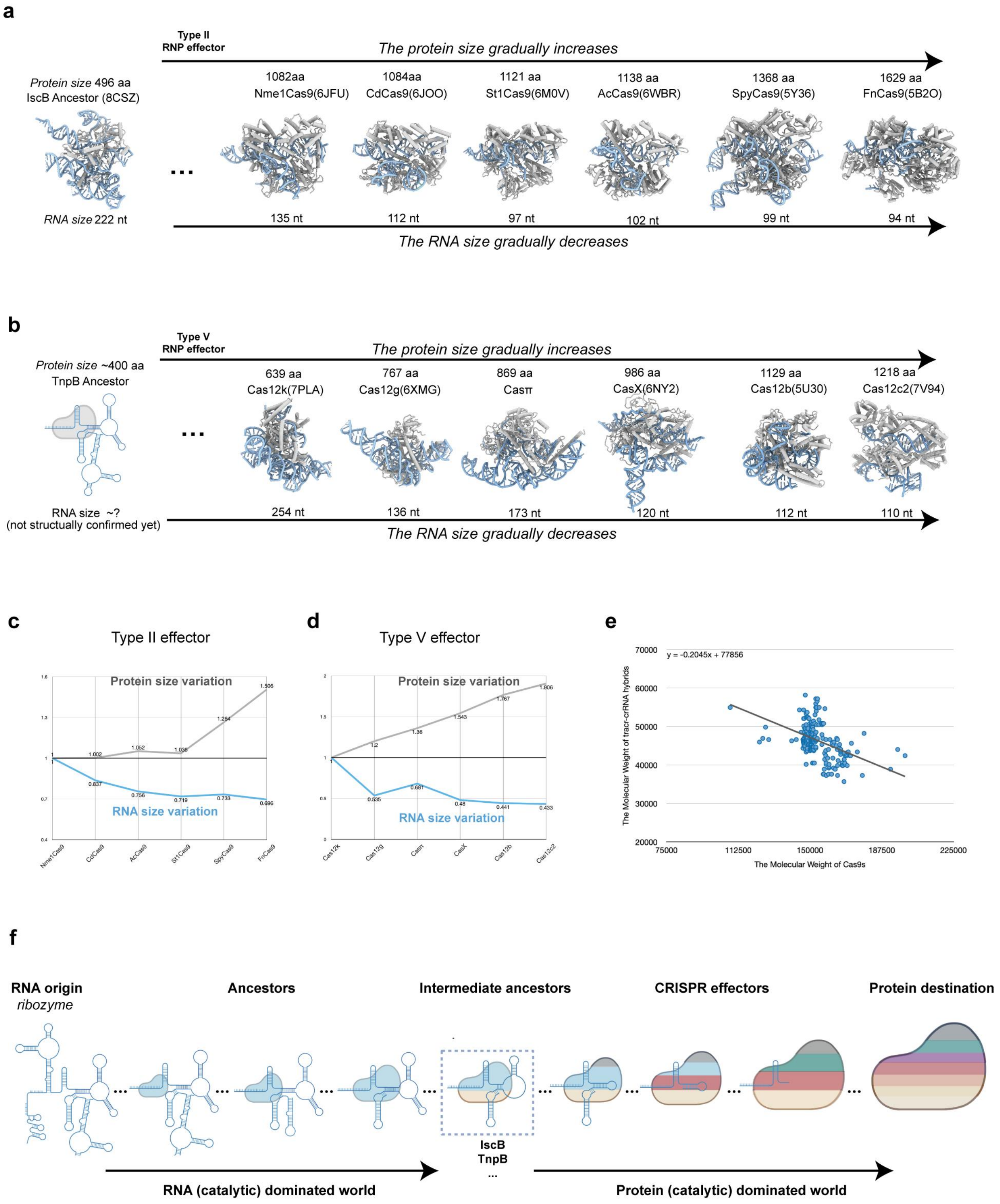


**Figure. S11 Molecular size analysis for the gRNA and protein in CRISPR effectors.**

(**a**) The summary of the RNA and protein size in type II CRISPR effectors comprising of a protein monomer and tracr-crRNA hybrid. The atomic model each effector was shown in the same size scale. The protein part was colored in grey and RNA part in cyan. The number of amino acids (aa) in each protein, the length of each RNA (nt), and the PDB code for each structure were labeled, accordingly. The structure of ancestral IscB effector was shown in the left panel.

(**b**) The summary of the RNA and protein size in type V CRISPR effectors comprising of a protein monomer and tracr-crRNA hybrid. The cartoon of ancestral TnpB effector was presented in the left panel.

(**c**) The RNA- and protein-size variation trend for type II effectors referring to Nme1Cas9 effector (pdb 6JFU). The variation value (y axis) for the protein in each effector equals its protein size divided by Nme1Cas9 protein size. The variation value for the RNA in each effector equals its RNA size divided by NmeCas9 RNA size. The variation value >1 suggests a size increase compared to NmeCas9, and variation value >1 suggests a size decrease.

(**d**) The RNA- and protein-size variation trend for type V effectors referring to Cas12k effector (pdb 7PLA).

(**e**) The correlation analysis of the protein length and the tracr-crRNA molecular weights within 383 bioinformatically identified Cas9 effectors（Cas9's molecular weight is between 100,000-200,000 Da and tracr-crRNA's molecular weight is between 30,000-60,000 Da）. The Y axis represents the tracr-crRNA molecular weights and the X axis indicates the molecular weight of Cas9 proteins in each effector.

(**f**) The hypothetical RNA-protein co-evolution within CRISPR effectors starting from the "RNA world". The size of the RNA and protein is positively correlated to the cartoon size.

**Table S1. The top 100 PAMs ranked by fold-change.**

| PAM preference of Cas$\pi$-1 | | | | | |
|---|---|---|---|---|---|
| 50 mM NaCl | | 150 mM NaCl | | 300 mM NaCl | |
| PAM | Fold change | PAM | Fold change | PAM | Fold change |
| TCCCC | 10.42 | TGCCC | 13.28 | TGCCC | 27.76 |
| GTCCC | 9.99 | TCCCC | 13.17 | TACCC | 26.72 |
| TTCCT | 9.74 | GTCCC | 13.01 | GACCC | 26.44 |
| TGCCC | 9.71 | TACCC | 12.56 | TGCCT | 26.39 |
| GACCT | 9.68 | GACCG | 12.37 | GACCT | 26.02 |
| TACCG | 9.65 | AACCT | 12.08 | TACCT | 25.43 |
| TACCC | 9.64 | AACCC | 12.03 | TTCCC | 25.23 |
| TTCCC | 9.59 | TTCCC | 11.99 | AACCT | 25.22 |
| GACCC | 9.58 | GGCCT | 11.98 | GGCCT | 24.95 |
| GGCCT | 9.56 | TGCCT | 11.98 | TCCCC | 24.90 |
| AGCCC | 9.50 | GACCC | 11.96 | TTCCT | 24.64 |
| GGCCC | 9.48 | GGCCC | 11.93 | AACCC | 24.58 |
| TGCCT | 9.44 | AGCCC | 11.88 | TCCCT | 24.26 |
| AACCC | 9.39 | TACCA | 11.83 | GTCCT | 24.15 |
| TTCCG | 9.35 | ACCCC | 11.76 | GGCCC | 23.99 |
| GTCCT | 9.33 | CTCCC | 11.71 | ATCCC | 23.99 |
| TACCT | 9.29 | ATCCT | 11.63 | AGCCC | 23.85 |
| ACCCT | 9.27 | TACCG | 11.62 | ATCCT | 23.67 |
| TCCCT | 9.27 | TTCCT | 11.59 | GTCCC | 23.58 |
| AACCT | 9.26 | CCCCC | 11.57 | TACCG | 23.26 |
| AACCA | 9.20 | TCCCT | 11.56 | AGCCT | 23.25 |
| GACCA | 9.18 | TGCCA | 11.55 | CTCCC | 22.62 |
| CTCCC | 9.17 | GTCCG | 11.53 | CACCC | 22.27 |
| TGCCG | 9.10 | ATCCC | 11.52 | CGCCT | 21.78 |
| GACCG | 9.09 | GCCCC | 11.51 | CCCCC | 21.40 |
| GTCCG | 9.09 | TACCT | 11.47 | GCCCT | 21.35 |
| GCCCC | 9.06 | TTCCG | 11.40 | GACCG | 21.25 |
| AGCCA | 9.04 | ACCCT | 11.35 | CGCCC | 20.82 |
| ATCCT | 9.04 | GTCCT | 11.34 | ACCCC | 20.80 |
| TACCA | 9.04 | TGCCG | 11.33 | CACCT | 20.61 |
| ATCCC | 8.98 | GACCT | 11.30 | ACCCT | 20.48 |
| ACCCC | 8.95 | AGCCT | 11.12 | AACCG | 20.03 |
| CCCCC | 8.93 | AACCA | 10.93 | TGCCG | 19.88 |
| TTCCA | 8.92 | ATCCG | 10.88 | CTCCT | 19.62 |
| GCCCG | 8.89 | GACCA | 10.88 | GCCCC | 19.41 |

| | | | | | |
|---|---|---|---|---|---|
| AGCCG | 8.87 | CACCC | 10.85 | CCCCT | 19.26 |
| TCCCG | 8.72 | GGCCG | 10.84 | TACCA | 17.08 |
| CACCG | 8.69 | AGCCA | 10.75 | AGCCG | 16.37 |
| TGCCA | 8.66 | AGCCG | 10.75 | ATCCG | 16.32 |
| AACCG | 8.65 | CACCG | 10.71 | TTCCG | 15.98 |
| GGCCA | 8.64 | AACCG | 10.68 | CACCG | 15.79 |
| ATCCG | 8.63 | CCCCT | 10.50 | TGCCA | 15.68 |
| GGCCG | 8.62 | CACCT | 10.35 | CGCCG | 15.53 |
| AGCCT | 8.62 | TTCCA | 10.33 | GGCCG | 15.26 |
| GTCCA | 8.56 | CGCCG | 10.29 | GTCCG | 15.19 |
| ATCCA | 8.56 | CACCA | 10.28 | AACCA | 14.96 |
| CACCC | 8.55 | CGCCC | 10.21 | GACCA | 14.70 |
| TCCCA | 8.39 | TCCCG | 10.16 | CACCA | 13.61 |
| CGCCC | 8.31 | ATCCA | 10.15 | AGCCA | 13.33 |
| CGCCT | 8.30 | GCCCT | 10.07 | TTCCA | 13.08 |
| CACCT | 8.19 | CGCCT | 10.04 | TCCCG | 12.92 |
| ACCCG | 8.18 | CTCCT | 9.93 | CGCCA | 12.90 |
| CACCA | 8.16 | GTCCA | 9.91 | GGCCA | 12.73 |
| CTCCG | 8.11 | CGCCA | 9.90 | CTCCG | 12.71 |
| CTCCT | 8.09 | ACCCG | 9.87 | GTCCA | 12.67 |
| ACCCA | 8.05 | CTCCG | 9.87 | ATCCA | 12.54 |
| CCCCT | 7.94 | TCCCA | 9.83 | ACCCG | 12.29 |
| GCCCA | 7.93 | GGCCA | 9.83 | TCCCA | 12.29 |
| CGCCA | 7.93 | GCCCG | 9.66 | GCCCG | 11.36 |
| CGCCG | 7.91 | GCCCA | 9.52 | ACCCA | 11.17 |
| CTCCA | 7.84 | CCCCG | 9.29 | GCCCA | 11.06 |
| CCCCG | 7.79 | CCCCA | 9.01 | CCCCG | 11.03 |
| CCCCA | 7.47 | ACCCA | 8.89 | CTCCA | 10.60 |
| GCCCT | 7.30 | CTCCA | 8.87 | CCCCA | 10.42 |
| ACCTA | 5.68 | GCCTA | 6.81 | TACTC | 2.81 |
| GCCTA | 5.65 | ACCTA | 6.64 | GTACT | 2.77 |
| TACTC | 5.61 | TACTC | 5.77 | TAACT | 2.64 |
| TCCTA | 5.54 | TAACT | 5.73 | GAACT | 2.50 |
| GTACT | 4.94 | GTACT | 5.47 | GCACT | 2.40 |
| TCACT | 4.82 | GATCT | 5.41 | GATCT | 2.32 |
| TAACT | 4.81 | TTACT | 5.31 | TTACT | 2.06 |
| GAACT | 4.71 | GAACT | 5.24 | TCACT | 2.01 |
| TTACT | 4.69 | TCCTA | 5.12 | ATACT | 1.99 |
| GATCT | 4.61 | GCACT | 5.04 | AAACT | 1.83 |
| ACCGA | 4.59 | ATACT | 4.99 | GTACC | 1.82 |

| | | | | | |
|---|---|---|---|---|---|
| GCCGA | 4.48 | ACACT | 4.79 | ACACT | 1.80 |
| ATACT | 4.38 | GTACC | 4.67 | AACTC | 1.75 |
| GACTC | 4.31 | TACTT | 4.65 | GCCTA | 1.75 |
| AACTC | 4.20 | TCACT | 4.64 | GACTC | 1.74 |
| GTACC | 4.19 | TGTCT | 4.63 | ACCTA | 1.70 |
| ACACT | 4.17 | ACCGA | 4.58 | GATCC | 1.69 |
| CACTC | 4.16 | TGACT | 4.56 | TACTT | 1.59 |
| GCACT | 4.13 | AAACT | 4.54 | GAACC | 1.57 |
| AAACT | 4.12 | GATCC | 4.49 | TAACC | 1.57 |
| TCCGA | 4.08 | GGACT | 4.41 | CACTC | 1.51 |
| TGACT | 4.06 | GAACC | 4.35 | TGACT | 1.50 |
| GATCC | 4.03 | GCCGA | 4.33 | TGTCT | 1.40 |
| TGTCT | 3.99 | AACTC | 4.28 | TGCTC | 1.37 |
| GGACT | 3.94 | GACTC | 4.24 | GCACC | 1.31 |
| CCCTA | 3.92 | TGCTC | 4.09 | GGACT | 1.31 |
| GAACC | 3.90 | TATCT | 4.07 | CAACT | 1.30 |
| TACTT | 3.90 | CACTC | 4.07 | GCTCT | 1.28 |
| TGCTC | 3.89 | TAACC | 4.01 | ATACC | 1.26 |
| TATCT | 3.80 | TCTCT | 3.97 | GTTCT | 1.25 |
| AGACT | 3.77 | GTTCT | 3.88 | GACTT | 1.24 |
| CAACT | 3.76 | GACTT | 3.88 | AAACC | 1.19 |
| TCACC | 3.76 | CAACT | 3.83 | TTACC | 1.19 |
| TGACC | 3.75 | ATACC | 3.81 | TCTCT | 1.18 |
| GCACC | 3.74 | AACTT | 3.74 | TATCT | 1.15 |
| GACTT | 3.71 | GCACC | 3.73 | AACTT | 1.15 |

| **PAM preference of Cas$\pi$-2** | | | | | |
|---|---|---|---|---|---|
| 50 mM NaCl | | 150 mM NaCl | | 300 mM NaCl | |
| PAM | Fold change | PAM | Fold change | PAM | Fold change |
| TACCC | 32.14 | TGCCC | 30.19 | TGCCC | 28.17 |
| GTCCC | 31.80 | GTCCC | 29.61 | TACCC | 27.20 |
| GACCC | 31.73 | TACCC | 29.21 | TTCCC | 27.09 |
| AGCCC | 30.35 | AACCC | 27.09 | TCCCC | 26.79 |
| TGCCC | 30.27 | GACCC | 27.02 | GACCC | 26.39 |
| GGCCC | 30.15 | GGCCC | 26.79 | GGCCC | 25.94 |
| AACCC | 29.18 | AGCCC | 26.45 | GTCCC | 25.16 |
| TTCCC | 28.05 | TTCCC | 26.43 | AGCCC | 24.84 |
| ATCCC | 27.76 | TCCCC | 26.17 | ATCCC | 24.80 |
| CACCC | 26.95 | CTCCC | 26.05 | AACCC | 24.18 |

| | | | | | |
|---|---|---|---|---|---|
| TCCCC | 26.93 | ATCCC | 25.76 | CCCCC | 24.16 |
| CTCCC | 26.85 | CCCCC | 25.28 | GGCCT | 23.99 |
| CGCCC | 26.29 | CACCC | 24.54 | TGCCT | 23.96 |
| CCCCC | 25.99 | AACCT | 24.06 | TCCCT | 23.46 |
| GACCT | 25.84 | TACCT | 23.97 | GCCCT | 23.42 |
| TACCT | 25.13 | TGCCT | 23.71 | GACCT | 23.42 |
| TGCCT | 24.67 | GGCCT | 23.47 | CTCCC | 23.25 |
| GGCCT | 24.40 | ATCCT | 23.22 | TTCCT | 23.13 |
| AACCT | 23.81 | ACCCC | 23.18 | ACCCC | 23.02 |
| GTCCT | 23.67 | GACCT | 23.00 | TACCT | 22.96 |
| GCCCC | 23.43 | GTCCT | 22.70 | GTCCT | 22.59 |
| ACCCC | 23.28 | CGCCC | 22.47 | CACCC | 22.55 |
| TTCCT | 23.09 | AGCCT | 22.38 | AACCT | 22.46 |
| ATCCT | 22.39 | GCCCC | 22.26 | GCCCC | 22.17 |
| AGCCT | 21.85 | TTCCT | 22.15 | AGCCT | 21.99 |
| CACCT | 21.50 | ACCCT | 20.80 | ATCCT | 21.84 |
| CGCCT | 21.26 | CACCT | 20.66 | CGCCC | 21.56 |
| ACCCT | 20.11 | CCCCT | 20.41 | ACCCT | 21.28 |
| CTCCT | 20.06 | TCCCT | 20.33 | CGCCT | 20.37 |
| TCCCT | 19.78 | CTCCT | 19.86 | CCCCT | 20.33 |
| CCCCT | 19.51 | CGCCT | 19.59 | CTCCT | 19.33 |
| GCCCT | 18.63 | GCCCT | 18.85 | CACCT | 18.96 |
| GACCA | 12.69 | TACCA | 15.07 | TACCA | 15.86 |
| TACCA | 12.59 | GACCA | 13.02 | GACCA | 13.87 |
| AGCCA | 11.05 | AGCCA | 12.51 | AGCCA | 13.47 |
| AACCA | 10.35 | AACCA | 12.32 | TGCCA | 13.09 |
| CACCA | 9.97 | TGCCA | 11.92 | AACCA | 12.75 |
| CGCCA | 9.77 | CGCCA | 11.58 | CGCCA | 12.59 |
| TACTC | 9.59 | CACCA | 11.54 | CACCA | 11.98 |
| GGCCA | 9.58 | TACTC | 10.97 | TACTC | 11.52 |
| TACCG | 9.57 | TACCG | 10.82 | GGCCA | 11.30 |
| TGCCA | 9.29 | GGCCA | 10.75 | TACCG | 10.74 |
| GACCG | 7.77 | GACCG | 9.88 | GTCCA | 9.36 |
| GTCCA | 7.39 | ATCCA | 8.81 | ATCCA | 9.36 |
| GACTC | 7.29 | GTCCA | 8.67 | GACCG | 7.97 |
| ATCCA | 7.27 | AGCCG | 8.25 | CACTC | 7.85 |
| AGCCG | 6.93 | GACTC | 8.06 | GACTC | 7.79 |
| CACTC | 6.83 | TGCTC | 8.04 | TGCTC | 7.78 |
| CTCCA | 6.55 | CACTC | 7.80 | TTCCA | 7.74 |
| CCCCA | 6.26 | TGCCG | 7.77 | CTCCA | 7.53 |

| | | | | | |
|---|---|---|---|---|---|
| TGCTC | 6.23 | CCCCA | 7.50 | AGCCG | 7.38 |
| GGCCG | 6.14 | CGCTC | 7.49 | TGCCG | 7.35 |
| TGCCG | 6.14 | CTCCA | 7.46 | CGCTC | 7.32 |
| CGCTC | 6.13 | CACCG | 7.44 | CCCCA | 7.25 |
| CACCG | 6.00 | GGCCG | 7.41 | AACCG | 6.98 |
| TTCCA | 5.92 | AACCG | 7.38 | AACTC | 6.74 |
| AACCG | 5.86 | TTCCA | 7.16 | GGCCG | 6.51 |
| ACCCA | 5.55 | GTCCG | 6.82 | TCCCA | 6.15 |
| GTCCG | 5.53 | CGCCG | 6.82 | CGCCG | 6.10 |
| AACTC | 5.50 | AACTC | 6.79 | AGCTC | 6.08 |
| CGCCG | 5.45 | ATCCG | 6.58 | ACCCA | 6.06 |
| GCCCA | 5.44 | AGCTC | 6.40 | ATCCG | 5.96 |
| AGCTC | 5.23 | GCCCA | 6.37 | CACCG | 5.93 |
| ATCCG | 5.23 | ACCCA | 6.23 | GCCCA | 5.86 |
| TCCCA | 5.06 | TCCCA | 6.11 | GTCTC | 5.81 |
| GGCTC | 4.80 | GGCTC | 5.75 | ATCTC | 5.68 |
| GTCTC | 4.59 | GTCTC | 5.66 | GTCCG | 5.55 |
| ATCTC | 4.09 | TTCCG | 5.05 | GGCTC | 5.37 |
| TTCCG | 4.04 | CTCCG | 4.99 | GTACC | 4.56 |
| CTCCG | 4.03 | ATCTC | 4.81 | TTCCG | 4.50 |
| CTCTC | 3.69 | CTCTC | 4.59 | TTCTC | 4.42 |
| GTACC | 3.60 | TTCTC | 4.23 | CTCTC | 4.35 |
| TTCTC | 3.56 | TACTT | 4.08 | CTCCG | 4.19 |
| CCCCG | 3.43 | GTACC | 4.02 | TGACC | 4.19 |
| TGACC | 3.29 | CCCCG | 3.79 | TGTCC | 3.62 |
| TCCTC | 2.94 | TGACC | 3.66 | TCCTC | 3.56 |
| CCCTC | 2.94 | TCCTC | 3.58 | TACTT | 3.49 |
| TGTCC | 2.81 | TGTCC | 3.44 | CCCCG | 3.47 |
| TCCCG | 2.68 | ACCCG | 3.39 | TCCCG | 3.19 |
| TACTT | 2.65 | CCCTC | 3.28 | CCCTC | 3.14 |
| ACCCG | 2.63 | TGCTT | 3.17 | ACCCG | 3.14 |
| TTACC | 2.57 | TCCCG | 3.10 | TTACC | 3.02 |
| GCCCG | 2.49 | GTACT | 3.05 | GTACT | 2.97 |
| GTACT | 2.47 | GCCTC | 2.92 | TGCTT | 2.89 |
| GCCTC | 2.36 | ACCTC | 2.82 | GCCTC | 2.87 |
| TGCTT | 2.35 | CGCTT | 2.73 | CGACC | 2.85 |
| CGTCC | 2.31 | GACTT | 2.72 | ACCTC | 2.79 |
| ACCTC | 2.26 | AGCTT | 2.71 | GTTCC | 2.72 |
| GTTCC | 2.25 | GCCCG | 2.68 | CGTCC | 2.51 |
| ATACC | 2.18 | CGTCC | 2.63 | ATACC | 2.47 |

| | | | | | |
|---|---|---|---|---|---|
| GACTT | 2.12 | TTACC | 2.63 | GTCTT | 2.40 |
| AGCTT | 2.05 | GTTCC | 2.62 | CGCTT | 2.39 |
| CGACC | 2.01 | GTCTT | 2.62 | GCACC | 2.39 |
| CGCTT | 1.97 | CGACC | 2.52 | GCCCG | 2.37 |
| TCACC | 1.91 | CACTT | 2.42 | TTACT | 2.29 |
| GCACC | 1.85 | GGCTT | 2.42 | TCACC | 2.26 |
| GCTCC | 1.84 | TTACT | 2.32 | GGACC | 2.26 |
| TTACT | 1.84 | AACTT | 2.22 | GACTT | 2.22 |
| GTCTT | 1.81 | ATACC | 2.21 | GGTCC | 2.19 |
| TCTCC | 1.81 | ATCTT | 2.20 | AGCTT | 2.17 |

**Table. S2. Plasmids generated in this study.**

| ID | Assay | Features | Selection marker |
|---|---|---|---|
| pAS001 | *In vitro* plasmids cleavage | pUC19-backbone derived plasmids containing TTGGCAGC PAM and a complementary protospacer region. | Ampicillin |
| pAS002 | *In vitro* plasmids cleavage | pUC19-backbone derived plasmids containing CCTGCCCC PAM and a complementary protospacer region. | Ampicillin |
| pAS003 | *In vitro* plasmids cleavage, cleavage sites determination | pUC19-backbone derived plasmids containing TGCCC PAM and a complementary protospacer region. | Ampicillin |
| pAS004 | PAM-depletion | pUC19-backbone derived plasmids containing five random nucleotides PAM (NNNNN) and a complementary protospacer region. | Ampicillin |
| pET28a-His-SUMO-Casπ-1 | Protein purification | pET28a-backbone derived plasmids containing N-terminal hexa-histidine and a SUMO tagged *Casπ-1* in MCS. | Kanamycin |
| pET28a-His-SUMO-Casπ-2 | Protein purification | pET28a-backbone derived plasmids containing N-terminal hexa-histidine and a SUMO tagged *Casπ-2* in MCS. | Kanamycin |
| pET28a-His-SUMO-Casπ-1 (D537A, E643A) | Protein purification | pET28a-backbone derived plasmids containing N-terminal hexa-histidine and a SUMO tagged d*Casπ-1* (D537A, E643A) in MCS. | Kanamycin |
| p-0380 | Cell genome editing | Plasmid contains *Homo sapiens* codon optimized *Casπ-1* driven by CMV promoter. Both N-/ C-termini of Casπ-1 fused to a SV40 NLS sequences, C-terminal containing a 2 × FLAG tag and linked to PuroR via the P2A peptide sequence. U6-promoter activating the guide transcription and a repeat-spacer unit terminated by a poly-T sequence. *Sap*I-GG stuffer spacer. | Ampicillin (*E. coli*) Puromycin (*H. sapiens*) |
| p-0381 | Cell genome editing | Plasmid contains *H. sapiens* codon optimized *Casπ-2* driven by CMV promoter. Both N-/ C-termini of Casπ-2 fused to a SV40 NLS sequences, C-terminal containing a 2 × FLAG tag and linked to PuroR via the P2A peptide sequence. U6-promoter activating the guide transcription and a repeat-spacer unit terminated by a poly-T sequence. *Sap*I-GG stuffer spacer. | Ampicillin (*E. coli*) Puromycin (*H. sapiens*) |
| p-0382 | Cell genome editing | Plasmid contains *H. sapiens* codon optimized Lb*Cas12a* driven by CMV promoter. Both N-/ C-termini of Cas12a fused to a SV40 NLS sequences, C-terminal containing a 2 × FLAG tag and linked to PuroR via the P2A peptide sequence. U6-promoter activating the guide transcription and a repeat-spacer unit terminated by a poly-T sequence. *Sap*I-GG stuffer spacer. | Ampicillin (*E. coli*) Puromycin (*H. sapiens*) |
| p-0383 | Cell genome editing | Plasmid contains *H. sapiens* codon optimized Spy*Cas9* driven by CMV promoter. Both N-/ C-termini of Cas9 fused to a SV40 NLS sequences, C-terminal containing a 2 × FLAG tag and | Ampicillin (*E. coli*) Puromycin (*H.* |

| | | | |
|---|---|---|---|
| | | linked to PuroR via the P2A peptide sequence. U6-promoter activating the guide transcription and a repeat-spacer unit terminated by a poly-T sequence. *Sap*I-GG stuffer spacer. | *sapiens*) |
| p-0384 | Cell genome editing | Plasmid contains *H. sapiens* codon optimized *Casπ-1* driven by CMV promoter. Both N-/ C-termini of Casπ-1 fused to a SV40 NLS sequences, C-terminal containing a 2 × FLAG tag and linked to PuroR via the P2A peptide sequence. U6-promoter activating the guide transcription and a repeat-spacer unit terminated by a poly-T sequence. MYH8 exon region targeting guide #π-sg1 spacer. | Ampicillin (*E. coli*) Puromycin (*H. sapiens*) |
| p-0385 | Cell genome editing | Plasmid contains *H. sapiens* codon optimized *Casπ-1* driven by CMV promoter. Both N-/ C-termini of Casπ-1 fused to a SV40 NLS sequences, C-terminal containing a 2 × FLAG tag and linked to PuroR via the P2A peptide sequence. U6-promoter activating the guide transcription and a repeat-spacer unit terminated by a poly-T sequence. MYH8 exon region targeting guide #π-sg2 spacer. | Ampicillin (*E. coli*) Puromycin (*H. sapiens*) |
| p-0386 | Cell genome editing | Plasmid contains *H. sapiens* codon optimized *Casπ-1* driven by CMV promoter. Both N-/ C-termini of Casπ-1 fused to a SV40 NLS sequences, C-terminal containing a 2 × FLAG tag and linked to PuroR via the P2A peptide sequence. U6-promoter activating the guide transcription and a repeat-spacer unit terminated by a poly-T sequence. MYH8 exon region targeting guide #π-sg3 spacer. | Ampicillin (*E. coli*) Puromycin (*H. sapiens*) |
| p-0387 | Cell genome editing | Plasmid contains *H. sapiens* codon optimized *Casπ-1* driven by CMV promoter. Both N-/ C-termini of Casπ-1 fused to a SV40 NLS sequences, C-terminal containing a 2 × FLAG tag and linked to PuroR via the P2A peptide sequence. U6-promoter activating the guide transcription and a repeat-spacer unit terminated by a poly-T sequence. MYH8 exon region targeting guide #π-sg4 spacer. | Ampicillin (*E. coli*) Puromycin (*H. sapiens*) |
| p-0388 | Cell genome editing | Plasmid contains *H. sapiens* codon optimized *Casπ-1* driven by CMV promoter. Both N-/ C-termini of Casπ-1 fused to a SV40 NLS sequences, C-terminal containing a 2 × FLAG tag and linked to PuroR via the P2A peptide sequence. U6-promoter activating the guide transcription and a repeat-spacer unit terminated by a poly-T sequence. MYH8 exon region targeting guide #π-sg5 spacer. | Ampicillin (*E. coli*) Puromycin (*H. sapiens*) |
| p-0389 | Cell genome editing | Plasmid contains *H. sapiens* codon optimized *Casπ-2* driven by CMV promoter. Both N-/ C-termini of Casπ-2 fused to a SV40 NLS sequences, C-terminal containing a 2 × FLAG tag and linked to PuroR via the P2A peptide sequence. U6-promoter activating the guide transcription and a repeat-spacer unit terminated by a poly-T sequence. MYH8 exon region targeting guide #π-sg1 spacer. | Ampicillin (*E. coli*) Puromycin (*H. sapiens*) |
| p-0390 | Cell genome editing | Plasmid contains *H. sapiens* codon optimized *Casπ-2* driven by CMV promoter. Both N-/ C-termini of Casπ-2 fused to a SV40 NLS sequences, C-terminal containing a 2 × FLAG tag and linked to PuroR via the P2A peptide sequence. U6-promoter activating the guide transcription and a repeat-spacer unit terminated by a poly-T sequence. MYH8 exon region targeting guide #π-sg2 spacer. | Ampicillin (*E. coli*) Puromycin (*H. sapiens*) |
| p-0391 | Cell genome editing | Plasmid contains *H. sapiens* codon optimized *Casπ-2* driven by CMV promoter. Both N-/ C-termini of Casπ-2 fused to a SV40 | Ampicillin (*E. coli*) |

| | | | |
|---|---|---|---|
| | | NLS sequences, C-terminal containing a 2 × FLAG tag and linked to PuroR via the P2A peptide sequence. U6-promoter activating the guide transcription and a repeat-spacer unit terminated by a poly-T sequence. MYH8 exon region targeting guide #π-sg3 spacer. | Puromycin (*H. sapiens*) |
| p-0392 | Cell genome editing | Plasmid contains *H. sapiens* codon optimized *Casπ-2* driven by CMV promoter. Both N-/ C-termini of Casπ-2 fused to a SV40 NLS sequences, C-terminal containing a 2 × FLAG tag and linked to PuroR via the P2A peptide sequence. U6-promoter activating the guide transcription and a repeat-spacer unit terminated by a poly-T sequence. MYH8 exon region targeting guide #π-sg4 spacer. | Ampicillin (*E. coli*) Puromycin (*H. sapiens*) |
| p-0393 | Cell genome editing | Plasmid contains *H. sapiens* codon optimized *Casπ-2* driven by CMV promoter. Both N-/ C-termini of Casπ-2 fused to a SV40 NLS sequences, C-terminal containing a 2 × FLAG tag and linked to PuroR via the P2A peptide sequence. U6-promoter activating the guide transcription and a repeat-spacer unit terminated by a poly-T sequence. MYH8 exon region targeting guide #π-sg5 spacer. | Ampicillin (*E. coli*) Puromycin (*H. sapiens*) |
| p-0394 | Cell genome editing | Plasmid contains *H. sapiens* codon optimized Lb*Cas12a* driven by CMV promoter. Both N-/ C-termini of Cas12a fused to a SV40 NLS sequences, C-terminal containing a 2 × FLAG tag and linked to PuroR via the P2A peptide sequence. U6-promoter activating the guide transcription and a repeat-spacer unit terminated by a poly-T sequence. MYH8 exon region targeting guide #12a-sg1 spacer. | Ampicillin (*E. coli*) Puromycin (*H. sapiens*) |
| p-0395 | Cell genome editing | Plasmid contains *H. sapiens* codon optimized Lb*Cas12a* driven by CMV promoter. Both N-/ C-termini of Cas12a fused to a SV40 NLS sequences, C-terminal containing a 2 × FLAG tag and linked to PuroR via the P2A peptide sequence. U6-promoter activating the guide transcription and a repeat-spacer unit terminated by a poly-T sequence. MYH8 exon region targeting guide #12a-sg2 spacer. | Ampicillin (*E. coli*) Puromycin (*H. sapiens*) |
| p-0396 | Cell genome editing | Plasmid contains *H. sapiens* codon optimized Lb*Cas12a* driven by CMV promoter. Both N-/ C-termini of Cas12a fused to a SV40 NLS sequences, C-terminal containing a 2 × FLAG tag and linked to PuroR via the P2A peptide sequence. U6-promoter activating the guide transcription and a repeat-spacer unit terminated by a poly-T sequence. MYH8 exon region targeting guide #12a-sg3 spacer. | Ampicillin (*E. coli*) Puromycin (*H. sapiens*) |
| p-0397 | Cell genome editing | Plasmid contains *H. sapiens* codon optimized Lb*Cas12a* driven by CMV promoter. Both N-/ C-termini of Cas12a fused to a SV40 NLS sequences, C-terminal containing a 2 × FLAG tag and linked to PuroR via the P2A peptide sequence. U6-promoter activating the guide transcription and a repeat-spacer unit terminated by a poly-T sequence. MYH8 exon region targeting guide #12a-sg4 spacer. | Ampicillin (*E. coli*) Puromycin (*H. sapiens*) |
| p-0398 | Cell genome editing | Plasmid contains *H. sapiens* codon optimized Lb*Cas12a* driven by CMV promoter. Both N-/ C-termini of Cas12a fused to a SV40 NLS sequences, C-terminal containing a 2 × FLAG tag and linked to PuroR via the P2A peptide sequence. U6-promoter activating the guide transcription and a repeat-spacer unit terminated by a poly-T sequence. MYH8 exon region targeting guide #12a-sg5 spacer. | Ampicillin (*E. coli*) Puromycin (*H. sapiens*) |

| | | | |
|---|---|---|---|
| p-0399 | Cell genome editing | Plasmid contains *H. sapiens* codon optimized Spy*Cas9* driven by CMV promoter. Both N-/ C-termini of Cas9 fused to a SV40 NLS sequences, C-terminal containing a 2 × FLAG tag and linked to PuroR via the P2A peptide sequence. U6-promoter activating the guide transcription and a repeat-spacer unit terminated by a poly-T sequence. MYH8 exon region targeting guide #Cas9-sg1 spacer. | Ampicillin (*E. coli*) Puromycin (*H. sapiens*) |
| p-0400 | Cell genome editing | Plasmid contains *H. sapiens* codon optimized Spy*Cas9* driven by CMV promoter. Both N-/ C-termini of Cas9 fused to a SV40 NLS sequences, C-terminal containing a 2 × FLAG tag and linked to PuroR via the P2A peptide sequence. U6-promoter activating the guide transcription and a repeat-spacer unit terminated by a poly-T sequence. MYH8 exon region targeting guide #Cas9-sg2 spacer. | Ampicillin (*E. coli*) Puromycin (*H. sapiens*) |
| p-0401 | Cell genome editing | Plasmid contains *H. sapiens* codon optimized Spy*Cas9* driven by CMV promoter. Both N-/ C-termini of Cas9 fused to a SV40 NLS sequences, C-terminal containing a 2 × FLAG tag and linked to PuroR via the P2A peptide sequence. U6-promoter activating the guide transcription and a repeat-spacer unit terminated by a poly-T sequence. MYH8 exon region targeting guide #Cas9-sg3 spacer. | Ampicillin (*E. coli*) Puromycin (*H. sapiens*) |
| p-0402 | Cell genome editing | Plasmid contains *H. sapiens* codon optimized Spy*Cas9* driven by CMV promoter. Both N-/ C-termini of Cas9 fused to a SV40 NLS sequences, C-terminal containing a 2 × FLAG tag and linked to PuroR via the P2A peptide sequence. U6-promoter activating the guide transcription and a repeat-spacer unit terminated by a poly-T sequence. MYH8 exon region targeting guide #Cas9-sg4 spacer. | Ampicillin (*E. coli*) Puromycin (*H. sapiens*) |
| p-0403 | Cell genome editing | Plasmid contains *H. sapiens* codon optimized Spy*Cas9* driven by CMV promoter. Both N-/ C-termini of Cas9 fused to a SV40 NLS sequences, C-terminal containing a 2 × FLAG tag and linked to PuroR via the P2A peptide sequence. U6-promoter activating the guide transcription and a repeat-spacer unit terminated by a poly-T sequence. MYH8 exon region targeting guide #Cas9-sg5 spacer.. | Ampicillin (*E. coli*) Puromycin (*H. sapiens*) |
| p-0404 | Cell genome editing | Lentiviral expression MCPyV-backbone plasmid containing the *MYH8* exon (270 bp) from *M. musculus* with 32 bp random flanking sequence linked to the start codon of *EGFP* under the control of CMV promoter. | Ampicillin (*E. coli*) Puromycin (*H. sapiens*) |
| p-0405 | Bacteria plasmid interference assay | Plasmid contains *Casπ-1* under the control of Trc promoter. Casπ-1 is N-terminally linked to a 6×His tag. Locus for guide expression is controlled by J23119-promoter and constitutes a repeat-spacer unit terminated by a T7 terminator. *Sap*I-GG stuffer spacer. | Streptomycin |
| p-0406 | Bacteria plasmid interference assay | Plasmid contains *Casπ-1* under the control of Trc promoter. Casπ-1 is N-terminally linked to a 6×His tag. Locus for guide expression is controlled by J23119-promoter and constitutes a repeat-spacer unit terminated by a T7 terminator. The ccdB gene targeting guide #ccdB-T spacer. | Streptomycin |
| p-0407 | Bacteria plasmid interference assay | Plasmid contains *Casπ-2* under the control of Trc promoter. Casπ-1 is N-terminally linked to a 6×His tag. Locus for guide expression is controlled by J23119-promoter and constitutes a repeat-spacer unit terminated by a T7 terminator. *Sap*I-GG stuffer spacer. | Streptomycin |

| | | | |
|---|---|---|---|
| p-0408 | Bacteria plasmid interference assay | Plasmid contains *Casπ-2* under the control of Trc promoter. Casπ-1 is N-terminally linked to a 6×His tag. Locus for guide expression is controlled by J23119-promoter and constitutes a repeat-spacer unit terminated by a T7 terminator. The ccdB gene targeting guide #ccdB-T spacer. | Streptomycin |
| p11-LacY-wtx1 | Bacteria plasmid interference assay | Inducible pBAD promoter activating toxin ccdB expression (addgene #69056) | Ampicillin |
| p-0409 | Cell genome editing | Plasmid contains *H. sapiens* codon optimized *Casπ-1* driven by CMV promoter. Both N-/ C-termini of Casπ-1 fused to a SV40 NLS sequences, C-terminal containing a 2 × FLAG tag and linked to PuroR via the P2A peptide sequence. U6-promoter activating the guide transcription and a repeat-spacer unit terminated by a poly-T sequence. MYH8 exon region targeting guide #π-*B2M*-1 spacer. | Ampicillin (*E. coli*) Puromycin (*H. sapiens*) |
| p-0410 | Cell genome editing | Plasmid contains *H. sapiens* codon optimized *Casπ-1* driven by CMV promoter. Both N-/ C-termini of Casπ-1 fused to a SV40 NLS sequences, C-terminal containing a 2 × FLAG tag and linked to PuroR via the P2A peptide sequence. U6-promoter activating the guide transcription and a repeat-spacer unit terminated by a poly-T sequence. MYH8 exon region targeting guide #π-*B2M*-2 spacer. | Ampicillin (*E. coli*) Puromycin (*H. sapiens*) |
| p-0411 | Cell genome editing | Plasmid contains *H. sapiens* codon optimized *Casπ-1* driven by CMV promoter. Both N-/ C-termini of Casπ-1 fused to a SV40 NLS sequences, C-terminal containing a 2 × FLAG tag and linked to PuroR via the P2A peptide sequence. U6-promoter activating the guide transcription and a repeat-spacer unit terminated by a poly-T sequence. MYH8 exon region targeting guide #π-*TP53*-1spacer. | Ampicillin (*E. coli*) Puromycin (*H. sapiens*) |
| p-0412 | Cell genome editing | Plasmid contains *H. sapiens* codon optimized *Casπ-2* driven by CMV promoter. Both N-/ C-termini of Casπ-2 fused to a SV40 NLS sequences, C-terminal containing a 2 × FLAG tag and linked to PuroR via the P2A peptide sequence. U6-promoter activating the guide transcription and a repeat-spacer unit terminated by a poly-T sequence. MYH8 exon region targeting guide #π-*B2M*-1 spacer. | Ampicillin (*E. coli*) Puromycin (*H. sapiens*) |
| p-0413 | Cell genome editing | Plasmid contains *H. sapiens* codon optimized *Casπ-2* driven by CMV promoter. Both N-/ C-termini of Casπ-2 fused to a SV40 NLS sequences, C-terminal containing a 2 × FLAG tag and linked to PuroR via the P2A peptide sequence. U6-promoter activating the guide transcription and a repeat-spacer unit terminated by a poly-T sequence. MYH8 exon region targeting guide #π-*B2M*-2 spacer. | Ampicillin (*E. coli*) Puromycin (*H. sapiens*) |
| p-0414 | Cell genome editing | Plasmid contains *H. sapiens* codon optimized *Casπ-1* driven by CMV promoter. Both N-/ C-termini of Casπ-2 fused to a SV40 NLS sequences, C-terminal containing a 2 × FLAG tag and linked to PuroR via the P2A peptide sequence. U6-promoter activating the guide transcription and a repeat-spacer unit terminated by a poly-T sequence. MYH8 exon region targeting guide #π-*TP53*-1 spacer. | Ampicillin (*E. coli*) Puromycin (*H. sapiens*) |

**Table. S3. Oligonucleotides and target sequences used in this study**

| ID # | Assay | Description | Sequences (5'-3') |
|---|---|---|---|
| oligo-001 | Cy5-labeled dsDNA cleavage of Casπ | Forward PCR primer for cy5-labeled dsDNA target amplification | Cy5-CGGTACCCGGGGATCC |
| oligo-002 | Cy5-labeled dsDNA cleavage of Casπ | Reverse PCR primes for cy5-labeled dsDNA target amplification of Casπ | HO-TATGACCATGATTAGGCCAAGCTTGTTCACCAGGGTGTCGCCCTGGGCAGGATCCCCGGGTACCG |
| oligo-003 | Cy5-labeled dsDNA cleavage, cleavage sites determination and *trans*-cleavage assay of Casπ | DNA non-target strand containing a TGCCC PAM (blue) and a protospacer region (red) for Casπ cleavage assay and cleavage sites determination | $NH_2$-CGGTACCCGGGGATCCTGCCCAGGGCGACACCCTGGTGAACAAGCTTGGCCTAATCATGGTCATA |
| oligo-004 | Cy5-labeled dsDNA cleavage, cleavage sites determination and *trans*-cleavage assay of Casπ | Non-labelled complementary strand of oligo-003 and oligo-006 | OH-TATGACCATGATTAGGCCAAGCTTGTTCACCAGGGTGTCGCCCTGGGCAGGATCCCCGGGTACCG |
| oligo-005 | Cy5-labeled dsDNA cleavage, ssDNA cleavage, cleavage sites determination and *trans*-cleavage assay of Casπ | DNA target strand for Casπ cleavage assay and cleavage sites determination of Casπ | $NH_2$-TATGACCATGATTAGGCCAAGCTTGTTCACCAGGGTGTCGCCCTGGGCAGGATCCCCGGGTACCG |
| oligo-006 | Cy5-labeled dsDNA cleavage, cleavage sites determination and *trans*-cleavage assay of Casπ | Non-labelled complementary strand containing a TGCCC PAM (blue) and a protospacer region (red) of oligo-004 and oligo-005. | OH-CGGTACCCGGGGATCCTGCCCAGGGCGACACCCTGGTGAACAAGCTTGGCCTAATCATGGTCATA |
| Oligo-007 | Cy5-labeled dsDNA cleavage and *trans*-cleavage assay of LbCas12a | Reverse PCR primer for cy5-labeled dsDNA target amplification and non-labelled DNA target strand for Cas12a *trans*-cleavage assay of LbCas12a. | HO-TATGACCATGATTAGGCCAAGCTTGTTCACCAGGGTGTCGCCCTGAAAAGGATCCCCGGGTACCG |
| Oligo-008 | *Trans*-cleavage assay of LbCas12a | Non-labelled DNA non-target strand containing a TTTTC PAM (blue) and a protospacer region (red) for Cas12a *trans*-cleavage assay. | HO-CGGTACCCGGGGATCCTTTTCAGGGCGACACCCTGGTGAACAAGCTTGGCCTAATCATGGTCATA |
| Oligo-010 | *Trans*-cleavage assay | ssDNA substrate. | $NH_2$-CTGAAAACTCGCAGCTACCTTAACGGCCGCCCATTCCTATGCGTACATCTGGATCAAGCC |
| Oligo-011 | Casπ R-loop ternary complex reconstitution | DNA non-target strand containing a TTTTC PAM (blue) and a protospacer region (red) for Casπ R-loop complex reconstitution. | HO-CGGGATGCCCAGGGCGACACAAGTTGTCCAGATCC |
| Oligo- | Casπ R-loop ternary | DNA target strand for Casπ R- | HO- |

| | | | |
|---|---|---|---|
| 012 | complex reconstitution | loop complex reconstitution. | GGATCGTTCACCAGGGTGTCGCC CTGGGCATCCCG |
| Oligo-013 | Cy5-labeled dsDNA mismatch cleavage assay | Reverse PCR primes for cy5-labeled dsDNA target amplification and mismatch cleavage assay with mismatch at position 1 (brown). | HO-TATGACCATGATTAGGCCAAG CTTGTTCACCAGGGTGTCGCC CGGGGCAGGATCCCCGGGTA CCG |
| Oligo-014 | Cy5-labeled dsDNA mismatch cleavage assay | Reverse PCR primes for cy5-labeled dsDNA target amplification and mismatch cleavage assay with mismatch at position 2 (brown) | HO-TATGACCATGATTAGGCCAAG CTTGTTCACCAGGGTGTCGCC ATGGGCAGGATCCCCGGGTA CCG |
| Oligo-015 | Cy5-labeled dsDNA mismatch cleavage assay | Reverse PCR primes for cy5-labeled dsDNA target amplification and mismatch cleavage assay with mismatch at position 3 (brown) | HO-TATGACCATGATTAGGCCAAG CTTGTTCACCAGGGTGTCGCA CTGGGCAGGATCCCCGGGTAC CG |
| Oligo-016 | Cy5-labeled dsDNA mismatch cleavage assay | Reverse PCR primes for cy5-labeled dsDNA target amplification and mismatch cleavage assay with mismatch at position 4 (brown) | HO-TATGACCATGATTAGGCCAAG CTTGTTCACCAGGGTGTCGAC CTGGGCAGGATCCCCGGGTAC CG |
| Oligo-017 | Cy5-labeled dsDNA mismatch cleavage assay | Reverse PCR primes for cy5-labeled dsDNA target amplification and mismatch cleavage assay with mismatch at position 5 (brown) | HO-TATGACCATGATTAGGCCAAG CTTGTTCACCAGGGTGTCTCC CTGGGCAGGATCCCCGGGTAC CG |
| Oligo-018 | Cy5-labeled dsDNA mismatch cleavage assay | Reverse PCR primes for cy5-labeled dsDNA target amplification and mismatch cleavage assay with mismatch at position 6 (brown) | HO-TATGACCATGATTAGGCCAAG CTTGTTCACCAGGGTGTAGCC CTGGGCAGGATCCCCGGGTAC CG |
| Oligo-019 | Cy5-labeled dsDNA mismatch cleavage assay | Reverse PCR primes for cy5-labeled dsDNA target amplification and mismatch cleavage assay with mismatch at position 7 (brown) | HO-TATGACCATGATTAGGCCAAG CTTGTTCACCAGGGTGGCGCC CTGGGCAGGATCCCCGGGTAC CG |
| Oligo-020 | Cy5-labeled dsDNA mismatch cleavage assay | Reverse PCR primes for cy5-labeled dsDNA target amplification and mismatch cleavage assay with mismatch at position 8 (brown) | HO-TATGACCATGATTAGGCCAAG CTTGTTCACCAGGGTTTCGCC CTGGGCAGGATCCCCGGGTAC CG |
| Oligo- | Cy5-labeled | Reverse PCR primes for cy5- | HO- |

| | | | |
|---|---|---|---|
| 021 | dsDNA mismatch cleavage assay | labeled dsDNA target amplification and mismatch cleavage assay with mismatch at position 9 (brown) | TATGACCATGATTAGGCCAAG CTTGTTCACCAGGGGGTCGCC CTGGGCAGGATCCCCGGGTAC CG |
| Oligo-022 | Cy5-labeled dsDNA mismatch cleavage assay | Reverse PCR primes for cy5-labeled dsDNA target amplification and mismatch cleavage assay with mismatch at position 10 (brown) | HO-TATGACCATGATTAGGCCAAG CTTGTTCACCAGGTTGTCGCC CTGGGCAGGATCCCCGGGTAC CG |
| Oligo-023 | Cy5-labeled dsDNA mismatch cleavage assay | Reverse PCR primes for cy5-labeled dsDNA target amplification and mismatch cleavage assay with mismatch at position 11 (brown) | HO-TATGACCATGATTAGGCCAAG CTTGTTCACCAGTGTGTCGCC CTGGGCAGGATCCCCGGGTAC CG |
| Oligo-024 | Cy5-labeled dsDNA mismatch cleavage assay | Reverse PCR primes for cy5-labeled dsDNA target amplification and mismatch cleavage assay with mismatch at position 12 (brown) | HO-TATGACCATGATTAGGCCAAG CTTGTTCACCATGGTGTCGCC CTGGGCAGGATCCCCGGGTAC CG |
| Oligo-025 | Cy5-labeled dsDNA mismatch cleavage assay | Reverse PCR primes for cy5-labeled dsDNA target amplification and mismatch cleavage assay with mismatch at position 13 (brown) | HO-TATGACCATGATTAGGCCAAG CTTGTTCACCCGGGTGTCGCC CTGGGCAGGATCCCCGGGTAC CG |
| Oligo-026 | Cy5-labeled dsDNA mismatch cleavage assay | Reverse PCR primes for cy5-labeled dsDNA target amplification and mismatch cleavage assay with mismatch at position 14 (brown) | HO-TATGACCATGATTAGGCCAAG CTTGTTCACAAGGGTGTCGCC CTGGGCAGGATCCCCGGGTAC CG |
| Oligo-027 | Cy5-labeled dsDNA mismatch cleavage assay | Reverse PCR primes for cy5-labeled dsDNA target amplification and mismatch cleavage assay with mismatch at position 15 (brown) | HO-TATGACCATGATTAGGCCAAG CTTGTTCAACAGGGTGTCGCC CTGGGCAGGATCCCCGGGTAC CG |
| Oligo-028 | Cy5-labeled dsDNA mismatch cleavage assay | Reverse PCR primes for cy5-labeled dsDNA target amplification and mismatch cleavage assay with mismatch at position 16 (brown) | HO-TATGACCATGATTAGGCCAAG CTTGTTCCCCAGGGTGTCGCC CTGGGCAGGATCCCCGGGTAC CG |
| Oligo-029 | Cy5-labeled dsDNA mismatch cleavage assay | Reverse PCR primes for cy5-labeled dsDNA target amplification and mismatch cleavage assay with mismatch at position 17 (brown) | HO-TATGACCATGATTAGGCCAAG CTTGTTAACCAGGGTGTCGCC CTGGGCAGGATCCCCGGGTAC CG |

| | | | |
|---|---|---|---|
| Oligo-030 | Cy5-labeled dsDNA mismatch cleavage assay | Reverse PCR primes for cy5-labeled dsDNA target amplification and mismatch cleavage assay with mismatch at position 18 (brown) | HO-TATGACCATGATTAGGCCAAGCTTGTGCACCAGGGTGTCGCCCTGGGCAGGATCCCCGGGTACCG |
| Oligo-031 | Cy5-labeled dsDNA mismatch cleavage assay | Reverse PCR primes for cy5-labeled dsDNA target amplification and mismatch cleavage assay with mismatch at position 19 (brown) | HO-TATGACCATGATTAGGCCAAGCTTGGTCACCAGGGTGTCGCCCTGGGCAGGATCCCCGGGTACCG |
| Oligo-032 | Cy5-labeled dsDNA mismatch cleavage assay | Reverse PCR primes for cy5-labeled dsDNA target amplification and mismatch cleavage assay with mismatch at position 20 (brown) | HO-TATGACCATGATTAGGCCAAGCTTCTTCACCAGGGTGTCGCCCTGGGCAGGATCCCCGGGTACCG |
| RNA-001 | *In vitro* plasmids cleavage | crRNA containing target spacer (red) of Casπ-1 | ACACUCUAAAGGAAUGAAAGAGGGCGACACCCUGGUGAAC |
| RNA - 002 | *In vitro* plasmids cleavage | crRNA containing target spacer (red) of Casπ-2 | AACGCUCUUAGGGAAUGAAAGAGGGCGACACCCUGGUGAAC |
| RNA - 003 | *In vitro* plasmids cleavage | tracrRNA sequence of Casπ-1 | GUCUGCCGAAGACGCCGCACGGAGCCUGGGCCGGAAUCGUAGAUCGAACGCGGCAUCGAAGCCCUGCAGCCCUUCGGGGCCAAGGCGGCGCAGCAAGCCUCUUUCAGGCGGCAGAGUCCUUUAGAGUGU |
| RNA - 004 | *In vitro* plasmids cleavage | tracrRNA sequence of Casπ-2 | GUCUCGACUAUGCCGUACCACUAGACCGAGCCUACACGGCACGCGGUCAUAGCGUUAACCAAGGCGUGGUGACAAGCCUCUUUCAGGCGUCGGACACUUAAGAGCGUU |
| RNA - 005 | *In vitro* plasmids cleavage, cy5-labeled dsDNA cleavage, cleavage sites determination, *trans*-cleavage and complex reconstitution | sgRNA joint by tracrRNA and crRNA, which contained target spacer (red), with GAGAG loop (green) of Casπ-1 | GUCUGCCGAAGACGCCGCACGGAGCCUGGGCCGGAAUCGUAGAUCGAACGCGGCAUCGAAGCCCUGCAGCCCUUCGGGGCCAAGGCGGCGCAGCAAGCCUCUUUCAGGCGGCAGAGUCCUUUAGAGUGUGAGAGACACUCUAAAGGAAUGAAAGAGGGCGACACCCUGGUGAAC |

| | | | |
|---|---|---|---|
| RNA - 006 | *In vitro* plasmids cleavage, cy5-labeled dsDNA cleavage, cleavage sites determination, *trans*-cleavage and complex reconstitution | sgRNA joint by tracrRNA and crRNA, which contained target spacer (red), with GAGAG loop (green) of Casπ-2 | GUCUCGACUAUGCCGUACCACUAGACCGAGCCUACACGGCACGCGGUCAUAGCGUUAACCAAGGCGUGGUGACAAGCCUCUUUCAGGCGUCGGACACUUAAGAGCGUUGAGAGAACGCUCUUAGGGAAUGAAAGAGGGCGACACCCUGGUGAAC |
| RNA - 007 | *In vitro* plasmids cleavage, cy5-labeled dsDNA cleavage, 12nt truncated A-II region | sgRNA joint by tracrRNA and crRNA, which contained target spacer (red), with GAGAG loop (green) and 12nt truncated A-II region of Casπ-1 | GUCUGCCGAAGACGCCGCACGGAGCCUGGGCCGGAAUCGUAGAUCGAACGCGGCAUCGAAGCCCUGCAGCCCUUCGGGGCCAAGGCGGCGCAGCAAGCCUCUUUCAGGCGGCAGAGUCCUUUAGAGAGUAAAGGAAUGAAAGAGGGCGACACCCUGGUGAAC |
| RNA - 008 | *In vitro* plasmids cleavage, cy5-labeled dsDNA cleavage, 24nt truncated A-II region | sgRNA joint by tracrRNA and crRNA, which contained target spacer (red), with GAGAG loop (green) and 24nt truncated A-II region of Casπ-1 | GUCUGCCGAAGACGCCGCACGGAGCCUGGGCCGGAAUCGUAGAUCGAACGCGGCAUCGAAGCCCUGCAGCCCUUCGGGGCCAAGGCGGCGCAGCAAGCCUCUUUCAGGCGGCAGAGUGAGAGAAUGAAAGAGGGCGACACCCUGGUGAAC |
| RNA - 009 | *In vitro* cy5-labeled dsDNA cleavage, and *trans*-cleavage | sgRNA of LbCas12a, which contained target spacer (red) | GAGAUUUCUACUCUUGUAGAUAGGGCGACACCCUGGUGAAC |
| π-sg1 | Cell genome editing | *EGFP* gene target 1 of Casπ, PAM of the targets: CCT | CTGCTCAGGTGGAGATGAAC |
| π-sg2 | Cell genome editing | *EGFP* gene target 2 of Casπ, PAM of the targets: CCC | GCTTCTTGTTCATCTCCACC |
| π-sg3 | Cell genome editing | *EGFP* gene target 3 of Casπ, PAM of the targets: CCC | GAGCTCAGCCACACTGTCCG |
| π-sg4 | Cell genome editing | *EGFP* gene target 4 of Casπ, PAM of the targets: CCC | CGAGCTCAGCCACACTGTCC |
| π-sg5 | Cell genome editing | *EGFP* gene target 5 of Casπ, PAM of the targets: CCT | TCTCCAGCTTCTGCTTCACC |
| π-dest | Cell genome editing | *EGFP* gene non-target of Casπ, | GGAAGAGCGAGCTCTTCC |
| 12a-sg1 | Cell genome editing | *EGFP* gene target 1 of Cas12a, PAM of the targets: TTTC | CCGCTTCTTGTTCATCTCCA |
| 12a-sg2 | Cell genome editing | *EGFP* gene target 2 of Cas12a, PAM of the targets: TTTC | TGAAACTCAGTTTCCCGCTT |
| 12a-sg3 | Cell genome editing | *EGFP* gene target 3 of Cas12a, PAM of the targets: CTTC | TTCCGAAGAGCGGCTGCTGT |
| 12a-sg4 | Cell genome editing | *EGFP* gene target 4 of Cas12a, PAM of the targets: CTTC | GGAAGAAGCACGCGGACAGT |
| 12a-sg5 | Cell genome editing | *EGFP* gene target 5 of Cas12a, PAM of the targets: CTTC | TCCTTCTCCAGCTTCTGCTT |

| | | | |
|---|---|---|---|
| 12a-dest | Cell genome editing | *EGFP* gene non-target of Cas12a | GGAAGAGCGAGCTCTTCC |
| Cas9-sg1 | Cell genome editing | *EGFP* gene target 1 of of SpyCas9, PAM of the targets: AGG | GTTCATCTCCACCTGAGCAG |
| Cas9-sg2 | Cell genome editing | *EGFP* gene target 2 of of SpyCas9, PAM of the targets: GGG | GGTGGAGATGAACAAGAAGC |
| Cas9-sg3 | Cell genome editing | *EGFP* gene target 3 of of SpyCas9, PAM of the targets: GGG | CGGACAGTGTGGCTGAGCTC |
| Cas9-sg4 | Cell genome editing | *EGFP* gene target 4 of of SpyCas9, PAM of the targets: GGG | GGACAGTGTGGCTGAGCTCG |
| Cas9-sg5 | Cell genome editing | *EGFP* gene target 5 of of SpyCas9, PAM of the targets: AGG | GGTGAAGCAGAAGCTGGAGA |
| Cas9-dest | Cell genome editing | *EGFP* gene non-target of SpyCas9 | GGAAGAGCGAGCTCTTCC |
| ccdB-T | Bacterial plasmids interference | *ccdB* gene target of Casπ | CGATAACGGAGACCGGCACA |
| CcdB-dest | Bacterial plasmids interference | *ccdB* gene non-target of Casπ | GGAAGAGCGAGCTCTTCC |
| *B2M*-1 | Cell genome editing | *B2M* gene target 1 of Casπ, PAM of the targets: CCC | GATATTCCTCAGGTACTCCA |
| *B2M*-2 | Cell genome editing | *B2M* gene target 2 of Casπ, PAM of the targets: CCT | GAATCTTTGGAGTACCTGAG |
| *TP53*-1 | Cell genome editing | *TP53* gene target 1 of Casπ, PAM of the targets: CCC | ACCATGAGCGCTGCTCAGAT |

**Table. S4. Statistics for cryo-EM analysis and model refinement.**

| | Casπ |
|---|---|
| PDB ID | 7YOJ |
| EMDB ID | EMD-33983 |
| **Data collection and processing** | |
| Microscope | Titan Krios |
| Detector | Gatan K3 with GIF Quantum (20eV slit) |
| CS (mm) | 0.01 |
| Magnification | 81K |
| Pixel size (Å) | 0.856 |
| Electron dose ($e^-$/ $Å^2$) | 50(32 frames) |
| Defocus range (μm) | -1.5 ~ -2.0 |
| Micrographs collected | 6,690 |
| Micrographs used | 6,690 |
| **Reconstruction** | |
| Software | RELION-3.1, cryoSPARC-3.3.1 |
| Particles picked | 2,544,599 |
| Particles refinement | 39,500 |
| Symmetry | C1 |
| Resolution (Å) | 3.4 |
| Sharpening B-factor ($Å^2$) | 111.3 |
| **Refinement** | |
| Software | PHENIX-1.20.1 |
| Model composition | |
| Number of atoms | 11,545 |
| Protein residues | 867 |
| Nucleotides | 216 |
| Ligand | 0/0 |
| B factors (Protein/Nucleotide/Ligand) | 113.04/191.67/0 |
| Bonds RMSD | |
| Bonds lengths (Å) | 0.004 |
| Bonds angles (°) | 0.681 |
| **Validation** | |
| MolProbity score | 2.02 |
| Clash score | 8 |

| | |
|---|---|
| Rotamer outliers (%) | 0.27 |
| C-beta outliers (%) | 0.00 |
| Ramachandran plot | |
| Favored (%) | 94% |
| Allowed (%) | 6% |
| Outlier (%) | 0 |
| Model vs. Data | |
| CC mask/box | 0.82/0.78 |

**Titles and Legends for Supplementary Video**

**Video. S1. The cryo-EM reconstruction and atomic model of Casπ R-loop complex.**

The protein domains, RNA elements and DNA target are presented and colored referring to Figures. 6 and 8.
